# Incipient resistance to an effective pesticide results from genetic adaptation and the canalization of gene expression

**DOI:** 10.1101/2019.12.19.882860

**Authors:** Xiaoshen Yin, Alexander S. Martinez, Abigail Perkins, Morgan M. Sparks, Avril M. Harder, Janna R. Willoughby, Maria S. Sepúlveda, Mark R. Christie

**Affiliations:** Department of Biological Sciences, Purdue University; 915 W. State St., West Lafayette, Indiana 47907-2054 USA; Department of Anatomy, Cell Biology & Physiology, Indiana University School of Medicine; 320 W. 15^th^ St., Indianapolis, Indiana 46202-2266 USA; School of Forestry and Wildlife Sciences, Auburn University; 602 Duncan Dr., Auburn, Alabama 36849 USA; Department of Forestry and Natural Resources, Purdue University; 715 W. State St., West Lafayette, Indiana 47907-2054 USA

**Author notes:** Authors contributed equally (Xiaoshen Yin, Alexander S. Martinez). To whom correspondence should be addressed: Mark Christie, 915 W. State Street, Department of Biological Sciences, Purdue University, West Lafayette, IN 47907-2054.

**Keywords:** ATP synthase, contemporary evolution, genetic adaptation, resistance, pesticides, RNA-seq

## Abstract

The resistance of bacteria, disease vectors, and pest species to chemical controls has vast ecological, economic, and societal costs. In most cases, resistance is only detected after spreading throughout an entire population. Detecting resistance in its incipient stages, by comparison, provides time to implement preventative strategies. Incipient resistance can be detected by coupling standard toxicology assays with large-scale gene expression experiments. We apply this approach to a system where an invasive parasite, sea lamprey (*Petromyzon marinus*), has been treated with the highly-effective pesticide 3-trifluoromethyl-4-nitrophenol (TFM) for 60 years. Toxicological experiments revealed that lamprey from treated populations did not have higher survival to TFM exposure than lamprey from their native range, demonstrating that full-fledged resistance has not yet evolved. In contrast, we find hundreds of genes differentially expressed in response to TFM in the population with the longest history of exposure, many of which relate to TFM’s primary mode of action, the uncoupling of oxidative phosphorylation and subsequent depletion of ATP. Three genes critical to oxidative phosphorylation, *ATP5PB, PLCB1*, and *NDUFA9*, were nearly fixed for alternative alleles in comparisons of SNPs between native and treated populations (*F*_*ST*_ > 5 SD from the mean). *ATP5PB* encodes subunit b of ATP synthase and an additional subunit, *ATP5F1B*, was canalized for high expression in treated populations, but remained plastic in response to TFM treatment in individuals from the native range. These combined genomic and transcriptomic results demonstrate that an adaptive, genetic response to TFM is driving incipient resistance in a damaging pest species.

## Introduction

The evolution of resistance often occurs in response to the continued and large-scale application of chemical compounds such as antibiotics, herbicides, and pesticides (Davies & Davies, 2010; Délye, Jasieniuk, & Le Corre, 2013; Georghiou & Taylor, 1977; Gould, Brown, & Kuzma, 2018). The costs associated with the evolution of resistance measure in the billions of dollars per year and cut across diverse fields ranging from healthcare to agriculture (Levy & Marshall, 2004; Pimentel & Burgess, 2014; Smith & Coast, 2013). Increased rates of disease, decreased food security, and large-scale environmental costs represent only some of the negative effects when microbes, pathogens, and pests can no longer be effectively controlled (Gould et al., 2018; Ranson & Lissenden, 2016). In light of these costs, there is a considerable need to prevent or delay the evolution of resistance. Unfortunately, resistance is often detected too late, after most individuals in a population have acquired a resistant phenotype (McKenna, 2013). Once this stage is reached, there is little to be done except to implement alternative control measures, which are often less effective (Ghosh, Sarkar, Issa, & Haldar, 2019; Peterson, Collavo, Ovejero, Shivrain, & Walsh, 2018), or stop employing control measures until resistant genotypes are eliminated from the population, which can take tens to hundreds of generations (Andersson & Hughes, 2010; Andersson & Levin, 1999; Christie, Sepúlveda, & Dunlop, 2019).

One solution to mitigating the costs of resistance is early detection; identifying the evolution of resistance in its incipient stages can allow time for developing alternative control measures or implementing management actions that can prevent the onset of population-wide resistance (Andow & Ives, 2002; Apple & Smith, 1976; Stenberg, 2017). One approach to detecting incipient resistance is to screen a large fraction of the population for resistant genotypes or phenotypes. However, such measures can be costly and impractical because of the large sample sizes required to identify small numbers of fully resistant individuals. Detectability issues (e.g., false positives and negatives) compound this problem. Alternatively, the coupling of experimental, eco-toxicological assays with the identification of genomic and transcriptomic responses in populations with varied treatment histories can provide key insights into how and whether resistance is beginning to evolve. We applied this design to an invasive vertebrate pest species that has been controlled with a chemical pesticide for 60 years.

Throughout the Laurentian Great Lakes, invasive sea lamprey (*Petromyzon marinus*) have negatively impacted native fish communities by feeding parasitically on other fishes (Lawrie, 1970). With the construction and improvement of shipping canals in the early 1900s, sea lamprey expanded their range from the northwestern Atlantic Ocean to the Great Lakes (Smith & Tibbles, 1980) (Figure 1). Adult sea lamprey are hematophagous parasites of other fishes and are not host specific (Figure 1d). The lack of host specialization, coupled with ample spawning habitat found throughout the Great Lakes region, spurred the growth of parasitic sea lamprey populations into the millions of individuals (Smith & Tibbles, 1980). High abundance of invasive sea lamprey in the Great Lakes contributed to rapid and substantial declines in native lake trout (*Salvelinus namaycush*) and other ecologically and commercially important fishes (Baldwin, Saalfeld, Dochoda, Buettner, & Eshenroder, 2002; Bronte et al., 2003). In response to the effects of sea lamprey on native fish and fisheries, there was an immediate and concerted effort to develop efficient means for controlling sea lamprey populations in the Great Lakes.

**Figure 1.**
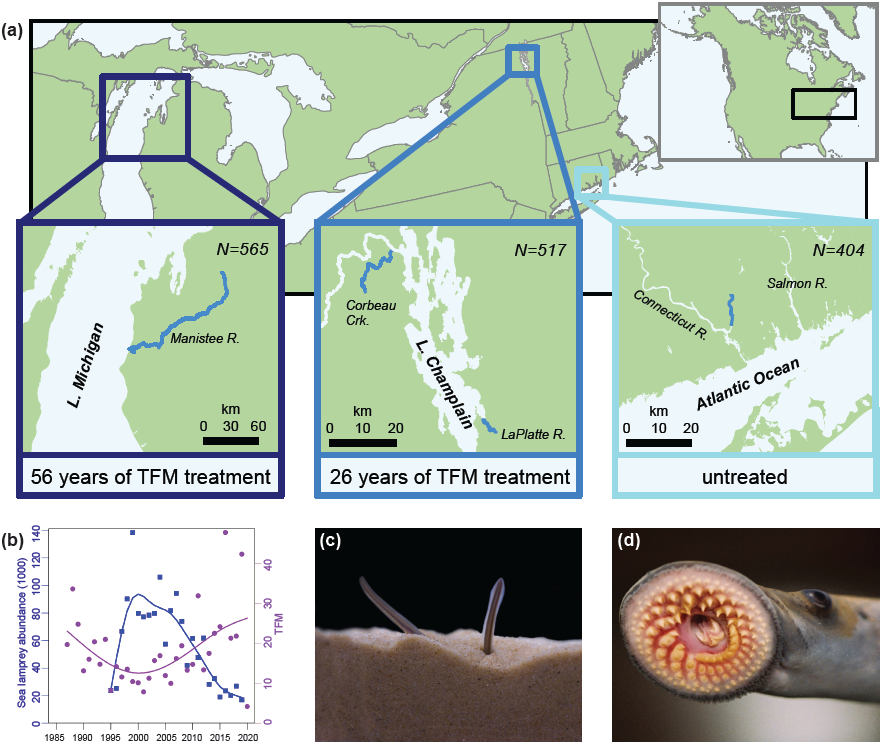
Sample collection sites, sample sizes, and pesticide (TFM) treatment histories for larval sea lamprey used in this study (a). Connecticut River sea lamprey were collected from their native range whereas sea lamprey from Lake Michigan and Lake Champlain are considered invasive. Sea lamprey abundance and metric tons of TFM applied to Lake Michigan from 1987 onwards (b); notice that TFM is an effective pesticide with decrease in lamprey abundance following increases in TFM application. The larval stage of sea lamprey remain buried in the sediment (c), aggregate in high densities, and are targeted with the pesticide TFM. After transformation, the juveniles are generalist parasites, using their circularly-arrayed teeth to attach to hosts and their rasping tongues to feed (d).

One effort, initiated in the 1950s, involved testing over 4000 chemical compounds on sea lamprey and other fish species (Applegate, 1957). The organic compound 3-trifluoromethyl-4-nitrophenol (hereafter, TFM) was found to effectively kill larval sea lamprey, but had no detectable effects on other fishes at low concentrations (Hubert, 2003; Middaugh, Sepúlveda, & Höök, 2014). The larval stage of sea lamprey (Figure 1c) are targeted because they are small (typically < 120 mm) and aggregate at high densities. The primary mode of action for TFM is to uncouple oxidative phosphorylation (Birceanu, McClelland, Wang, Brown, & Wilkie, 2011; Birceanu, McClelland, Wang, & Wilkie, 2009), and because sea lamprey are highly sensitive to this mode of action, TFM is a very effective pesticide (Figure 1b). Since the application of TFM to locations with abundant larval lamprey, invasive sea lamprey populations have declined by up to 90% (Heinrich et al., 2003; Smith & Tibbles, 1980). However, TFM applications kill most, but not all, larval sea lamprey (Dunlop et al., 2017) and it is possible that resistant individuals could survive and reproduce. If this process is repeated over enough generations, then resistant individuals could increase in frequency – a scenario documented in many systems where pests have been controlled by chemical means (Whalon, Mota-Sanchez, & Hollingworth, 2008). The evolution of resistance would greatly reduce the effectiveness of TFM, which is currently the primary control method for invasive sea lamprey (Dunlop et al., 2017), and decreased pesticide effectiveness would likely hamper the restoration of native and commercially-important fish populations throughout the Great Lakes.

To test for the evolution of resistance, we first collected larval sea lamprey from three populations with varied histories of TFM treatment (Figure 1a): Lake Michigan (56 years of TFM treatment), Lake Champlain (26 years of TFM treatment), and Connecticut River (native range; 0 years of TFM treatment). Larvae were acclimated in a common environment for 4 months before being experimentally exposed to TFM at lethal, sublethal, and control (no TFM) levels. Detailed survival and toxicological analyses were coupled with tissue-specific RNA sequencing (RNA-seq). Survival analyses revealed that the outright resistance of sea lamprey to lethal concentrations of TFM could not be detected, but genomic and transcriptomic analyses suggest that the early, incipient stages of resistance are well under way in populations with a history of pesticide exposure.

## Materials and Methods

### Sample collection and experimental design

We collected a total of 2,035 larval sea lamprey (ammocoetes) from five locations during summers of 2016 and 2017 using pulsed-DC backpack electrofishers, focusing our sampling efforts on populations with different durations of exposure to TFM (Figure 1; Table S1). In 2016, we obtained 565 ammocoetes from the Manistee River in Lake Michigan, 517 ammocoetes from Corbeau Creek and the LaPlatte River in Lake Champlain, and 404 ammocoetes from Connecticut River (Figure 1). Lake Michigan ammocoetes have been treated with TFM since 1960 (Lavis, Henson, Johnson, Koon, & Ollila, 2003), Lake Champlain ammocoetes have been treated with TFM since 1990 (Marsden et al., 2003), and Connecticut River is part of the sea lamprey’s native range and has never been treated with TFM. To examine lamprey of similar ages and developmental stages, we only collected ammocoetes from a narrow size range (80-120 mm). All ammocoetes were collected within five calendar days (July 11-July 15, 2016) of each other and were transported to Purdue’s Aquaculture Research Lab immediately after collection. Upon arrival at Purdue’s Aquaculture Research Lab, ammocoetes were acclimated into 24, 30-gallon flow-through holding tanks (see *SI Materials and Methods* for details). In 2017, we collected an additional 284 ammocoetes from the Platte River in Lake Michigan and 265 ammocoetes from Turner Falls, Massachusetts, to independently validate our 2016 toxicological analyses (Table S1, Table S2).

### Lethal exposure trials

After acclimation, we constructed an experimental array to conduct exposure trials in a temperature-controlled environmental chamber (12.8°C) consisting of 36, 2.5-gallon glass aquaria (see *SI Materials and Methods* for details). With ammocoetes collected in 2016 (2016 trials; Table S2), we conducted two toxicological exposure trials to directly assess whether sea lamprey populations exhibited differential mortality in response to TFM exposure. We began each exposure by transferring ammocoetes from their holding tanks into randomized experimental array tanks pre-filled with 7 L of water. For the first 2016 trial (TOX 1; Table S2), each population was allotted 12 aquaria with seven ammocoetes per tank (n = 84 lamprey per population). Within a population, eight treatment aquaria received TFM (n = 56 individuals per population) while four control aquaria (n = 28 individuals per population) received a volume of water equal to the volume of TFM distributed into treatment aquaria. In TOX 1, we aimed for high (i.e., > 80%) mortality to allow us to assess whether survivorship varied among populations and we challenged ammocoetes in treatment tanks with a TFM concentration of 3 mg/L, as determined by a multi-concentration, small-scale pilot exposure trial. We allowed ammocoetes to acclimate for a period of one hour to the experimental tanks before introducing TFM with peristaltic pumps. To simulate conditions associated with a typical TFM stream application, ammocoetes were exposed to TFM for ten hours followed by a two-hour ‘wash-out period’. TFM concentrations were monitored hourly and verified by measuring TFM absorption using a spectrophotometer. Mortality was assessed hourly over the 12-hour duration of the experiment. At the conclusion of the experiment, dead individuals were measured for weight and length. The second 2016 trial (TOX 2; Table S2) had the same experimental setup as TOX 1, except that a TFM concentration of 1.67 mg/L was administered to treatment tanks to target more modest levels of mortality (Table S2). To analyze survivorship data, we conducted a Kaplan-Meier survival analysis (Kaplan & Meier, 1958) that generates an estimate of survival probability (*S*) at time t_i_ and is calculated as:

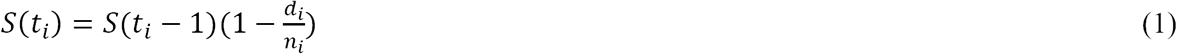

where *S*(*t*_*i*_ – 1)is the probability of being alive at time *t*_*i*_ – 1, *n*_*i*_ is the number of individuals alive just prior to *t*_*i*_, and *d*_*i*_ is the number of deaths occurring at *t*_*i*_ (Kaplan & Meier, 1958). We performed the survival analysis in R (R core team, 2019), using both the SURVIVAL and OIsurv packages to create survival curves and assess statistical differences between the survival curves of three populations (Therneau, 2015). *P*-values associated with pairwise tests used to determine which populations differ in their survivorship were FDR adjusted using the Benjamini-Hochberg method (Benjamini & Hochberg, 1995).

To independently confirm toxicological analyses from 2016 trials, we conducted three additional toxicological exposure trials with ammocoetes collected in 2017 (2017 trials; Table S2). After a six-week acclimation, we conducted toxicological exposure trials similarly to the 2016 trials with three modifications to increase power for the detection of lethal concentrations in a given population. First, in the 2017 trials, each trial examined only one population at a time. Second, trials lasted for 20 hours instead of 12 hours. Third, these trials consisted of five treatments—each with three replicate tanks of seven individuals—including the control (0, 0.5, 1.2, 1.5, and 2 mg/L of TFM), instead of a single TFM treatment (Table S2). With this experimental design, we could quantify lethal concentrations (e.g., LC50). To compare differences in survival, we computed lethal concentrations for each trial using the DRC package (v 3.0-1) (Ritz, Baty, Streibig, & Gerhard, 2015), and made comparisons among trials at different dosage levels using Kaplan-Meier survival analyses in the SURVIVAL package (Therneau, 2015) with visualization using the SURVMINER package (Kassambara & Kosinski, 2018).

### Sublethal exposure trials

To assess whether resistance was evolving at a sublethal level, we conducted two additional exposure trials (i.e., gene expression trials GE 1 and GE 2; Table S2) to assess (i) how gene expression profiles changed in response to TFM, and (ii) whether TFM exposure altered patterns of gene expression differently among three populations with varying histories of exposure. The experimental design for sublethal exposure was similar to that for the toxicological exposure trials outlined above with two key differences. First, we exposed ammocoetes to sublethal levels of TFM (i.e., 0.3 mg/L in GE 1 and 0.2 mg/L in GE 2; Table S2). Second, we preserved tissue from ammocoetes at a predetermined time point (t = hour 6) during the 12-hour gene expression exposure trial for RNA-seq. All populations were run concurrently, and all individuals were sampled at the same time such that any differences in gene expression could not be due to differences in time of day, week, or month. Sampling consisted of removing a single ammocoete per replicate tank and immediately euthanizing it in a lethal dose of MS-222 before weighing, measuring, and flash-freezing in cryovials with liquid nitrogen. At the completion of the experiment, cryovials were directly transferred to a −80°C freezer.

### RNA-seq and differential gene expression analysis

We used tissue-specific RNA-seq to compare patterns of gene expression in ammocoetes from the three populations with varying histories of exposure to TFM. Comparing tissue-specific patterns of expression required that we first dissect specific tissue samples from the body segments collected during gene expression trials. Given TFM’s established mode of action, we targeted ammocoete muscle, liver, and brain tissues for downstream analyses of gene expression profiles. We first transferred individuals frozen at −80°C into a solution of RNAlater-ICE (Ambion Inc., Austin, TX, USA) at 10 volumes of solution relative to sample mass and allowed the solution to permeate into the tissue for 16 hours at −20°C. Once the solution had permeated into the tissue and segments had thawed, we dissected muscle, liver, and brain tissues for subsequent mRNA extractions (Table S3). In total, we performed 71 mRNA extractions across three populations (Lake Michigan, Lake Champlain, and Connecticut River), and three tissue types (muscle, liver, and brain) using the Qiagen RNeasy Mini Kit (Table S3). Library preparation and mRNA sequencing were performed by the Purdue Genomics Core Facility with samples from GE 1 and GE 2 (Table S2) being sequenced on either an Illumina 2500 or an S2 NovaSeq lane. We obtained a total of 3.25 billion 150 bp paired-end reads, which we first used to create an annotated transcriptome (see *SI Materials and Methods* for details on transcriptome assembly).

We aligned all reads to our assembled sea lamprey transcriptome and estimated transcript abundances using RSEM (Li & Dewey, 2011), and identified differentially expressed genes (hereafter, DEGs) using a GLM with TFM concentration and sequencing machine as fixed effects in edgeR (McCarthy, Chen, & Smyth, 2012; Robinson, McCarthy, & Smyth, 2010). We compared gene expression profiles of treated versus untreated ammocoetes within each population to identify genes that were significantly up- and down-regulated in response to TFM treatment and were specific to each population. For genes to be classified as differentially expressed, we set minimum thresholds for significance (FDR correction via the Benjamini-Hochberg method; *p*-value < 0.05). While we did not explicitly filter DEGs by log fold change, the minimum log2-fold changes we observed for DEGs were > 2 across all three populations (2.04, 3.50, and 2.74 for Lake Michigan, Lake Champlain, and Connecticut River populations, respectively). Finally, GOseq was used to determine which gene ontology functional categories were overrepresented among differentially expressed genes in TFM treated individuals for each population-tissue combination (Young, Wakefield, Smyth, & Oshlack, 2010). Only functionally enriched categories significant below an FDR-adjusted *p*-value threshold of 0.05 were retained. We constructed gene ontology (GO) hierarchy networks of the top 100 significant GO terms identified with DEGs (FDR-corrected *p*-value < 0.01) using the metacoder package (Foster, Sharpton, & Grünwald, 2017).

### Genomic analysis

Using the trimmed RNA-seq reads, we called SNPs following the variant discovery pipeline provided by the Genome Analysis Toolkit (GATK) (McKenna et al., 2010). We called SNPs using both RNA-seq and DNA-seq variant calling pipelines (Brouard, Schenkel, Marete, & Bissonnette, 2019) for muscle (n=44) and liver (n=19) tissue samples separately (see *SI Materials and Methods* for details). After comparing variants from both pipelines, we only included SNPs that were identified in both the DNA-seq and RNA-seq pipelines. We used the genotypes from the DNA-seq pipeline because it calls genotypes that are homozygous for the reference allele, while the RNA-seq pipeline cannot. For each individual, genotypes with fewer than five reads were set as missing values. Muscle samples from two individuals 401 and 402 (Table S3) had over 90% missing data and thus were excluded from further analyses. Finally, loci genotyped across at least 80% of all samples were retained as high-quality SNPs, and this resulted in SNPs called for 42 and 19 samples from muscle and liver tissues, respectively (Table S3). Separate analyses were conducted on high-quality SNPs obtained from different tissue samples (i.e., muscle vs. liver). In each population, we further excluded loci out of Hardy-Weinberg equilibrium, which we identified using the package HWxtest (likelihood ratio *p*-value < 0.05) (Engels, 2009). We calculated pairwise *F*_*ST*_ for each locus among all three populations (i.e., Lake Michigan vs. Lake Champlain, Lake Michigan vs. Connecticut River, and Lake Champlain vs. Connecticut River populations) using Weir and Cockerham’s unbiased estimator (Weir & Cockerham, 1984). To detect outliers, we z-transformed *F*_*ST*_ in each comparison and defined loci with *F*_*ST*_ greater than five standard deviations from the mean as outlier loci (Axelsson et al., 2013; Willoughby, Harder, Tennessen, Scribner, & Christie, 2018). We identified the corresponding genes on which outlier loci were located by mapping outlier loci to the sea lamprey genome (Smith et al., 2013; Smith et al., 2018) and calculated the frequency of all alleles at each outlier locus.

## Results

We performed multiple independent toxicological assays over a two-year period using larvae collected from historically untreated (native range) and treated tributaries (Figure 1a; Table S1). The larvae collected from treated tributaries were not previously exposed to TFM prior to the toxicological assays (i.e., any resistance, if detected, could not be due to inducible tolerance (Hua, Morehouse, & Relyea, 2013)). If outright resistance has evolved, we would expect to observe higher survivorship in sea lamprey collected from populations with a history of treatment (i.e., Lake Michigan or Lake Champlain) in comparison to sea lamprey collected from areas that have never been treated (i.e., the native range, Connecticut River). We found no evidence to support this prediction. In 2016 trials (TOX 1-2; Table S2), the mortality for treated individuals was highly similar across all populations over a 12-hour exposure period. The combined mortality for treated individuals was 96.4%, 100%, and 91.1% for Lake Michigan, Lake Champlain, and Connecticut River populations, respectively (Figure 2a). There was no mortality within the control group for any population. We observed statistically significant, but not biologically meaningful, differences in survivorship among populations over time (*p*-value = 0.019) where post-hoc pairwise comparisons using the log-rank test revealed that individuals from the native, Connecticut River population had slightly higher survivorship than individuals from either Lake Michigan or Lake Champlain populations (*p*-value = 0.031). However, these slight differences in survivorship only translated to three additional Connecticut River individuals surviving until the end of the experiment. In our 2017 trials (TOX 3-5; Table S2), survivorship through time again did not vary substantially among populations (Figure S1) and estimates of lethal concentrations required to kill 50% of the population (LC50) varied only slightly among populations and trials (Figure 2b). The Lake Michigan population had lower estimated lethal concentration values for 25% and 50% mortality (LC25: mean 0.74 ± s.e. 0.01; LC50: mean 0.78 ± s.e. 0.01) than for larvae collected from the native range, Connecticut River (LC25: mean 1.39 ± s.e. 0.17; LC50: mean 1.48 ± s.e. 0.24) (Figure 2b). The lethal concentration values for the Connecticut River population were larger than that of the Lake Michigan population at both LC25 and LC50 values. Furthermore, no individuals from Lake Michigan survived TFM concentrations greater than 2 mg/L (Figure S1). When TFM is applied in the field, stream concentrations of TFM range from 5.5 to 9 mg/L for the pH and alkalinity values used in our experimental treatments (Figure 2b). In total, we performed five independent toxicological trials with six tested concentrations and never detected any meaningful differences in survival across populations (Table S2). From these analyses, we conclude that outright resistance to TFM has not yet evolved, as no treated individuals survived TFM exposure at concentrations they would be exposed to under ordinary stream treatment conditions.

**Figure 2.**
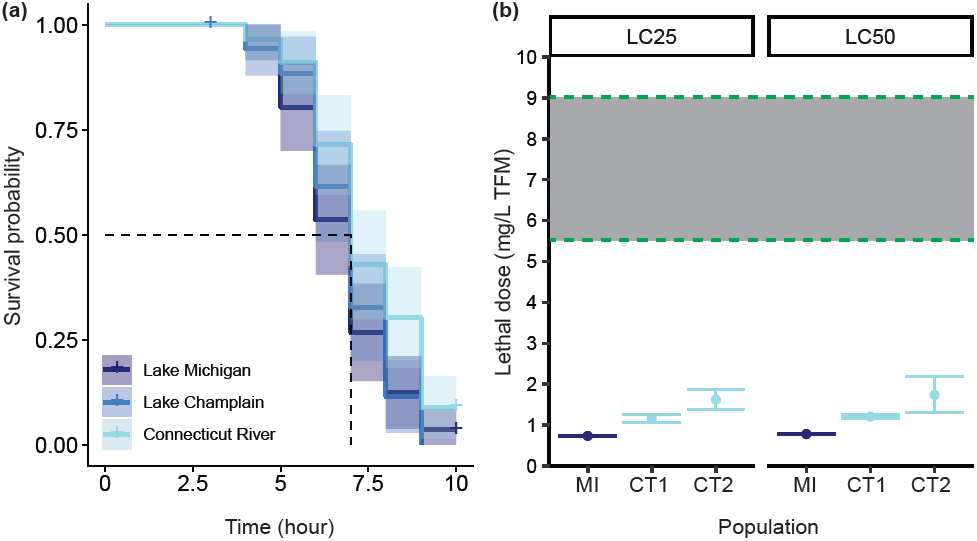
Results of toxicology assays illustrating survival through time and lethal concentrations. In 2016 trials, survival analyses revealed that previously unexposed larvae collected from treated populations did not survive at higher rates than larvae collected from the native population during exposure to lethal concentrations of TFM (a). In 2017 trials, independent toxicology assays from additional larvae collected from Lake Michigan (MI) and Connecticut River (native range; CT) again revealed that larvae collected from populations with a history of exposure did not survive lethal doses of TFM at higher rates than larvae collected from the native range (b). Lethal concentrations for 25% (LC25) and 50% (LC50) mortality were calculated and two separate assays were performed for Connecticut River individuals (CT1 and CT2). Dashed green lines border the range of TFM values larvae would be exposed to under typical stream application conditions. Collectively, these results illustrate that resistance to lethal concentrations of TFM has not yet evolved.

By contrast, we observed remarkably large differences in patterns of gene expression among sea lamprey from different populations exposed to the same, sublethal concentration of TFM. Briefly, we exposed 56 larvae from each population to either 0.2 or 0.3 mg/L of TFM for 6 hours (Table S2). An equal number of control individuals per population were included (i.e., individuals not exposed to TFM), individuals from all populations had the exact same acclimation history (4 months), and all populations were simultaneously included in each of two large experiments to avoid confounding circadian (Conesa et al., 2016) or environmental effects (Table S2, Table S3). Remarkably, when comparing samples treated with 0.3 mg/L TFM to control samples, we found 336 genes that were differentially expressed (275 upregulated, 61 downregulated in comparison to control individuals) in muscle tissue in the Lake Michigan population, dwarfing the number of differentially expressed genes (DEGs) found in both Lake Champlain (n = 21) and Connecticut River (n = 68) populations (Figure 3a-c; Table S4). No DEGs were shared among all three populations and different population responses were observed when lower concentrations (i.e., 0.2 mg/L) of TFM were applied and when different tissues were examined (Figures S2-S5).

**Figure 3.**
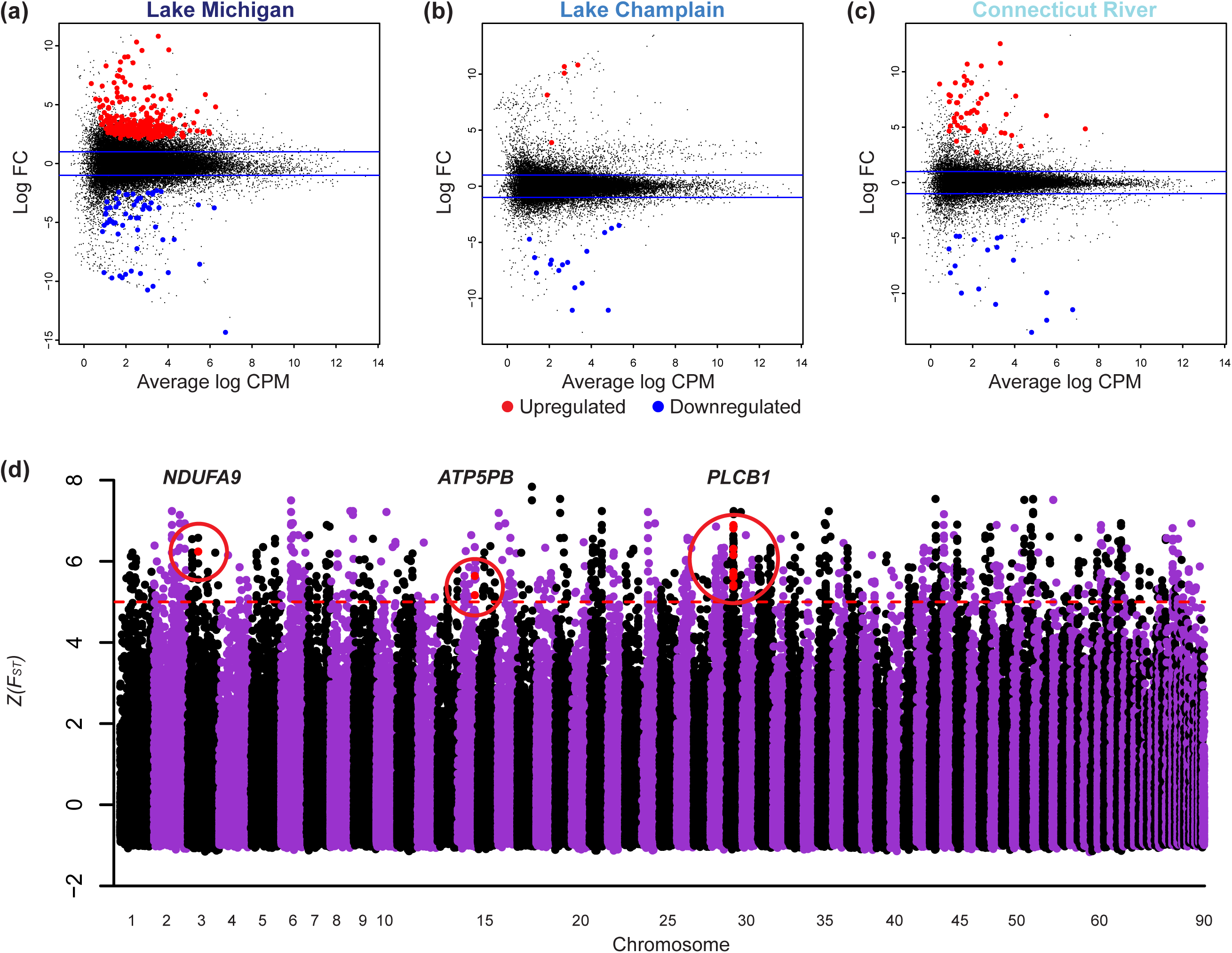
Transcriptomic and genomic evidence of incipient resistance (IR) in sea lamprey. A total of 336 genes were differentially expressed in response to sublethal concentrations of TFM in larval sea lamprey collected from Lake Michigan, the population with the longest history of TFM treatment (a). By contrast, only 21 and 68 genes were identified as differentially expressed in sea lamprey from Lake Champlain (b) and Connecticut River (native range) (c), respectively. After aligning reads back to the sea lamprey reference genome (sea lamprey have 99 chromosomes, 90 of which are assembled), calling SNPs and calculating *F*_*ST*_ between Lake Michigan and Connecticut River, three outliers relating to TFM’s primary mode of action, uncoupling the electron transport chain, were identified: *NDUFA9*, a gene encoding a subunit of complex I in the electron transport chain, *ATP5PB*, a gene encoding subunit b of ATP synthase, and *PLCB1*, a gene encoding phospholipase c beta 1.

The primary mode of action for TFM is to uncouple oxidative phosphorylation in the mitochondria resulting in severe ATP depletion and eventual mortality (Birceanu et al., 2011; Birceanu et al., 2009; Birceanu et al., 2014). Of the 336 DEGs identified in Lake Michigan, several genes were directly related to this mode of action including *CRCM1* (LogFC = 2.60), a calcium release-activated channel protein that controls the influx of calcium into cells when depleted, and *PLCD4* (LogFC = 2.41), an enzyme, phospholipase c delta 4, responsible for hydrolyzing phosphatidylinositol 4,5-bisphosphate into two secondary messenger molecules, one of which (inositol 1,4,5-trisphosphate or ‘IP3’) is known to release cellular calcium stores and increase calcium concentrations within the cell (Berridge, 1993). Upregulated expression of genes that increase intracellular concentrations of calcium could enhance oxidative phosphorylation in Lake Michigan larvae and partially compensate for the uncoupling effects of TFM (Figure 4a). In addition, calcium plays an integral role in activating and driving the oxidative phosphorylation cascade in mitochondria, and studies have demonstrated a positive relationship between nanomolar free calcium and the rate of phosphorylation (Fink, Bai, Yu, & Sivitz, 2017; Glancy, Willis, Chess, & Balaban, 2013). In gene ontology (GO) hierarchy networks, we also found regulation of calcium ion transport, regulation of calcium ion transmembrane transport, regulation of calcium ion transmembrane transporter activity, and regulation of voltage-gated calcium channel activity as significant biological processes (Figure 305 S6).

**Figure 4.**
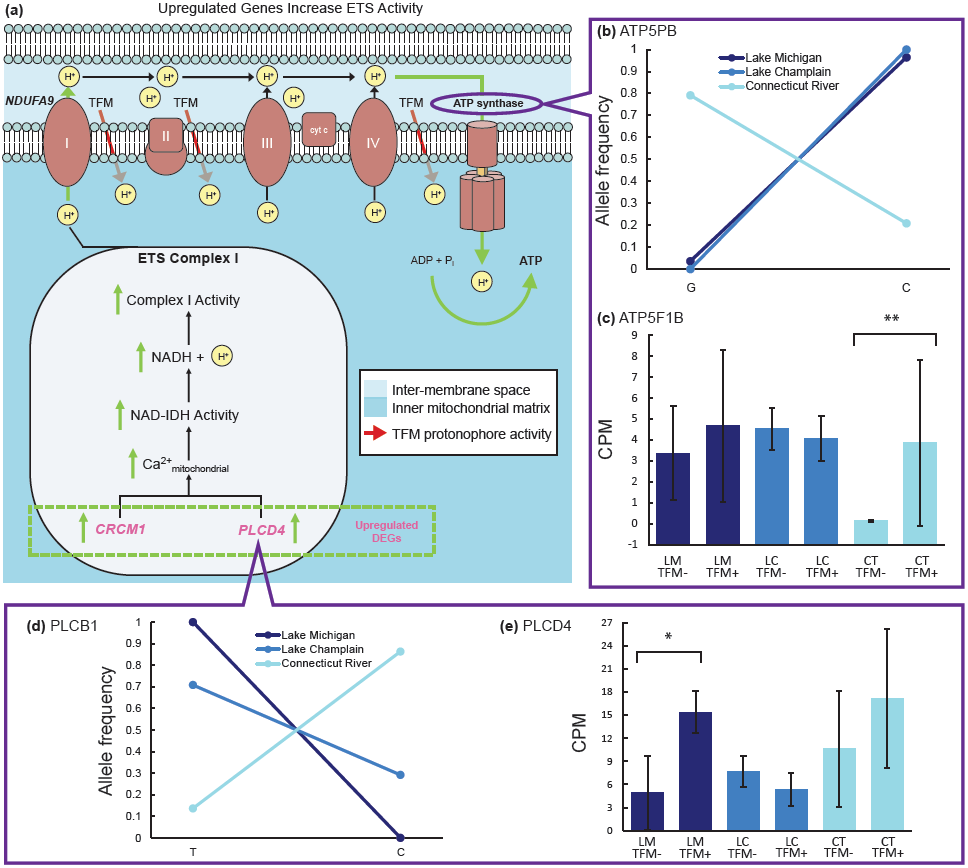
The pesticide TFM’s primary mode of action is to act as a protonophore, translocating protons from the inter-membrane space back into the inner mitochondrial matrix, thus disrupting oxidative phosphorylation (a). Several genes upregulated in Lake Michigan, the population with the longest history of exposure, may play a role in responding to TFM’s mode of action. For example, the upregulation of *CRCM1* and *PLCD4* increases intracellular concentrations of calcium and likely enhance oxidative phosphorylation in Lake Michigan ammocoetes when exposed to TFM (a, e). The gene *ATP5PB*, which encodes subunit b of ATP synthase and was identified as a putatively adaptive outlier (Fig. 3d), is nearly fixed at a bi-allelic SNP in the treated Lake Michigan and Lake Champlain populations, but segregates for different alleles in the native-range sea lamprey (b). A gene encoding an additional subunit of ATP synthase, *ATP5F1B*, is canalized for high expression in treated populations (i.e., the gene is expressed at high levels in both treatment (TFM+) and control (TFM-) individuals from the treated Lake Michigan (LM) and Lake Champlain (LC) populations) (c). By contrast, this gene is not constitutively upregulated in native-range, Connecticut River (CT) sea lamprey, but responds plastically to treatment. The gene *PLCB1*, which encodes phospholipase c beta 1 and was also identified as an outlier (Fig. 3d), is nearly fixed for alternative alleles at a bi-allelic SNP in the treated Lake Michigan and the native-range sea lamprey populations (d). A directly related gene encoding phospholipase c delta 4, *PLCD4*, shows significantly different levels of expression between control (TFM-) and treatment (TFM+) in Lake Michigan (e). Error bars depict standard deviation, *p*-value calculated from FDR-corrected tests of differential gene expression.

We next aligned our reads to the sea lamprey genome (Smith et al., 2013; Smith et al., 2018), called single nucleotide polymorphisms (SNPs), and identified outliers based on Z-scores of genetic divergence (Weir and Cockerham’s *F*_*ST*_) (Weir & Cockerham, 1984; Willoughby et al., 2018). Out of 554,685 SNPs detected with muscle tissue samples, an outlier, *ATP5PB*, was found in the comparison between Lake Michigan and Connecticut River populations and had an *F*_*ST*_ value greater than five standard deviations from the mean (Figure 3d). *ATP5PB* encodes a subunit of ATP synthase (subunit b) and, similar to many of the genes identified by expression analyses, appears to be responding to strong selection in order to restore normal oxidative phosphorylation in response to TFM’s primary mode of action (Figure 4a). Closer examination of the genotypes at the bi-allelic SNP found in this gene illustrates that a single allele is nearly fixed in both Lake Michigan and Lake Champlain, but only occurs rarely in the native range (Figure 4b). This outlier was also observed in comparisons between Lake Champlain and Connecticut River populations across tissue types (Figure S7, Figure S9). Although the native-range, Connecticut River lamprey are not fixed at the genetic level for this variant, they were able to respond plastically with respect to TFM at ATP synthase. The gene *ATP5F1B*, which codes for subunit beta of ATP synthase, was significantly upregulated in Connecticut River lamprey in response to TFM treatment (LogFC = 4.68, FDR-corrected *p*-value < 0. 0072, Figure 4c) when compared to unexposed, control individuals, suggesting a plastic response. Interestingly, *ATP5F1B* was upregulated in both control and treatment individuals from Lake Michigan and Lake Champlain in comparison to Connecticut River control individuals, suggesting a genetic response to constitutively increase expression of this gene.

An additional gene, *PLCB1*, was identified as an outlier between Lake Michigan and Connecticut River populations and had an *F*_*ST*_ value greater than six standard deviations from the mean (Figure 3d). Similar to the differentially expressed *PLCD4* (LogFC = 2.41; Figure 4a), *PLCB1*, a nearly identical variant of *PLCD4*, encodes an enzyme catalyzing the production of inositol 1,4,5-trisphosphate (‘IP3’). Alternative alleles are almost fixed at this bi-allelic SNP in Lake Michigan and Connecticut River populations, and similar patterns of allele frequency are observed in Lake Michigan and Lake Champlain (Figure 4d). This outlier was consistently detected in comparisons between Lake Michigan and Connecticut River across tissue types (Figure S8). *PLCD4* was upregulated in treated individuals in Lake Michigan. Lastly, an outlier, *NDUFA9*, a gene encoding complex I in the electron transport chain (ETC) was found in comparisons between Lake Michigan and Connecticut River and had an *F*_*ST*_ value greater than six standard deviations from the mean (Figure 3d), but not in comparisons between Lake Champlain and Connecticut River. The remaining 116 outlier genes (Figure 3d) have little to do with TFM’s known mode of action, many of which appear to be related to adaptation to the recently-colonized environments.

## Discussion

Here we provide evidence for the evolution of incipient resistance in invasive sea lamprey treated with a pesticide. We found 336 genes differentially expressed in response to TFM in larvae from Lake Michigan, the population with the longest history of using TFM (56 years). This transcriptional response represents a nearly 5-fold greater number of differentially expressed genes in comparison to larvae collected from the native range. The pesticide TFM’s primary mode of action is to serve as a protonophore, translocating protons across the inner mitochondrial membrane, thus uncoupling oxidative phosphorylation and depleting ATP. Many of the DEGs identified in Lake Michigan represent direct or indirect responses to ATP depletion. We also identified genomic evidence of a response to selection imposed by TFM. In comparisons between Lake Michigan and Connecticut River, the gene coding for ATP synthase subunit b, *ATP5PB*, had an *F*_*ST*_ value greater than five standard deviations from the mean (Figure 3d). These large shifts in allele frequencies indicate a response to selection in a gene that are critical to successful oxidative phosphorylation. We also identified a directly-related gene, *ATP5F1B*, that was differentially expressed in the native Connecticut River population, but not the other, treated populations. Interestingly, this gene is constitutively expressed at high levels in treated populations, suggesting that an additional component of ATP synthase has responded to pesticide-induced selection. Thus, the two populations with a lengthy history of TFM treatment show adaptive responses in ATP synthase at the genomic level, while the naïve, native population shows plastic responses in transcription of the same enzyme in response to TFM.

Two additional genes, *NDUFA9* and *PLCB1*, show genetic signatures of a response to TFM-mediated selection (Figure 3, Figure 4). *NDUFA9* encodes a subunit of the ETC complex I, and genetic modifications to this protein may allow for the more efficient transport of protons when inundated with TFM. *PLCB1* is known to increase calcium concentrations within the cell (Berridge, 1993), which could further assist with the transport of protons into the mitochondrial inter-membrane space when cells are confronted with TFM. A nearly identical variant, *PLCD4*, plastically increases expression in Lake Michigan and (albeit not significantly) in the native range in response to TFM (Figure 4e) whereas the expression of *PLCD4* does not respond to TFM in Lake Champlain. The observation that *NDUFA9* was not identified as a genetic outlier in comparisons between Lake Champlain and the native range (Connecticut River) and that *PLCD4* does not change expression in the presence of TFM in Lake Champlain, suggests that modifications to the ETC complex I are more extensive in Lake Michigan, which has been treated with TFM for a longer period of time.

Lake Michigan and Lake Champlain have been treated with TFM for 56 and 26 years, respectively. Both show evidence of selection at *ATP5PB*, however, both responded to TFM differently at the gene expression level with 336 and 21 genes differentially expressed in Lake Michigan and Lake Champlain, respectively. These large differences in the number of genes differentially expressed may reflect different evolutionary responses to TFM, one of which is mediated exclusively at the genetic level and one that is mediated at both genetic and transcriptomic levels. Alternatively, these differences may reflect the deeper evolutionary histories between these two populations or the different duration and intensities of pesticide usage (Whitehead, Clark, Reid, Hahn, & Nacci, 2017). We also observed large differences in gene expression depending on the concentration of TFM used and the tissue examined (Figures S2-S5). These similarities in adaptive, genetic responses and differences in broad patterns of gene expression suggest that the TFM-mediated selection results in a convergent, genetic response while the different evolutionary and treatment histories of these populations affect overall patterns of gene expression.

We identified large genetic and transcriptomic differences among populations but did not detect any individuals able to survive concentrations of TFM applied to streams in practice (Figure 2). Thus, we may be documenting the earliest stages of resistance evolving in treated populations (i.e., incipient resistance). Survival and reproductive advantages by larvae exposed to sublethal concentrations of TFM, for example individuals at the peripheries of TFM treatment or individuals buried deeper in the sediment, are inherited by their offspring and have spread throughout the treated populations. It may only be a matter of time before further mutations allow individuals to survive higher concentrations of TFM. In fact, recent modelling efforts have revealed that given the estimated strength of selection imposed by lamprey control, full-fledged resistance to TFM is predicted to occur soon (Christie et al., 2019). Because of the high gene flow and lack of population structure found throughout the Great Lakes (i.e., Lake Michigan is one, large panmictic population, see *SI Materials and Methods* for details), fully-resistant individuals may already be spreading throughout the population even though it may take many more years for successful detection (Christie et al., 2019). Alternatively, it may be that our toxicological assays were simply not sensitive enough to detect resistance to lethal concentrations of TFM. From a pragmatic standpoint, the detection of incipient resistance (IR) means that the development of alternative control measures and the implementation of adaptive management strategies focusing on delaying resistance should be a priority for the continued control of invasive sea lamprey. The detection of resistance in its earliest stages, as documented here, allows time for such actions to assist with the continued restoration of native and commercially-important fishes. Regardless of the system, the detection of incipient resistance can greatly assist in efforts to mitigate and delay the costly consequences of resistance.

## Acknowledgments

We thank B.J. Allaire, Bryan Apell, William Ardren, Steve Gephard, Nick Johnson, Robert Stira, Bradley Young, the Conte Anadromous Fish Research Laboratory, Michigan DNR, Connecticut Department of Energy and Environmental Protection, Hammond Bay Biological Station, and U.S. Fish & Wildlife for assistance with collecting larval lamprey. We thank Lori Criger, Sam Guffey, Elizabeth LaRue, Phillip San Miguel, Bob Rode, Mike Siefkes, Jeramiah Smith, Benson Solomon, and the Purdue Genomics Core for assistance with experiments and analyses. Matthew Byrnes, Matthew Campbell, Lindsey Dice, Jessica Elliott, Megan Gannon, Nathan Greiling, Kimberly Gulbranson, Jason Jaworski, Tara Novak, Ashe Owens, Emily Reverman, and Cassidy Robinson aided in the second set of toxicological trials. This research was funded by the Great Lakes Fishery Commission Project ID: 2016_CHR_54053.

## Data Accessibility Statement

Code and scripts are available at https://github.com/ChristieLab/sea_lamprey_TFM. Trimmed reads will be available via https://www.ncbi.nlm.nih.gov/sra upon acceptance of manuscript.

## Author Contributions

MRC, ASM, and MSS designed the project, all authors provided logistic support, ASM performed molecular analyses, XY, AMH, and ASM performed gene expression and genomic analyses, ASM and MMS performed survival analyses, and XY, ASM, and MRC wrote the manuscript with input from all authors.

## Competing Interests

The authors declare no competing interests.

