## Supporting Information for "Incipient resistance to an effective pesticide results from genetic adaptation and the canalization of gene expression"

### Materials and Methods

#### *Sea lamprey life history*

The anadromous sea lamprey (*Petromyzon marinus*) has a bipartite life cycle consisting of a filter-feeding larval stage and a parasitic adult stage. Throughout its native range, which includes much of the Northern Atlantic, the larval sea lamprey (also known as ammocoetes) remain buried in detritus-rich substrate where they filter feed for an average of 5 to 8 years (documented range 2 to 19 years) (Renaud, 2011). The metamorphosis of larval sea lamprey into adults involves substantial behavioral and physiological modifications required for their hematophagous, parasitic lifestyle (Youson, 2003). After metamorphosis, juvenile sea lamprey enter the ocean and begin searching for hosts. Sea lamprey have been observed feeding on a diverse array of species such as herring, mackerel, and salmon (Kottelat & Freyhof, 2007). Sea lamprey attach to their hosts with rows of sharp teeth and they secrete anticoagulants that allow them to feed continuously (Gage & Gage-Day, 1927). Throughout their native range, sea lamprey typically do not kill their hosts and an individual lamprey will often switch between hosts (Kottelat & Freyhof, 2007). After feeding for 20 to 36 months, the lamprey mature and begin to search for rivers and streams suitable for spawning, migrating anywhere between 20 to 850 km inland. Unlike other anadromous fishes, such as salmon, sea lamprey do not exhibit natal philopatry. Instead, sea lamprey cue in on chemicals produced by larvae living in freshwater streams and rivers (Bjerselius et al., 2000; Sorensen & Hoye, 2007). This reproductive strategy means that sea lamprey populations are largely panmictic, though there is evidence for restricted gene flow between populations on opposite sides of the Atlantic Ocean (Bryan et al., 2005;

Waldman, Grunwald, & Wirgin, 2008). Sea lamprey are strictly semelparous; after spawning all adults die.

In the Great Lakes, sea lamprey retain most of these life history characteristics, however, there are some notable exceptions. First, metamorphosed sea lamprey treat the Great Lakes as a surrogate ocean, thus they never physiologically acclimate to a salt-water environment. Second, growth rates are faster, the larval duration is shorter, adults are smaller, and fecundity is slightly lower (Heinrich et al., 2003; Young, Kelso, & Weise, 1990). Lastly, sea lamprey in the Great Lakes spend a longer period attached to a single host and an individual host can have many more sea lamprey attached to them than host species found in the Atlantic Ocean (Hansen et al., 2016). This last difference, possibly due to higher adult lamprey abundance and/or a greater lamprey to host ratio, means that sea lamprey are directly responsible for mortally wounding large numbers of host fishes in the Great Lakes (Farmer & Beamish, 1973). Even if invasive sea lamprey do not directly kill their hosts, many host fish later die from subsequent fungal infections associated with lamprey parasitism and fish that continue to survive have been shown to have decreased reproductive success (Swink, 1990; Swink & Hanson, 1989).

***Sample collection and experimental design***

Upon arrival at the Aquaculture Research Laboratory at Purdue University, ammocoetes were acclimated into 24, 30-gallon flow-through holding tanks corresponding to population of origin at similar densities of 60-67 individuals per tank. Each tank received flow-through well water and was lined with 10 cm of sand to allow ammocoetes to burrow. During acclimation, ammocoetes were fed a mixture of baker’s yeast and water (1 gram of yeast per ammocoete) two times per week. Larval mortality in the holding tanks was minimal (< 2.1% across all populations, n = 30 individuals) during the acclimation period.

For the sublethal and lethal effects studies, we constructed an experimental array to conduct exposure trials in a temperature-controlled environmental chamber (12.8°C) consisting of 36, 2.5-gallon glass aquaria. Each tank received freshwater (average temperature = 13.9°C, pH = 8.3) that was gravity-fed from storage tanks on top of the array. Alkalinity within experimental tanks was maintained within a constant range across exposures (96-140 ppm) by mixing well water (alkalinity > 200 ppm) with water treated via reverse osmosis (alkalinity ~ 20 ppm). Each treatment tank also received an inflow line from one of two adjacent multi-channel peristaltic pumps (Gilson MINIPULS Evolution; Fisherbrand FH100M). A single air stone was suspended in the center of each tank to provide aeration and promote mixing.

***RNA-seq and transcriptome assembly***

We first trimmed RNA-seq reads using Trimmomatic v0.36 (Bolger, Lohse, & Usadel, 2014) with suggested parameters (MacManes, 2014), and then used FastQC to ensure that read quality was acceptable for transcriptome assembly. We normalized trimmed reads in silico and assembled our multi-tissue transcriptome (i.e., all muscle, liver, and brain samples from GE 1; Table S2, Table S3) de novo using Trinity (v2.5.1) (Haas et al., 2013) with the following command: Trinity --seqType fq --max_memory 512G --samples_file samples.txt --CPU 20 --full_cleanup --SS_lib_type RF. Once the assembly was complete, we checked assembly quality and completeness with BUSCO (Simão, Waterhouse, Ioannidis, Kriventseva, & Zdobnov, 2015) using ‘eukaryota’ (eukaryotes) as the species clade rather than the ‘vertebrata’ group, as sea lamprey are explicitly excluded in the orthologous gene set for vertebrates due to high sequence divergence (Simão et al., 2015).

Next, we annotated transcripts using the Trinotate bioinformatics annotation protocol (http://trinotate.github.io) (Bryant et al., 2017). Briefly, this annotation pipeline leverages BLASTX and BLASTP (after predicting probable coding regions within the transcripts using TransDecoder [http://transdecoder.github.io]) to search for similarities between transcripts and proteins in both Swiss-Prot and Uniref90 protein databases (downloaded June 5, 2018). Additionally, Trinotate searches for conserved protein domains and signal peptides within the transcript sequences. Gene ontology (GO) terms were associated with Trinity transcripts by matching proteins in the Swiss-Prot database with proteins predicted from transcripts by TransDecoder.

***Genomic analysis***

First, we called SNPs following the RNA-seq pipeline. We started by mapping RNA-seq reads to the sea lamprey genome (Smith et al., 2013; Smith et al., 2018) following the STAR 2-pass alignment steps. We set --outFilterMultimapNmax to 1, --outSJfilterReads to Unique, --alignEndsType to EndToEnd, --chimMainSegmentMultNmax to 1, --limitGenomeGenerateRAM to 250000000000, --sjdbOverhang to 149, --runThreadN to 19, and --limitSjdbInsertNsj to 25000000. All other parameters were set to default values. By adding read groups, sorting, marking duplicates, and creating indices, we obtained BAM files from SAM files generated by the STAR 2-pass alignment steps. After generating BAM files, we applied the GATK tool, SplitNCigarReads, to BAM files, which split reads into exon segments and cut sequences extending to intronic regions. We next called variants using the GATK tool, HaplotypeCaller, in which we picked the option -dontUseSoftClippedBases, and set -stand_call_conf to 20.0 and -

maxAltAlleles to 100. Finally, we filtered variants using the GATK tool, VariantFiltration, by setting -window to 35 and -cluster to 3 and applying filters "FS > 30.0" and "QD < 2.0", and removed indels. We additionally called SNPs following the GATK DNA-seq pipeline. The differences between the RNA-seq and DNA-seq pipelines begin with the application of HaplotypeCaller. In comparison to the RNA-seq pipeline, we set --genotyping_mode to DISCOVERY, --emitRefConfidence to GVCF, --variant_index_type to LINEAR, --variant_index_parameter to 128000, -pairHMM to VECTOR_LOGLESS_CACHING, -ploidy to 2, and -maxAltAlleles to 100. Finally, we performed joint genotyping using the GATK tool, GenotypeGVCFs, with --max_alternate_alleles set to 100, and removed all indels.

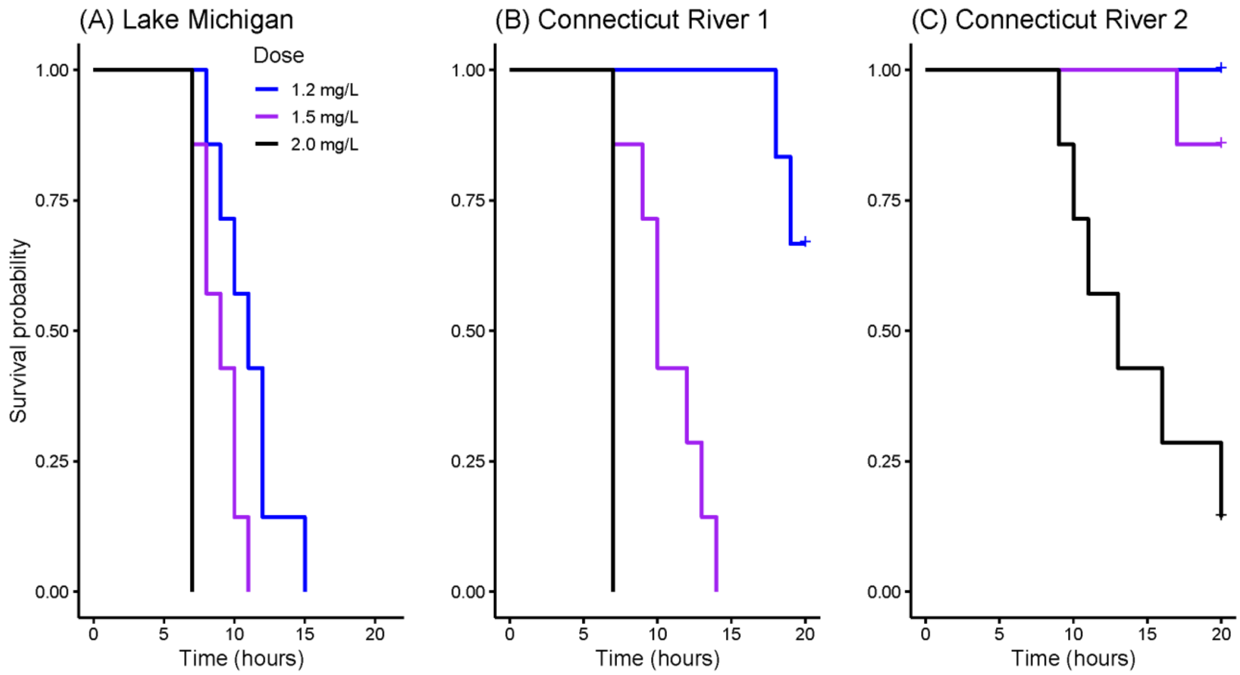


**Figure S1.** Survival probability through time for three trials (TOX 3,4,5; Table S2) conducted during the second set of toxicological assays (2017). Plots are arranged by population where panel A depicts the historically TFM-treated population (Lake Michigan, TOX 4), and panels B and C depict the TFM-naïve population (Connecticut River, TOX 3,5). The blue line represents the survival probability of sea lamprey larvae at a given time point at the TFM concentrations of 1.2 mg/L, the purple line at 1.5 mg/L, and the black line at 2.0 mg/L. Additional doses of 0 and 0.5 mg/L were excluded because all individuals in all trials survived until the end of the experiment. Sample sizes represent 21 individuals per dose (7 individuals in each of 3 replicate aquaria).

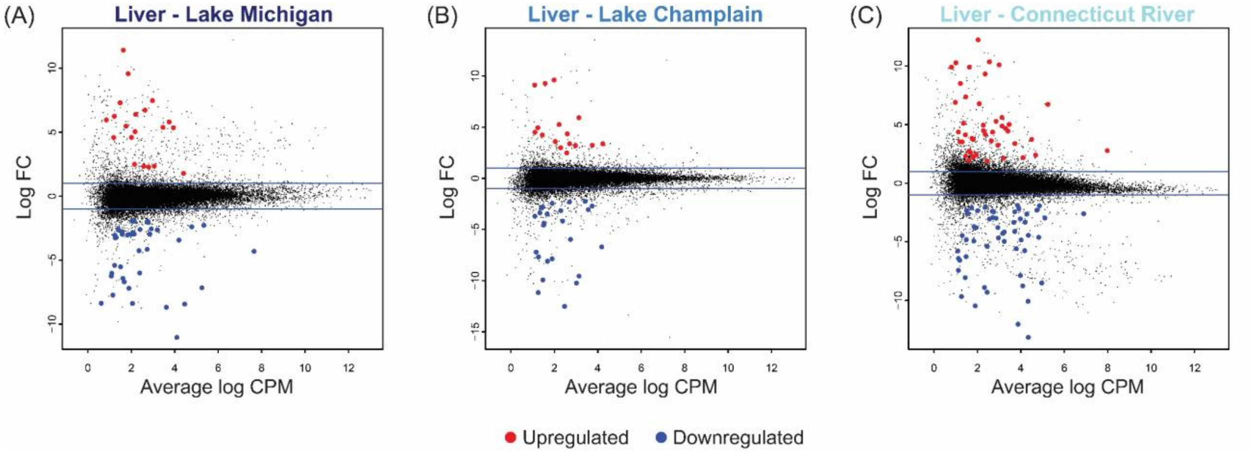


**Figure S2.** Gene expression experiment with liver tissue shows that a total of 60, 41 and 99 genes were differentially expressed in response to sublethal concentrations of TFM (0.2 mg/L) in larval sea lamprey collected from Lake Michigan (A), Lake Champlain (B) and Connecticut River (C), respectively. (LM control: n=3, LM treated: n=4, LC control: n=3, LC treated: n=3, CT control: n=3, CT treated: n=3; see Table S3 for the identities of all individuals, Table S5 for the identities of all genes).

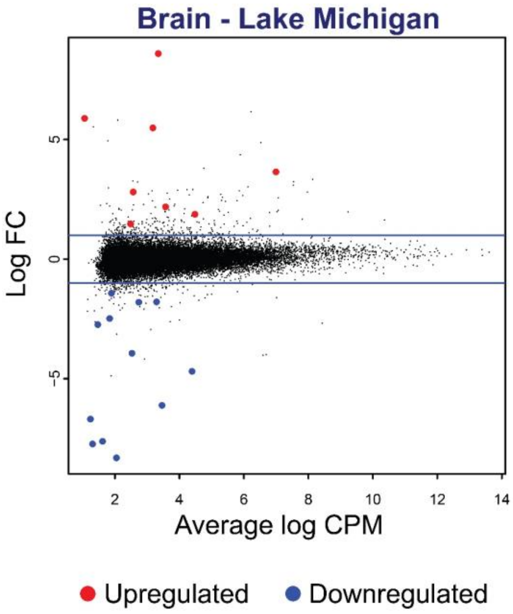


**Figure S3.** Gene expression experiment with brain tissue shows that a total of 20 genes were differentially expressed in response to sublethal concentrations of TFM (0.2 mg/L) in larval sea lamprey collected from Lake Michigan. (LM control: n=4, LM treated: n=4; see Table S3 for the identities of all individuals, Table S6 for the identities of all genes).

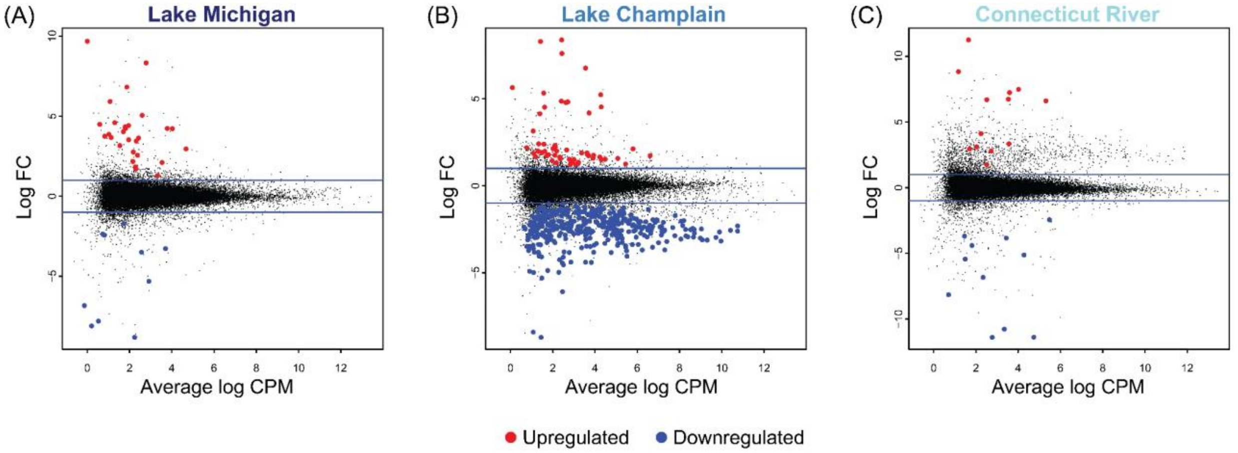


**Figure S4.** Gene expression experiment with muscle tissue shows that a total of 38, 451 and 24 genes were differentially expressed in response to a lower, sublethal concentration of TFM (0.2 mg/L) in larval sea lamprey collected from Lake Michigan (A), Lake Champlain (B) and Connecticut River (C), respectively. (LM control: n=6, LM treated: n=6, LC control: n=5, LC treated: n=6, CT control: n=5, CT treated: n=6; see Table S3 for the identities of all individuals, Table S7 for the identities of all genes). Sequencing machine was included as a fixed effect to identify differentially expressed genes using edgeR's GLM functionality in R. Lake Champlain has an unusually large number of downregulated genes in response to TFM treatment. Based on over-represented GO terms, many of these downregulated genes are related to regulation of biological processes, response to abiotic factors, metabolism, muscle morphogenesis, muscle development, muscle contraction, heart morphogenesis, heart development, and heart contraction. Furthermore, the percentage of downregulated genes was substantially higher in GE 2 compared to GE 1 (Lake Michigan: 0.26 vs. 0.18, Lake Champlain: 0.87 vs. 0.76, and Connecticut River: 0.46 vs. 0.28; Table S2). Although we could not identify the precise mechanism, we speculate that these fish were in poor condition, perhaps due to unusually high capture stress on the day of the experiment. Alternatively, different populations may respond differently to lower subleathal concentrations of TFM.

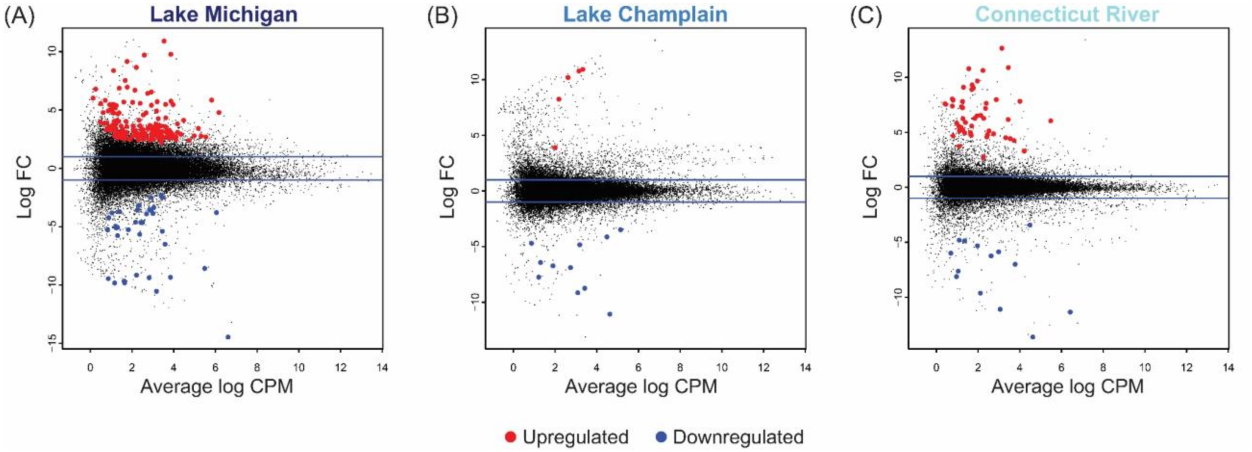


**Figure S5.** Gene expression analysis with muscle tissue exposed to sublethal concentrations of TFM (0.3 mg/L) in combined GE 1 and GE 2 experiments (Table S2). In comparison to the results illustrated in Figure 3a-c (main text), which only includes sea lamprey exposed to 0.3 mg/L of TFM from a single experiment (GE 1; Table S3), the analysis generating Figure S5 includes two additional sea lamprey exposed to 0.3 mg/L of TFM in a second experiment (GE 2; Table S3). A total of 202, 16 and 64 genes were differentially expressed in larval sea lamprey collected from Lake Michigan (A), Lake Champlain (B) and Connecticut River (C), respectively. We performed this additional analysis to verify that the patterns of gene expression do not change dramatically as the sample size goes up. (LM control: n=4, LM treated: n=4, LC control: n=2, LC treated: n=3, CT control: n=3, CT treated: n=3; Table S3). Sequencing machine was included as a fixed effect to identify differentially expressed genes using edgeR's GLM functionality in R.

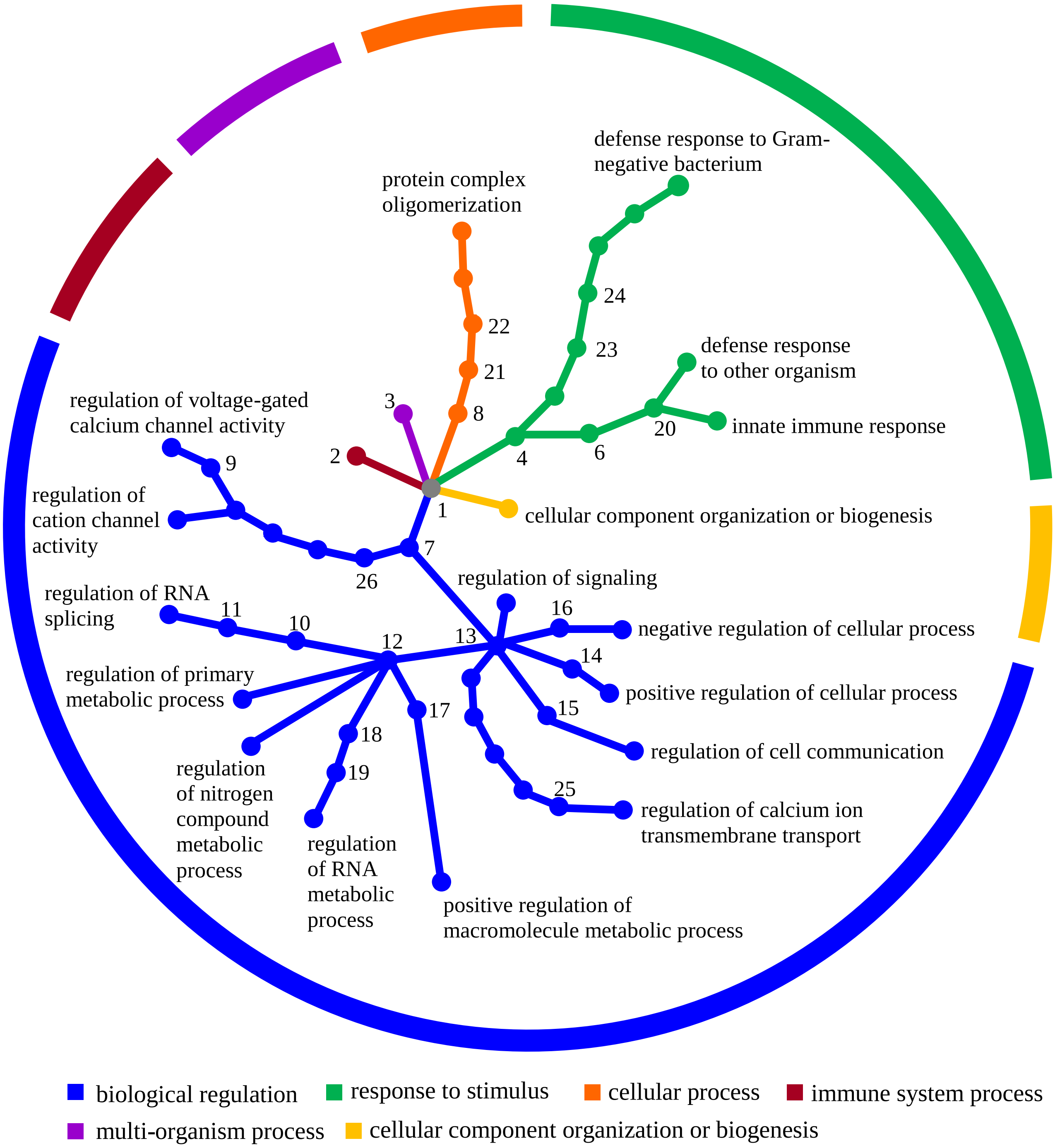


1. biological process; 2. immune system process; 3. multi-organism process; 4. response to stimulus; 6. response to stress; 7. biological regulation; 8. cellular process; 9. regulation of calcium ion transmembrane transporter activity; 10. regulation of macromolecule metabolic process; 11. regulation of gene expression; 12. regulation of metabolic process; 13. regulation of biological process; 14. positive regulation of biological process; 15. regulation of cellular process; 16. negative regulation of biological process; 17. positive regulation of metabolic process; 18. regulation of cellular metabolic process; 19. regulation of nucleobase-containing compound metabolic process; 20. defense response; 21. cellular component organization; 22. cellular component assembly; 23. response to external biotic stimulus; 24. response to other organism; 25. regulation of calcium ion transport; 26. regulation of molecular function

**Figure S6.** Gene ontology (GO) hierarchy networks constructed with top 100 GO terms identified with differentially expressed genes (FDR-corrected *p*-value = 0.01) using the metacoder package in R suggest biological regulation, response to stimulus, cellular process, immune system process, multi-organism process and cellular component organization or biogenesis as major biological processes in response to TFM in the Lake Michigan population (muscle tissue). Branch and node colors indicate the biological process child term to which distal nodes belong, with the central grey node representing the biological process level of the GO hierarchy. Nodes labeled with texts or numbers represent significant biological processes.

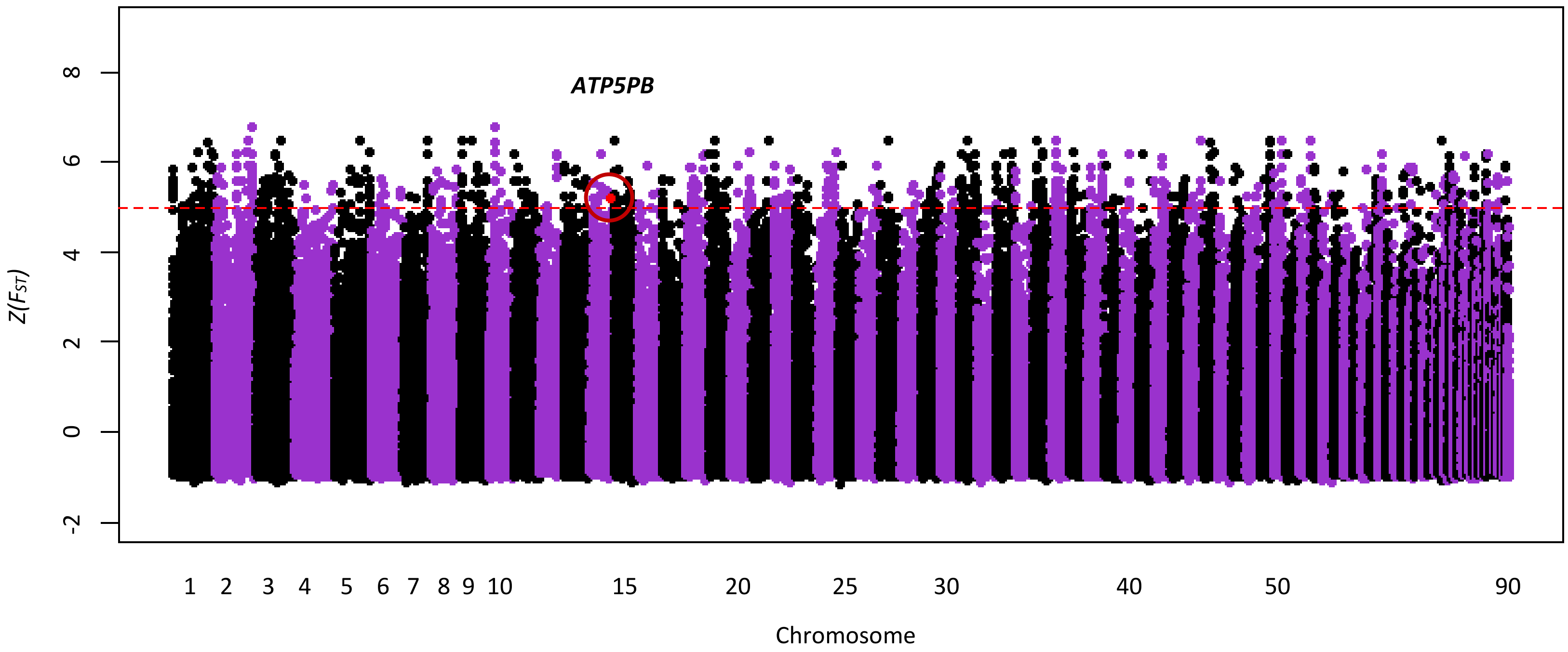


**Figure S7.** After aligning reads back to the sea lamprey reference genome (sea lamprey have 99 chromosomes, 90 of which are assembled), calling SNPs and calculating $F_{ST}$, an outlier SNP (5.22 standard deviations greater than the mean; red point) was also found in the comparison between Lake Champlain and Connecticut River for muscle tissue samples. This outlier SNP belongs to *ATP5PB*, a gene encoding subunit b of ATP synthase. In this comparison, *NDUFA9* and *PLCB1* were not identified as outliers (Z($F_{ST}$) < 5; cf. Figure 3).

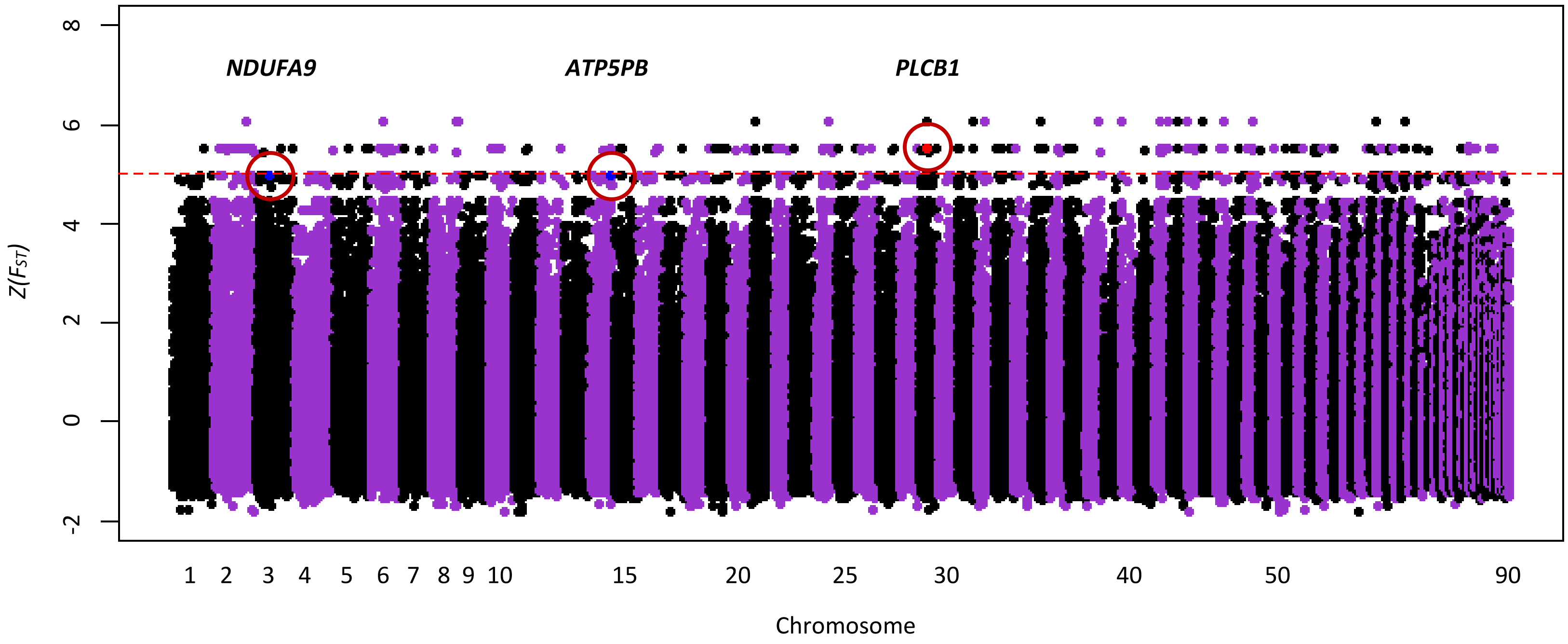


**Figure S8.** After aligning reads back to the sea lamprey reference genome (sea lamprey have 99 chromosomes, 90 of which are assembled), calling SNPs and calculating $F_{ST}$, an outlier SNP (5.51 standard deviations greater than the mean; red point) was found in the comparison between Lake Michigan and Connecticut River for liver tissue samples. This outlier SNP belongs to *PLCB1*, a gene encoding phospholipase c beta 1. In this comparison, *ATP5PB* and *NDUFA9* were not identified as outliers ($Z(F_{ST}) < 5$; cf. Figure 3), but had $Z(F_{ST})$ values very close to 5 (highlighted in blue). The white space at high $Z(F_{ST})$ values is due to small liver sample sizes compared to muscle sample sizes (n = 42 for muscle samples vs. n = 19 for liver samples; Table S3), which constrains the total number of possible $F_{ST}$ values.

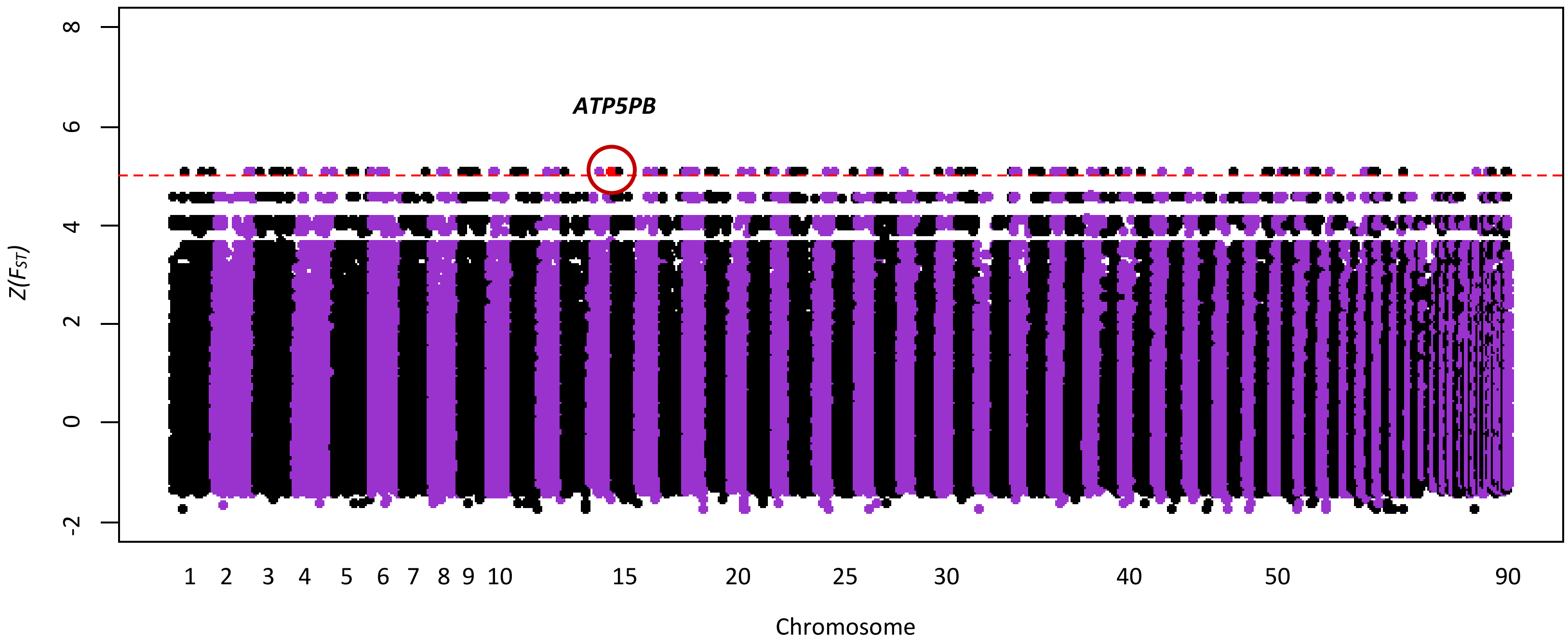


**Figure S9.** After aligning reads back to the sea lamprey reference genome (sea lamprey have 99 chromosomes, 90 of which are assembled), calling SNPs and calculating $F_{ST}$, an outlier SNP (5.11 standard deviations greater than the mean; red point) was found in the comparison between Lake Champlain and Connecticut River for liver tissue samples. This outlier SNP belongs to *ATP5PB*, a gene encoding subunit b of ATP synthase. In this comparison, *NDUFA9* and *PLCB1* were not identified as outliers (Z($F_{ST}$) < 5; cf. Figure 3). The white space at high Z($F_{ST}$) values is due to small liver sample sizes compared to muscle sample sizes (n = 42 for muscle samples vs. n = 19 for liver samples; Table S3), which constrains the total number of possible $F_{ST}$ values.

**Table S1.** Population, Collection Date, Collection Location and Sample Sizes.

| **Population** | **Date Collected** | **Collection Location** | **Latitude, Longitude** | **Sample Sizes** |
|---|---|---|---|---|
| Lake Michigan | 07/13/2016 | Manistee River, MI | 44.263651, -86.367834 | 565 |
| Lake Champlain | 07/11/2016-07/13/2016 | Corbeau Creek, NY | 44.903821, -73.536097 | 517 |
| Connecticut River | 07/14/2016-07/15/2016 | Connecticut River, CT | 41.331972, -72.293014 | 404 |
| Lake Michigan | 09/13/2017 | Platte River, MI | 44.709791, -86.150832 | 284 |
| Connecticut River | 09/18/2017 | Turner Falls, MA | 42.591592, -72.577404 | 265 |
| Total | | | | 2035 |

**Table S2.** Summary of Toxicological Assays and Gene Expression Experiments.

| Date | Collection Year | Type | TFM Concentrations | No. Replicate Tanks (without controls) | No. Control Tanks | No. Individuals per Tank | Total Sample Sizes | Duration of Exposure (hrs) | Population |
|---|---|---|---|---|---|---|---|---|---|
| 11/21/2016 | 2016 | GE 1 | 0.2, 0.3 mg/L | 24 | 12 | 7 | 252 | (6)12* | LM, LC, CT |
| 11/30/2016 | 2016 | GE 2 | 0.2, 0.3 mg/L | 24 | 6 | 7 | 210 | (6)12* | LM, LC, CT |
| 02/24/2017 | 2016 | TOX 1 | 3 mg/L | 24 | 12 | 7 | 252 | 12 | LM, LC, CT |
| 02/27/2017 | 2016 | TOX 2 | 1.67 mg/L | 18 | 9 | 5 | 135 | 12 | LM, LC, CT |
| 11/13/2017 | 2017 | TOX 3 | 0.5, 1.2, 1.5, 2 mg/L | 12 | 3 | 7 | 105 | 20 | CT |
| 11/15/2017 | 2017 | TOX 4 | 0.5, 1.2, 1.5, 2 mg/L | 12 | 3 | 7 | 105 | 20 | LM |
| 11/20/2017 | 2017 | TOX 5 | 0.5, 1.2, 1.5, 2 mg/L | 12 | 3 | 7 | 105 | 20 | CT |

Collection Year is the year of sample collection. TOX stands for toxicological assay; GE stands for gene expression experiment. LM, LC, and CT stand for Lake Michigan, Lake Champlain, and Connecticut River (Table S1), respectively. *Individuals collected for gene expression analyses (Table S3) were collected at hour 6; the remaining individuals were monitored for 12 hours to ensure no subsequent mortality occurred (i.e., sublethal exposure).

**Table S3.** Summary of Samples Sequenced with RNA-seq.

| Sample No. | ID | Experiment | Date | Population | Tissue type | TFM [] | Sequencing machine | # mapped reads |
|---|---|---|---|---|---|---|---|---|
| 402 | lmc_brn_1 | GE 1 | 11/21/2016 | Lake Michigan | brain | 0 | NovaSeq 6000 | 20457400 |
| 403 | lmc_brn_2 | GE 1 | 11/21/2016 | Lake Michigan | brain | 0 | NovaSeq 6000 | 24447032 |
| 409 | lmc_brn_3 | GE 1 | 11/21/2016 | Lake Michigan | brain | 0 | NovaSeq 6000 | 29118690 |
| 415 | lmc_brn_4 | GE 1 | 11/21/2016 | Lake Michigan | brain | 0 | NovaSeq 6000 | 24086639 |
| 407 | lmt_brn_1 | GE 1 | 11/21/2016 | Lake Michigan | brain | 0.2 | NovaSeq 6000 | 29861948 |
| 419 | lmt_brn_2 | GE 1 | 11/21/2016 | Lake Michigan | brain | 0.2 | NovaSeq 6000 | 23022615 |
| 422 | lmt_brn_3 | GE 1 | 11/21/2016 | Lake Michigan | brain | 0.2 | NovaSeq 6000 | 29152139 |
| 427 | lmt_brn_4 | GE 1 | 11/21/2016 | Lake Michigan | brain | 0.2 | NovaSeq 6000 | 17610659 |
| 402 | lmc_liv_1 | GE 1 | 11/21/2016 | Lake Michigan | liver | 0 | NovaSeq 6000 | 19802319 |
| 403 | lmc_liv_2 | GE 1 | 11/21/2016 | Lake Michigan | liver | 0 | NovaSeq 6000 | 23747347 |
| 409 | lmc_liv_3 | GE 1 | 11/21/2016 | Lake Michigan | liver | 0 | NovaSeq 6000 | 28566552 |
| 407 | lmt_liv_1 | GE 1 | 11/21/2016 | Lake Michigan | liver | 0.2 | NovaSeq 6000 | 28658889 |
| 419 | lmt_liv_2 | GE 1 | 11/21/2016 | Lake Michigan | liver | 0.2 | NovaSeq 6000 | 28876080 |
| 422 | lmt_liv_3 | GE 1 | 11/21/2016 | Lake Michigan | liver | 0.2 | NovaSeq 6000 | 25875166 |
| 427 | lmt_liv_4 | GE 1 | 11/21/2016 | Lake Michigan | liver | 0.2 | NovaSeq 6000 | 20142378 |
| 402 | lm_c_1 | GE 1 | 11/21/2016 | Lake Michigan | muscle | 0 | HiSeq 2500 | 11765624 |
| 403 | lm_c_2 | GE 1 | 11/21/2016 | Lake Michigan | muscle | 0 | HiSeq 2500 | 16989420 |
| 409 | lm_c_4 | GE 1 | 11/21/2016 | Lake Michigan | muscle | 0 | HiSeq 2500 | 12741348 |
| 415 | lm_c_5 | GE 1 | 11/21/2016 | Lake Michigan | muscle | 0 | NovaSeq 6000 | 30008269 |
| 407 | lm_tl_1 | GE 1 | 11/21/2016 | Lake Michigan | muscle | 0.2 | HiSeq 2500 | 13866543 |
| 419 | lm_tl_3 | GE 1 | 11/21/2016 | Lake Michigan | muscle | 0.2 | NovaSeq 6000 | 24738305 |
| 422 | lm_tl_4 | GE 1 | 11/21/2016 | Lake Michigan | muscle | 0.2 | NovaSeq 6000 | 28538823 |
| 427 | lm_tl_5 | GE 1 | 11/21/2016 | Lake Michigan | muscle | 0.2 | NovaSeq 6000 | 26030162 |
| 418 | lm_th_1 | GE 1 | 11/21/2016 | Lake Michigan | muscle | 0.3 | HiSeq 2500 | 10924994 |
| 423 | lm_th_2 | GE 1 | 11/21/2016 | Lake Michigan | muscle | 0.3 | HiSeq 2500 | 15576432 |
| 400 | lcc_liv_1 | GE 1 | 11/21/2016 | Lake Champlain | liver | 0 | NovaSeq 6000 | 28506290 |
| 401 | lcc_liv_2 | GE 1 | 11/21/2016 | Lake Champlain | liver | 0 | NovaSeq 6000 | 28588031 |
| 426 | lcc_liv_3 | GE 1 | 11/21/2016 | Lake Champlain | liver | 0 | NovaSeq 6000 | 30269008 |
| 404 | lct_liv_1 | GE 1 | 11/21/2016 | Lake Champlain | liver | 0.2 | NovaSeq 6000 | 29966156 |

| | | | | | | | | |
|---|---|---|---|---|---|---|---|---|
| 410 | lct_liv_2 | GE 1 | 11/21/2016 | Lake Champlain | liver | 0.2 | NovaSeq 6000 | 30003410 |
| 417 | lct_liv_3 | GE 1 | 11/21/2016 | Lake Champlain | liver | 0.2 | NovaSeq 6000 | 21071509 |
| 400 | lc_c_1 | GE 1 | 11/21/2016 | Lake Champlain | muscle | 0 | HiSeq 2500 | 11489986 |
| 401 | lc_c_2 | GE 1 | 11/21/2016 | Lake Champlain | muscle | 0 | HiSeq 2500 | 4771684 |
| 404 | lc_tl_1 | GE 1 | 11/21/2016 | Lake Champlain | muscle | 0.2 | NovaSeq 6000 | 25174478 |
| 410 | lc_tl_2 | GE 1 | 11/21/2016 | Lake Champlain | muscle | 0.2 | NovaSeq 6000 | 30140422 |
| 417 | lc_tl_3 | GE 1 | 11/21/2016 | Lake Champlain | muscle | 0.2 | NovaSeq 6000 | 24695462 |
| 405 | lc_th_1 | GE 1 | 11/21/2016 | Lake Champlain | muscle | 0.3 | HiSeq 2500 | 11612986 |
| 408 | lc_th_2 | GE 1 | 11/21/2016 | Lake Champlain | muscle | 0.3 | HiSeq 2500 | 14019682 |
| 420 | lc_th_3 | GE 1 | 11/21/2016 | Lake Champlain | muscle | 0.3 | HiSeq 2500 | 14592886 |
| 412 | ctc_liv_1 | GE 1 | 11/21/2016 | Connecticut River | liver | 0 | NovaSeq 6000 | 20933885 |
| 413 | ctc_liv_2 | GE 1 | 11/21/2016 | Connecticut River | liver | 0 | NovaSeq 6000 | 32367561 |
| 424 | ctc_liv_3 | GE 1 | 11/21/2016 | Connecticut River | liver | 0 | NovaSeq 6000 | 19769485 |
| 411 | ctt_liv_1 | GE 1 | 11/21/2016 | Connecticut River | liver | 0.2 | NovaSeq 6000 | 17750173 |
| 414 | ctt_liv_2 | GE 1 | 11/21/2016 | Connecticut River | liver | 0.2 | NovaSeq 6000 | 22848428 |
| 416 | ctt_liv_3 | GE 1 | 11/21/2016 | Connecticut River | liver | 0.2 | NovaSeq 6000 | 26290947 |
| 412 | ct_c_1 | GE 1 | 11/21/2016 | Connecticut River | muscle | 0 | HiSeq 2500 | 16566749 |
| 413 | ct_c_2 | GE 1 | 11/21/2016 | Connecticut River | muscle | 0 | HiSeq 2500 | 13439299 |
| 424 | ct_c_3 | GE 1 | 11/21/2016 | Connecticut River | muscle | 0 | HiSeq 2500 | 14310452 |
| 411 | ct_tl_1 | GE 1 | 11/21/2016 | Connecticut River | muscle | 0.2 | NovaSeq 6000 | 19107299 |
| 414 | ct_tl_2 | GE 1 | 11/21/2016 | Connecticut River | muscle | 0.2 | NovaSeq 6000 | 26711086 |
| 416 | ct_tl_3 | GE 1 | 11/21/2016 | Connecticut River | muscle | 0.2 | NovaSeq 6000 | 19192978 |
| 406 | ct_th_1 | GE 1 | 11/21/2016 | Connecticut River | muscle | 0.3 | HiSeq 2500 | 15215385 |
| 421 | ct_th_2 | GE 1 | 11/21/2016 | Connecticut River | muscle | 0.3 | HiSeq 2500 | 11272452 |
| 429 | ct_th_3 | GE 1 | 11/21/2016 | Connecticut River | muscle | 0.3 | HiSeq 2500 | 15014323 |
| 567 | lmc_mus_ge2_1 | GE 2 | 11/30/2016 | Lake Michigan | muscle | 0 | NovaSeq 6000 | 32472014 |
| 581 | lmc_mus_ge2_2 | GE 2 | 11/30/2016 | Lake Michigan | muscle | 0 | NovaSeq 6000 | 26929835 |
| 573 | lmt_mus_ge2_3 | GE 2 | 11/30/2016 | Lake Michigan | muscle | 0.2 | NovaSeq 6000 | 25405100 |
| 576 | lmt_mus_ge2_4 | GE 2 | 11/30/2016 | Lake Michigan | muscle | 0.2 | NovaSeq 6000 | 25323666 |
| 577 | lmt_mus_ge2_1 | GE 2 | 11/30/2016 | Lake Michigan | muscle | 0.3 | NovaSeq 6000 | 29819726 |
| 591 | lmt_mus_ge2_2 | GE 2 | 11/30/2016 | Lake Michigan | muscle | 0.3 | NovaSeq 6000 | 21902654 |
| 566 | lcc_mus_ge2_1 | GE 2 | 11/30/2016 | Lake Champlain | muscle | 0 | NovaSeq 6000 | 28348812 |

| | | | | | | | | |
|---|---|---|---|---|---|---|---|---|
| 572 | lcc_mus_ge2_2 | GE 2 | 11/30/2016 | Lake Champlain | muscle | 0 | NovaSeq 6000 | 31274558 |
| 580 | lcc_mus_ge2_3 | GE 2 | 11/30/2016 | Lake Champlain | muscle | 0 | NovaSeq 6000 | 29105459 |
| 571 | lct_mus_ge2_1 | GE 2 | 11/30/2016 | Lake Champlain | muscle | 0.2 | NovaSeq 6000 | 32878565 |
| 587 | lct_mus_ge2_2 | GE 2 | 11/30/2016 | Lake Champlain | muscle | 0.2 | NovaSeq 6000 | 15326492 |
| 588 | lct_mus_ge2_3 | GE 2 | 11/30/2016 | Lake Champlain | muscle | 0.2 | NovaSeq 6000 | 22899739 |
| 565 | ctc_mus_ge2_1 | GE 2 | 11/30/2016 | Connecticut River | muscle | 0 | NovaSeq 6000 | 21423126 |
| 583 | ctc_mus_ge2_2 | GE 2 | 11/30/2016 | Connecticut River | muscle | 0 | NovaSeq 6000 | 22933267 |
| 569 | ctt_mus_ge2_1 | GE 2 | 11/30/2016 | Connecticut River | muscle | 0.2 | NovaSeq 6000 | 23906705 |
| 579 | ctt_mus_ge2_2 | GE 2 | 11/30/2016 | Connecticut River | muscle | 0.2 | NovaSeq 6000 | 23538391 |
| 582 | ctt_mus_ge2_3 | GE 2 | 11/30/2016 | Connecticut River | muscle | 0.2 | NovaSeq 6000 | 23770608 |

**Table S4.** Differentially Expressed Genes Detected in Lake Michigan, Lake Champlain and Connecticut River sea lamprey populations in response to 0.3 mg/L of TFM with muscle tissue samples (GE 1; Table S2). logFC stands for log2-fold changes. Annotation names come from Swiss-Prot and Uniref90 via the Trinotate annotation protocol (http://trinotate.github.io).

| Trinity Gene | Population | logFC | Annotation |
|---|---|---|---|
| TRINITY_DN240949_c11_g3 | Lake Michigan | 6.6472 | |
| TRINITY_DN177977_c9_g1 | Lake Michigan | 5.4985 | |
| TRINITY_DN206830_c12_g5 | Lake Michigan | 4.1453 | KLF2 |
| TRINITY_DN172847_c2_g1 | Lake Michigan | 6.4501 | C163A,DMBT1 |
| TRINITY_DN164606_c0_g1 | Lake Michigan | 5.6634 | |
| TRINITY_DN234143_c2_g1 | Lake Michigan | -8.5510 | EF2 |
| TRINITY_DN246732_c2_g1 | Lake Michigan | 4.1212 | INPP |
| TRINITY_DN233991_c7_g1 | Lake Michigan | 3.6670 | ZNFX1 |
| TRINITY_DN130326_c1_g1 | Lake Michigan | -9.2629 | |
| TRINITY_DN236020_c8_g2 | Lake Michigan | 3.6986 | |
| TRINITY_DN222882_c9_g2 | Lake Michigan | 3.7836 | |
| TRINITY_DN100240_c0_g1 | Lake Michigan | 4.9082 | |
| TRINITY_DN232012_c7_g1 | Lake Michigan | 5.4586 | |
| TRINITY_DN243609_c8_g1 | Lake Michigan | 8.5512 | |
| TRINITY_DN199132_c5_g1 | Lake Michigan | 3.3210 | |
| TRINITY_DN242471_c6_g3 | Lake Michigan | 3.5786 | CRBG1 |
| TRINITY_DN165096_c4_g2 | Lake Michigan | 4.7667 | CTND2,PKP4 |
| TRINITY_DN202153_c2_g1 | Lake Michigan | 3.3023 | GFPT1,GFPT2 |
| TRINITY_DN217487_c10_g1 | Lake Michigan | 3.6717 | |
| TRINITY_DN241178_c4_g1 | Lake Michigan | -9.7161 | |
| TRINITY_DN242347_c3_g2 | Lake Michigan | 3.6683 | AHNK |
| TRINITY_DN190231_c5_g2 | Lake Michigan | 4.6850 | |
| TRINITY_DN246854_c5_g1 | Lake Michigan | 3.2945 | NIN |
| TRINITY_DN233649_c4_g2 | Lake Michigan | 4.0412 | PAR12,ZCCHV |
| TRINITY_DN234408_c7_g1 | Lake Michigan | -9.7006 | |
| TRINITY_DN199896_c9_g1 | Lake Michigan | 5.7901 | CUZD1 |
| TRINITY_DN179559_c6_g1 | Lake Michigan | 3.6727 | CTND2,PKP3 |
| TRINITY_DN238773_c0_g1 | Lake Michigan | 3.6197 | NEB1 |
| TRINITY_DN242233_c7_g1 | Lake Michigan | -3.6962 | |
| TRINITY_DN189725_c4_g1 | Lake Michigan | 3.2509 | VASN |
| TRINITY_DN223020_c5_g1 | Lake Michigan | 5.6473 | |
| TRINITY_DN232012_c1_g1 | Lake Michigan | 4.8111 | |
| TRINITY_DN245284_c0_g3 | Lake Michigan | -5.0671 | |
| TRINITY_DN242671_c1_g1 | Lake Michigan | 3.5630 | PLEC,MACF1,DYST |
| TRINITY_DN245355_c9_g1 | Lake Michigan | 9.0405 | |
| TRINITY_DN239774_c4_g3 | Lake Michigan | 2.8601 | |

| | | | |
|---|---|---|---|
| TRINITY_DN156208_c2_g1 | Lake Michigan | 2.8144 | |
| TRINITY_DN227876_c6_g2 | Lake Michigan | 3.2813 | AHNK |
| TRINITY_DN198801_c15_g1 | Lake Michigan | 5.4564 | |
| TRINITY_DN240021_c6_g3 | Lake Michigan | 3.9789 | |
| TRINITY_DN171179_c6_g3 | Lake Michigan | 2.7724 | |
| TRINITY_DN247697_c7_g2 | Lake Michigan | 2.9929 | TRI29 |
| TRINITY_DN197040_c0_g3 | Lake Michigan | -5.2436 | |
| TRINITY_DN188612_c14_g1 | Lake Michigan | 3.2263 | TRI25 |
| TRINITY_DN174894_c4_g1 | Lake Michigan | 2.8241 | |
| TRINITY_DN162028_c4_g1 | Lake Michigan | 8.2892 | |
| TRINITY_DN191439_c4_g1 | Lake Michigan | 3.3290 | |
| TRINITY_DN214124_c3_g1 | Lake Michigan | 3.2374 | AHNK,AHNK2,PRAX |
| TRINITY_DN204648_c10_g1 | Lake Michigan | -6.4718 | |
| TRINITY_DN214110_c5_g1 | Lake Michigan | 5.1636 | |
| TRINITY_DN243134_c7_g1 | Lake Michigan | 3.1749 | REL |
| TRINITY_DN230376_c14_g1 | Lake Michigan | 2.9403 | |
| TRINITY_DN144780_c0_g1 | Lake Michigan | 6.4940 | |
| TRINITY_DN179836_c0_g2 | Lake Michigan | 3.8661 | |
| TRINITY_DN178101_c1_g1 | Lake Michigan | 4.6022 | |
| TRINITY_DN178880_c9_g1 | Lake Michigan | 4.7562 | |
| TRINITY_DN247145_c7_g1 | Lake Michigan | 3.6244 | |
| TRINITY_DN163626_c7_g1 | Lake Michigan | 4.0747 | |
| TRINITY_DN194795_c5_g1 | Lake Michigan | 3.0962 | AT8B1 |
| TRINITY_DN246208_c2_g1 | Lake Michigan | 2.8892 | ITA1,ITA10,ITA11 |
| TRINITY_DN201033_c10_g1 | Lake Michigan | 2.5078 | |
| TRINITY_DN228840_c3_g4 | Lake Michigan | 3.4469 | DMBT1 |
| TRINITY_DN247358_c5_g3 | Lake Michigan | 3.0840 | EPS15 |
| TRINITY_DN242347_c2_g1 | Lake Michigan | 2.8023 | AHNK |
| TRINITY_DN193825_c11_g1 | Lake Michigan | 3.2484 | |
| TRINITY_DN199829_c9_g1 | Lake Michigan | 6.9289 | |
| TRINITY_DN226770_c5_g1 | Lake Michigan | 2.9659 | MAVS |
| TRINITY_DN232509_c0_g1 | Lake Michigan | 2.9550 | CO1A2,CO4A2,COHA1,BAG,OTO1A |
| TRINITY_DN169265_c7_g1 | Lake Michigan | -10.4249 | |
| TRINITY_DN222587_c5_g1 | Lake Michigan | 3.2235 | |
| TRINITY_DN172830_c5_g2 | Lake Michigan | 5.8540 | |
| TRINITY_DN186749_c13_g2 | Lake Michigan | 4.9290 | |
| TRINITY_DN238401_c8_g1 | Lake Michigan | 2.6037 | |
| TRINITY_DN222752_c157_g4 | Lake Michigan | -2.6338 | |
| TRINITY_DN170573_c15_g1 | Lake Michigan | 4.7892 | |
| TRINITY_DN249590_c3_g3 | Lake Michigan | -9.1349 | |
| TRINITY_DN212280_c3_g1 | Lake Michigan | 3.6077 | IP3KB |
| TRINITY_DN210369_c6_g3 | Lake Michigan | -3.6142 | |
| TRINITY_DN218001_c8_g1 | Lake Michigan | 2.2310 | |

| | | | |
|---|---|---|---|
| TRINITY_DN160168_c3_g1 | Lake Michigan | 10.8109 | |
| TRINITY_DN246813_c5_g2 | Lake Michigan | 3.1429 | TBFA |
| TRINITY_DN214124_c2_g1 | Lake Michigan | 2.6329 | AHNK |
| TRINITY_DN248720_c0_g4 | Lake Michigan | -5.6388 | COX5B |
| TRINITY_DN191403_c4_g3 | Lake Michigan | -9.2610 | |
| TRINITY_DN223093_c3_g1 | Lake Michigan | 3.4450 | |
| TRINITY_DN180979_c12_g1 | Lake Michigan | 3.1019 | CUTA,RL18 |
| TRINITY_DN168996_c7_g2 | Lake Michigan | 2.7766 | STON2 |
| TRINITY_DN237634_c8_g2 | Lake Michigan | 2.9212 | |
| TRINITY_DN217771_c5_g1 | Lake Michigan | 3.5733 | CASP9 |
| TRINITY_DN244457_c1_g3 | Lake Michigan | 3.0791 | ASTA |
| TRINITY_DN208086_c2_g2 | Lake Michigan | 2.8699 | RIPK1 |
| TRINITY_DN206384_c28_g1 | Lake Michigan | 2.5079 | |
| TRINITY_DN195594_c6_g4 | Lake Michigan | -3.8044 | |
| TRINITY_DN185295_c0_g1 | Lake Michigan | -3.7649 | OLFM |
| TRINITY_DN185233_c1_g1 | Lake Michigan | 3.1669 | PGS2 |
| TRINITY_DN198442_c9_g1 | Lake Michigan | 3.0876 | |
| TRINITY_DN243808_c3_g3 | Lake Michigan | 2.7328 | NCOR1,NCOR2 |
| TRINITY_DN236699_c14_g1 | Lake Michigan | 2.7666 | TRIM8 |
| TRINITY_DN169498_c15_g1 | Lake Michigan | 3.1044 | |
| TRINITY_DN246488_c9_g2 | Lake Michigan | 2.6594 | SRC8,HCLS1 |
| TRINITY_DN235370_c3_g1 | Lake Michigan | 5.3911 | COR2B |
| TRINITY_DN213424_c2_g2 | Lake Michigan | 3.0249 | SIK1,SIK2,SIK3 |
| TRINITY_DN189145_c1_g1 | Lake Michigan | 3.0835 | PSA,PSAL |
| TRINITY_DN235776_c2_g1 | Lake Michigan | 2.6748 | |
| TRINITY_DN223704_c3_g1 | Lake Michigan | 5.7403 | |
| TRINITY_DN244095_c24_g2 | Lake Michigan | 2.4647 | |
| TRINITY_DN173260_c11_g1 | Lake Michigan | 2.8546 | |
| TRINITY_DN175315_c1_g1 | Lake Michigan | 3.4524 | YPF08,MUC5A,MUC5B |
| TRINITY_DN145770_c0_g1 | Lake Michigan | -5.3871 | |
| TRINITY_DN186814_c12_g2 | Lake Michigan | 2.7094 | |
| TRINITY_DN198880_c9_g2 | Lake Michigan | -5.0340 | |
| TRINITY_DN246514_c4_g1 | Lake Michigan | 2.6795 | RASEF |
| TRINITY_DN202094_c6_g1 | Lake Michigan | 2.7652 | LYVE1 |
| TRINITY_DN194327_c6_g1 | Lake Michigan | -4.2239 | |
| TRINITY_DN211133_c0_g1 | Lake Michigan | -3.7568 | |
| TRINITY_DN246579_c3_g1 | Lake Michigan | 3.3688 | DIP2,DIP2A,DIP2B,DIP2C |
| TRINITY_DN154523_c0_g1 | Lake Michigan | 9.0696 | |
| TRINITY_DN216107_c3_g2 | Lake Michigan | 3.3848 | |
| TRINITY_DN169366_c10_g1 | Lake Michigan | -9.3384 | |
| TRINITY_DN226795_c23_g18 | Lake Michigan | 5.4043 | |
| TRINITY_DN233748_c1_g2 | Lake Michigan | 2.4108 | DDX58 |
| TRINITY_DN165555_c3_g1 | Lake Michigan | 7.4179 | |

| | | | |
|---|---|---|---|
| TRINITY_DN231305_c10_g1 | Lake Michigan | 2.7781 | |
| TRINITY_DN247290_c12_g3 | Lake Michigan | 3.2818 | AEP1 |
| TRINITY_DN192411_c10_g2 | Lake Michigan | 2.6000 | CRCM1 |
| TRINITY_DN238000_c1_g5 | Lake Michigan | -3.8480 | |
| TRINITY_DN220663_c7_g1 | Lake Michigan | 2.8159 | SPAS2 |
| TRINITY_DN229631_c1_g1 | Lake Michigan | 3.6332 | UROL1 |
| TRINITY_DN215572_c6_g4 | Lake Michigan | -9.5478 | |
| TRINITY_DN169825_c5_g4 | Lake Michigan | 3.3603 | CAV1 |
| TRINITY_DN177077_c4_g1 | Lake Michigan | 2.7627 | TLR2 |
| TRINITY_DN241903_c1_g4 | Lake Michigan | 3.0973 | |
| TRINITY_DN217493_c3_g1 | Lake Michigan | 2.5196 | KC1A |
| TRINITY_DN230937_c7_g1 | Lake Michigan | 2.8833 | CAN5 |
| TRINITY_DN226801_c3_g1 | Lake Michigan | -3.5210 | |
| TRINITY_DN244809_c1_g1 | Lake Michigan | 2.9870 | FRY,FRYL |
| TRINITY_DN235808_c5_g1 | Lake Michigan | 2.8604 | PEPL |
| TRINITY_DN244782_c3_g1 | Lake Michigan | 2.7049 | KIF5C,KINH |
| TRINITY_DN194368_c2_g3 | Lake Michigan | 9.6015 | |
| TRINITY_DN248570_c6_g1 | Lake Michigan | 4.7377 | |
| TRINITY_DN205658_c0_g1 | Lake Michigan | 4.2853 | CAD13 |
| TRINITY_DN217249_c3_g1 | Lake Michigan | 5.4118 | VLPB,FAR1 |
| TRINITY_DN223017_c5_g1 | Lake Michigan | 3.0445 | |
| TRINITY_DN201842_c8_g1 | Lake Michigan | 2.9851 | |
| TRINITY_DN177441_c11_g1 | Lake Michigan | -4.9290 | |
| TRINITY_DN200168_c5_g4 | Lake Michigan | 3.8631 | |
| TRINITY_DN247290_c4_g1 | Lake Michigan | 2.9591 | AEP1 |
| TRINITY_DN204487_c3_g3 | Lake Michigan | 2.9420 | FA83H |
| TRINITY_DN246238_c17_g1 | Lake Michigan | -3.5094 | |
| TRINITY_DN167885_c6_g1 | Lake Michigan | -7.2280 | |
| TRINITY_DN211018_c13_g1 | Lake Michigan | -9.4090 | |
| TRINITY_DN176921_c4_g2 | Lake Michigan | 2.7400 | PXDC2 |
| TRINITY_DN208776_c6_g2 | Lake Michigan | -2.5002 | |
| TRINITY_DN188295_c5_g1 | Lake Michigan | 2.8607 | |
| TRINITY_DN193254_c6_g1 | Lake Michigan | 3.1185 | CHST6 |
| TRINITY_DN212162_c5_g4 | Lake Michigan | -4.9077 | |
| TRINITY_DN197078_c13_g1 | Lake Michigan | 3.7921 | |
| TRINITY_DN164669_c8_g2 | Lake Michigan | 3.5461 | CAN2 |
| TRINITY_DN231225_c0_g1 | Lake Michigan | 2.7081 | CE350 |
| TRINITY_DN248486_c3_g1 | Lake Michigan | 3.5568 | IRS1A,IRS1B |
| TRINITY_DN190269_c9_g2 | Lake Michigan | 3.7079 | |
| TRINITY_DN1315_c0_g1 | Lake Michigan | 6.8672 | |
| TRINITY_DN223650_c5_g1 | Lake Michigan | 2.6753 | ZBT21,ZN628 |
| TRINITY_DN231661_c3_g1 | Lake Michigan | 2.4182 | IKZF2 |
| TRINITY_DN244667_c9_g1 | Lake Michigan | 3.1162 | |

| | | | |
|---|---|---|---|
| TRINITY_DN248641_c3_g5 | Lake Michigan | 2.5544 | ELMO2 |
| TRINITY_DN205784_c1_g1 | Lake Michigan | 2.6870 | AHNK |
| TRINITY_DN164430_c1_g1 | Lake Michigan | 3.3523 | FBN1,FBN3 |
| TRINITY_DN241905_c5_g1 | Lake Michigan | 3.4157 | |
| TRINITY_DN243350_c4_g3 | Lake Michigan | 3.0445 | |
| TRINITY_DN169600_c3_g1 | Lake Michigan | -3.3354 | |
| TRINITY_DN206206_c9_g1 | Lake Michigan | 3.0416 | TRI25,TRI29,TRI72,NF7B |
| TRINITY_DN231693_c11_g2 | Lake Michigan | 4.1524 | ANKR1 |
| TRINITY_DN172207_c15_g4 | Lake Michigan | 2.5457 | |
| TRINITY_DN168430_c6_g1 | Lake Michigan | 2.4647 | JUN |
| TRINITY_DN247227_c2_g1 | Lake Michigan | -5.9846 | HFM1 |
| TRINITY_DN199480_c6_g1 | Lake Michigan | 2.6526 | |
| TRINITY_DN236596_c2_g2 | Lake Michigan | 10.3174 | |
| TRINITY_DN237070_c2_g1 | Lake Michigan | 2.6136 | TM131 |
| TRINITY_DN232122_c4_g1 | Lake Michigan | 2.5873 | |
| TRINITY_DN210828_c2_g2 | Lake Michigan | 6.0371 | |
| TRINITY_DN220129_c3_g1 | Lake Michigan | -4.5858 | ZCPW1 |
| TRINITY_DN204395_c4_g12 | Lake Michigan | -3.8895 | |
| TRINITY_DN205101_c3_g1 | Lake Michigan | 3.3283 | |
| TRINITY_DN245438_c3_g2 | Lake Michigan | 2.8054 | AGRE1,FBP1,NOTC4,NOTCH,TENA,UPK3A |
| TRINITY_DN213140_c5_g2 | Lake Michigan | 2.8624 | |
| TRINITY_DN201937_c7_g3 | Lake Michigan | 3.0130 | |
| TRINITY_DN204288_c11_g2 | Lake Michigan | 2.8358 | |
| TRINITY_DN236246_c3_g1 | Lake Michigan | 2.4095 | PLCD4 |
| TRINITY_DN247346_c3_g2 | Lake Michigan | 3.0103 | ABCA3 |
| TRINITY_DN243612_c1_g1 | Lake Michigan | 2.5905 | HELZ2 |
| TRINITY_DN249209_c5_g2 | Lake Michigan | 3.3101 | C356 |
| TRINITY_DN197114_c2_g1 | Lake Michigan | -5.2487 | PODO |
| TRINITY_DN194437_c8_g1 | Lake Michigan | 5.3604 | |
| TRINITY_DN223582_c0_g1 | Lake Michigan | 2.4100 | |
| TRINITY_DN239596_c2_g1 | Lake Michigan | 2.8487 | F198A |
| TRINITY_DN186430_c3_g1 | Lake Michigan | -2.9732 | |
| TRINITY_DN206141_c10_g1 | Lake Michigan | 2.5411 | I17RA |
| TRINITY_DN236521_c6_g1 | Lake Michigan | 3.1327 | TRI62,TRI65 |
| TRINITY_DN245953_c4_g1 | Lake Michigan | 2.8183 | GRM2A,GRM2B |
| TRINITY_DN219420_c5_g1 | Lake Michigan | 2.8252 | |
| TRINITY_DN240942_c7_g1 | Lake Michigan | 2.6726 | AT8B1 |
| TRINITY_DN246089_c5_g1 | Lake Michigan | 2.7219 | SORL |
| TRINITY_DN236264_c1_g1 | Lake Michigan | 2.4895 | MOV10 |
| TRINITY_DN196367_c2_g1 | Lake Michigan | 3.3077 | M4K3 |
| TRINITY_DN242765_c3_g1 | Lake Michigan | 7.3034 | TCNA |
| TRINITY_DN169090_c8_g2 | Lake Michigan | 3.9739 | CEBPA |
| TRINITY_DN232771_c1_g1 | Lake Michigan | 2.3406 | SSFA2 |

| | | | |
|---|---|---|---|
| TRINITY_DN245719_c6_g1 | Lake Michigan | 2.9388 | ITB4 |
| TRINITY_DN190765_c1_g1 | Lake Michigan | 2.8377 | PPN |
| TRINITY_DN249590_c5_g1 | Lake Michigan | -4.6278 | |
| TRINITY_DN189803_c1_g1 | Lake Michigan | 3.3184 | SF3A2 |
| TRINITY_DN249710_c8_g1 | Lake Michigan | 2.9518 | |
| TRINITY_DN213538_c6_g1 | Lake Michigan | 2.5454 | |
| TRINITY_DN125322_c0_g1 | Lake Michigan | 5.4300 | |
| TRINITY_DN170044_c0_g2 | Lake Michigan | 2.8949 | AHNK |
| TRINITY_DN204851_c6_g1 | Lake Michigan | 2.3794 | |
| TRINITY_DN202943_c5_g1 | Lake Michigan | 2.5470 | PAR14,DTX3L |
| TRINITY_DN218550_c3_g1 | Lake Michigan | 3.0072 | FAT1 |
| TRINITY_DN223195_c3_g2 | Lake Michigan | 2.7619 | |
| TRINITY_DN246200_c8_g2 | Lake Michigan | 2.4894 | CEBPA,CEBPB |
| TRINITY_DN249319_c6_g1 | Lake Michigan | 4.0988 | APOB |
| TRINITY_DN244752_c3_g1 | Lake Michigan | 2.8051 | PKHG1,PKHG3 |
| TRINITY_DN200112_c6_g3 | Lake Michigan | -3.3851 | |
| TRINITY_DN181398_c5_g2 | Lake Michigan | 3.6721 | FKB1A,FKB1B |
| TRINITY_DN232786_c2_g2 | Lake Michigan | 2.5401 | PTN4 |
| TRINITY_DN197634_c5_g1 | Lake Michigan | 3.1920 | AT2C1 |
| TRINITY_DN231891_c2_g2 | Lake Michigan | 2.2672 | |
| TRINITY_DN210496_c2_g1 | Lake Michigan | 2.4958 | CASP3,CASP7 |
| TRINITY_DN235384_c0_g1 | Lake Michigan | -4.2707 | MARH4 |
| TRINITY_DN175768_c0_g1 | Lake Michigan | 3.1221 | GRN |
| TRINITY_DN239236_c15_g1 | Lake Michigan | 3.5436 | |
| TRINITY_DN189394_c4_g2 | Lake Michigan | 2.5609 | FOXN3 |
| TRINITY_DN172246_c3_g1 | Lake Michigan | 3.2610 | |
| TRINITY_DN219727_c2_g2 | Lake Michigan | 4.3372 | MOXD1,MOXD2 |
| TRINITY_DN246854_c5_g2 | Lake Michigan | 3.1272 | NIN |
| TRINITY_DN240436_c3_g1 | Lake Michigan | 2.7897 | ZNRF2 |
| TRINITY_DN225105_c9_g3 | Lake Michigan | 2.7592 | AT2A2,AT2A3 |
| TRINITY_DN171663_c7_g1 | Lake Michigan | 7.9424 | |
| TRINITY_DN245115_c2_g1 | Lake Michigan | 2.4466 | SHRM3 |
| TRINITY_DN175759_c4_g2 | Lake Michigan | 3.7388 | NAS36 |
| TRINITY_DN162219_c0_g1 | Lake Michigan | 5.5234 | |
| TRINITY_DN247511_c6_g1 | Lake Michigan | 9.6540 | |
| TRINITY_DN180126_c6_g3 | Lake Michigan | -4.6149 | |
| TRINITY_DN193878_c2_g1 | Lake Michigan | 2.3492 | TTPAL |
| TRINITY_DN222451_c1_g1 | Lake Michigan | -4.7685 | TO6BL |
| TRINITY_DN244469_c5_g1 | Lake Michigan | 2.9897 | ZSWM8 |
| TRINITY_DN184509_c12_g1 | Lake Michigan | -3.1791 | |
| TRINITY_DN190269_c9_g5 | Lake Michigan | 3.2027 | |
| TRINITY_DN188992_c3_g1 | Lake Michigan | 3.7946 | TRAK1 |
| TRINITY_DN212486_c1_g3 | Lake Michigan | -14.3348 | MIOX |

| | | | |
|---|---|---|---|
| TRINITY_DN177907_c2_g1 | Lake Michigan | 5.4829 | |
| TRINITY_DN248392_c8_g2 | Lake Michigan | 2.8233 | OTOF |
| TRINITY_DN221002_c0_g2 | Lake Michigan | -5.7759 | |
| TRINITY_DN237055_c13_g7 | Lake Michigan | 2.2884 | |
| TRINITY_DN220794_c3_g1 | Lake Michigan | -3.2377 | |
| TRINITY_DN242663_c7_g1 | Lake Michigan | 6.7926 | PRT1B |
| TRINITY_DN242255_c7_g1 | Lake Michigan | 2.2624 | RERG |
| TRINITY_DN166003_c4_g1 | Lake Michigan | 7.4609 | |
| TRINITY_DN244758_c1_g1 | Lake Michigan | 2.3790 | DC1L1,DC1L2 |
| TRINITY_DN187133_c3_g1 | Lake Michigan | -4.3035 | |
| TRINITY_DN228994_c7_g3 | Lake Michigan | -2.4598 | |
| TRINITY_DN234693_c6_g1 | Lake Michigan | 3.0195 | BORG4 |
| TRINITY_DN159483_c0_g1 | Lake Michigan | 5.5008 | |
| TRINITY_DN160209_c2_g1 | Lake Michigan | 6.4080 | |
| TRINITY_DN225177_c10_g1 | Lake Michigan | 2.6281 | |
| TRINITY_DN209096_c0_g1 | Lake Michigan | 2.3443 | |
| TRINITY_DN195312_c3_g2 | Lake Michigan | 3.9308 | FAT1 |
| TRINITY_DN195299_c0_g1 | Lake Michigan | 2.7121 | MLP3A |
| TRINITY_DN249432_c3_g1 | Lake Michigan | 2.9164 | AOXB,ALDO1,XDH |
| TRINITY_DN218935_c4_g2 | Lake Michigan | 3.0388 | |
| TRINITY_DN236502_c10_g1 | Lake Michigan | -2.2949 | |
| TRINITY_DN248130_c2_g3 | Lake Michigan | 2.6087 | PLEC |
| TRINITY_DN241984_c13_g2 | Lake Michigan | 4.1383 | MYH1 |
| TRINITY_DN225677_c6_g1 | Lake Michigan | 2.5897 | |
| TRINITY_DN206910_c4_g2 | Lake Michigan | 2.9692 | RBP2A |
| TRINITY_DN235488_c2_g1 | Lake Michigan | 3.6904 | |
| TRINITY_DN208488_c4_g1 | Lake Michigan | 2.0358 | PAK1 |
| TRINITY_DN176027_c5_g1 | Lake Michigan | -2.5882 | |
| TRINITY_DN201876_c2_g2 | Lake Michigan | 2.7946 | MXRA5,IGS10 |
| TRINITY_DN250334_c10_g1 | Lake Michigan | 2.4750 | MBOA2 |
| TRINITY_DN248664_c4_g2 | Lake Michigan | 2.6216 | PARP4 |
| TRINITY_DN241984_c16_g2 | Lake Michigan | 3.8963 | MYH7 |
| TRINITY_DN192484_c4_g6 | Lake Michigan | 2.6292 | NUMB |
| TRINITY_DN226859_c0_g1 | Lake Michigan | 4.4190 | MYH1,MYH3,MYH4,MYSS |
| TRINITY_DN181539_c5_g1 | Lake Michigan | -10.7403 | TKRA,GYAR |
| TRINITY_DN247520_c1_g4 | Lake Michigan | 2.6536 | NADK |
| TRINITY_DN180898_c4_g1 | Lake Michigan | 2.4495 | MIB2 |
| TRINITY_DN234464_c8_g1 | Lake Michigan | 2.2688 | |
| TRINITY_DN178855_c3_g3 | Lake Michigan | 2.6430 | EHF |
| TRINITY_DN205579_c4_g1 | Lake Michigan | 2.5826 | |
| TRINITY_DN246635_c11_g1 | Lake Michigan | -3.7522 | |
| TRINITY_DN248170_c10_g1 | Lake Michigan | 4.5125 | |
| TRINITY_DN239613_c16_g1 | Lake Michigan | 2.5517 | DYST |

| | | | |
|---|---|---|---|
| TRINITY_DN247906_c12_g1 | Lake Michigan | -2.6352 | |
| TRINITY_DN317500_c2_g1 | Lake Michigan | -2.9725 | |
| TRINITY_DN243490_c2_g2 | Lake Michigan | 2.5837 | MYO1E |
| TRINITY_DN241358_c3_g3 | Lake Michigan | 2.7568 | KDM6A,KDM6B,UTY |
| TRINITY_DN234256_c2_g1 | Lake Michigan | 3.5969 | MYH4,MYH6,MYH7 |
| TRINITY_DN210860_c6_g2 | Lake Michigan | 2.8747 | |
| TRINITY_DN231746_c7_g1 | Lake Michigan | 3.3042 | ANTR1 |
| TRINITY_DN243577_c2_g1 | Lake Michigan | 2.2420 | ECE,ECE1,ECE2,EFCE2,NEP,NEP4 |
| TRINITY_DN174047_c4_g2 | Lake Michigan | -5.0402 | |
| TRINITY_DN248359_c5_g3 | Lake Michigan | 2.4903 | EMP2,PMP22 |
| TRINITY_DN242911_c0_g3 | Lake Michigan | 2.4469 | TGO1 |
| TRINITY_DN215107_c3_g3 | Lake Michigan | 2.1418 | AHNK |
| TRINITY_DN161298_c10_g1 | Lake Michigan | 6.5791 | ENTK |
| TRINITY_DN317516_c0_g1 | Lake Michigan | -6.4479 | |
| TRINITY_DN163260_c7_g1 | Lake Michigan | 2.7141 | |
| TRINITY_DN201356_c4_g1 | Lake Michigan | 2.1614 | |
| TRINITY_DN233432_c6_g1 | Lake Michigan | 2.4054 | G6PT2,G6PT3 |
| TRINITY_DN213122_c0_g1 | Lake Michigan | -3.7441 | |
| TRINITY_DN216640_c4_g1 | Lake Michigan | -2.9215 | |
| TRINITY_DN240509_c9_g3 | Lake Michigan | 2.8183 | PDCD4 |
| TRINITY_DN198466_c8_g1 | Lake Michigan | 3.3152 | |
| TRINITY_DN211201_c10_g1 | Lake Michigan | 3.0919 | |
| TRINITY_DN218187_c1_g1 | Lake Michigan | 2.8900 | CAH2,CAH7,CAHZ |
| TRINITY_DN246696_c4_g2 | Lake Michigan | -2.4316 | |
| TRINITY_DN170237_c16_g1 | Lake Michigan | 2.8642 | SDHF4 |
| TRINITY_DN159045_c5_g1 | Lake Michigan | 8.6272 | |
| TRINITY_DN215107_c3_g2 | Lake Michigan | 2.9570 | AHNK |
| TRINITY_DN193373_c6_g1 | Lake Michigan | 2.3603 | RN217 |
| TRINITY_DN238183_c6_g1 | Lake Michigan | 2.4302 | PK3CB,PK3CD |
| TRINITY_DN166363_c0_g4 | Lake Michigan | 3.4550 | |
| TRINITY_DN208682_c3_g2 | Lake Michigan | 2.9349 | GBP |
| TRINITY_DN241984_c13_g3 | Lake Michigan | 5.2262 | MYH1 |
| TRINITY_DN248248_c3_g1 | Lake Michigan | 2.1301 | SON |
| TRINITY_DN195019_c2_g1 | Lake Michigan | 2.6280 | ARVC,CTND1 |
| TRINITY_DN249716_c13_g2 | Lake Michigan | 3.8729 | STK24,STK25 |
| TRINITY_DN234328_c3_g4 | Lake Michigan | 4.3291 | EAA3 |
| TRINITY_DN203841_c16_g1 | Lake Michigan | -3.5137 | |
| TRINITY_DN137150_c1_g1 | Lake Michigan | 3.4785 | |
| TRINITY_DN192900_c3_g2 | Lake Michigan | -2.6105 | |
| TRINITY_DN235084_c0_g1 | Lake Michigan | 3.0610 | |
| TRINITY_DN161209_c10_g1 | Lake Michigan | 2.9893 | ARHG8,ARHG3 |
| TRINITY_DN204201_c6_g1 | Lake Michigan | -2.3721 | |
| TRINITY_DN250403_c1_g1 | Lake Champlain | -7.7360 | |

| | | | |
|---|---|---|---|
| TRINITY_DN225692_c1_g2 | Lake Champlain | -6.9509 | |
| TRINITY_DN221620_c6_g1 | Lake Champlain | -8.6389 | |
| TRINITY_DN170451_c2_g1 | Lake Champlain | -5.8012 | |
| TRINITY_DN192514_c8_g2 | Lake Champlain | 10.0785 | |
| TRINITY_DN160374_c7_g1 | Lake Champlain | -6.3593 | UBIQP,RL403,RL40 |
| TRINITY_DN248570_c6_g1 | Lake Champlain | -9.0480 | |
| TRINITY_DN213658_c3_g1 | Lake Champlain | -4.1294 | |
| TRINITY_DN203610_c97_g4 | Lake Champlain | 3.8937 | |
| TRINITY_DN158767_c0_g1 | Lake Champlain | -3.7500 | |
| TRINITY_DN156381_c2_g1 | Lake Champlain | 8.1341 | UBIQP |
| TRINITY_DN249845_c10_g1 | Lake Champlain | -11.0569 | |
| TRINITY_DN172246_c3_g1 | Lake Champlain | -7.0018 | |
| TRINITY_DN236448_c6_g1 | Lake Champlain | -6.7912 | |
| TRINITY_DN196516_c7_g1 | Lake Champlain | -4.7207 | |
| TRINITY_DN157312_c10_g1 | Lake Champlain | -11.0561 | |
| TRINITY_DN203809_c17_g1 | Lake Champlain | -3.4992 | ADT3 |
| TRINITY_DN213644_c1_g2 | Lake Champlain | -7.5112 | |
| TRINITY_DN209792_c2_g1 | Lake Champlain | 10.6691 | UBIQP |
| TRINITY_DN163694_c3_g1 | Lake Champlain | 10.8108 | TYB10 |
| TRINITY_DN175759_c4_g2 | Lake Champlain | -6.5867 | NAS36 |
| TRINITY_DN221029_c7_g4 | Connecticut River | 10.7799 | |
| TRINITY_DN167794_c0_g1 | Connecticut River | 10.5248 | |
| TRINITY_DN238272_c11_g4 | Connecticut River | 7.9519 | |
| TRINITY_DN178227_c7_g1 | Connecticut River | 7.6062 | |
| TRINITY_DN242696_c19_g1 | Connecticut River | 6.4255 | |
| TRINITY_DN195091_c9_g2 | Connecticut River | 7.8064 | DYRK4 |
| TRINITY_DN236507_c5_g1 | Connecticut River | -7.5183 | RS27A |
| TRINITY_DN199829_c9_g1 | Connecticut River | 9.0175 | |
| TRINITY_DN234143_c2_g1 | Connecticut River | 6.0424 | EF2 |
| TRINITY_DN157312_c10_g1 | Connecticut River | -13.5094 | |
| TRINITY_DN233708_c7_g4 | Connecticut River | 7.2146 | |
| TRINITY_DN199164_c1_g1 | Connecticut River | 6.1549 | RS12 |
| TRINITY_DN237137_c5_g1 | Connecticut River | -7.0019 | CLCA1,CLCA4,CA3A1 |
| TRINITY_DN245355_c9_g1 | Connecticut River | 8.9792 | |
| TRINITY_DN196018_c18_g1 | Connecticut River | -5.8369 | |
| TRINITY_DN192514_c8_g2 | Connecticut River | -6.0804 | |
| TRINITY_DN177977_c17_g1 | Connecticut River | 12.5344 | |
| TRINITY_DN249845_c10_g1 | Connecticut River | -10.9864 | |
| TRINITY_DN209644_c5_g1 | Connecticut River | 7.9208 | |
| TRINITY_DN247013_c6_g1 | Connecticut River | 3.2837 | ALF2 |
| TRINITY_DN240998_c14_g1 | Connecticut River | 8.7969 | |
| TRINITY_DN186532_c5_g1 | Connecticut River | -4.8473 | |
| TRINITY_DN247918_c2_g1 | Connecticut River | 3.7320 | |

| | | | |
|---|---|---|---|
| TRINITY_DN215498_c1_g1 | Connecticut River | 9.2126 | |
| TRINITY_DN169125_c0_g3 | Connecticut River | 4.6800 | ATPB |
| TRINITY_DN247586_c5_g1 | Connecticut River | 8.9986 | |
| TRINITY_DN175721_c13_g1 | Connecticut River | 7.2903 | |
| TRINITY_DN245968_c8_g4 | Connecticut River | 4.9357 | |
| TRINITY_DN237686_c2_g4 | Connecticut River | 7.1499 | CXG1,CXG2 |
| TRINITY_DN250638_c20_g2 | Connecticut River | -8.1435 | |
| TRINITY_DN245882_c8_g1 | Connecticut River | -9.5951 | FRIH |
| TRINITY_DN162967_c2_g2 | Connecticut River | -11.4664 | CD109 |
| TRINITY_DN177269_c6_g1 | Connecticut River | 5.8791 | |
| TRINITY_DN234209_c3_g1 | Connecticut River | 4.5785 | |
| TRINITY_DN176914_c4_g1 | Connecticut River | 5.8234 | |
| TRINITY_DN165094_c7_g1 | Connecticut River | -5.9834 | |
| TRINITY_DN243862_c14_g1 | Connecticut River | 5.4977 | |
| TRINITY_DN168165_c3_g1 | Connecticut River | -12.4183 | |
| TRINITY_DN232267_c5_g3 | Connecticut River | -4.8353 | |
| TRINITY_DN225469_c1_g1 | Connecticut River | 6.5367 | NRCAM,L1CAM,NFASC |
| TRINITY_DN179044_c45_g2 | Connecticut River | 4.8484 | |
| TRINITY_DN204102_c1_g1 | Connecticut River | 5.1605 | |
| TRINITY_DN237686_c2_g2 | Connecticut River | 6.2525 | CXG1 |
| TRINITY_DN184677_c4_g1 | Connecticut River | -9.9330 | |
| TRINITY_DN159045_c5_g1 | Connecticut River | 10.6895 | |
| TRINITY_DN157620_c0_g1 | Connecticut River | 7.7929 | |
| TRINITY_DN210098_c3_g1 | Connecticut River | 6.3224 | |
| TRINITY_DN239082_c2_g1 | Connecticut River | 6.1887 | CXG1 |
| TRINITY_DN178070_c9_g3 | Connecticut River | 7.8262 | |
| TRINITY_DN249486_c8_g3 | Connecticut River | 5.1282 | |
| TRINITY_DN228258_c10_g1 | Connecticut River | 4.5059 | |
| TRINITY_DN200770_c3_g3 | Connecticut River | -5.1589 | |
| TRINITY_DN245659_c9_g1 | Connecticut River | -9.9673 | |
| TRINITY_DN194490_c4_g3 | Connecticut River | 8.8927 | COLA1 |
| TRINITY_DN217726_c2_g1 | Connecticut River | 7.2022 | CATA |
| TRINITY_DN170673_c10_g1 | Connecticut River | 6.2682 | |
| TRINITY_DN195091_c9_g5 | Connecticut River | 4.6501 | |
| TRINITY_DN196639_c7_g1 | Connecticut River | -5.0047 | |
| TRINITY_DN187825_c15_g5 | Connecticut River | 9.5782 | BGBP |
| TRINITY_DN238312_c4_g1 | Connecticut River | 4.4557 | ZN180,ZN391,ZN483,ZN544,ZNF79 |
| TRINITY_DN159951_c6_g1 | Connecticut River | 5.1105 | |
| TRINITY_DN244174_c4_g1 | Connecticut River | 4.8628 | PTK7,NFASC |
| TRINITY_DN227320_c3_g1 | Connecticut River | 4.9592 | |
| TRINITY_DN250665_c0_g1 | Connecticut River | 2.7430 | |
| TRINITY_DN244186_c3_g3 | Connecticut River | 4.7889 | |
| TRINITY_DN194953_c2_g1 | Connecticut River | -3.4333 | TBA1 |

| | | |
|---|---|---|
| TRINITY_DN167215_c5_g1 | Connecticut River | -4.8980 |
| TRINITY_DN246847_c1_g3 | Connecticut River | 4.2552 |

**Table S5.** Differentially Expressed Genes Detected in Lake Michigan, Lake Champlain and Connecticut River sea lamprey populations in response to 0.2 mg/L of TFM with liver tissue samples (GE 1; Table S2). logFC stands for log2-fold changes. Annotation names come from Swiss-Prot and Uniref90 via the Trinotate annotation protocol (http://trinotate.github.io).

| Trinity Gene | Population | logFC | Annotation |
|---|---|---|---|
| TRINITY_DN234143_c2_g1 | Lake Michigan | -7.1550 | EF2 |
| TRINITY_DN240949_c11_g3 | Lake Michigan | 6.3922 | |
| TRINITY_DN198801_c15_g1 | Lake Michigan | 5.3466 | |
| TRINITY_DN230856_c44_g6 | Lake Michigan | -8.6823 | |
| TRINITY_DN206515_c3_g1 | Lake Michigan | -11.0362 | |
| TRINITY_DN159831_c7_g4 | Lake Michigan | -2.8478 | |
| TRINITY_DN246651_c7_g1 | Lake Michigan | -3.2557 | |
| TRINITY_DN135565_c0_g1 | Lake Michigan | 5.3717 | |
| TRINITY_DN163926_c5_g1 | Lake Michigan | -3.2305 | |
| TRINITY_DN230856_c40_g10 | Lake Michigan | 11.4044 | |
| TRINITY_DN237651_c24_g4 | Lake Michigan | 4.5892 | |
| TRINITY_DN183168_c22_g1 | Lake Michigan | -3.4351 | |
| TRINITY_DN138776_c1_g1 | Lake Michigan | 7.4653 | |
| TRINITY_DN222483_c7_g1 | Lake Michigan | 6.7205 | Y7014 |
| TRINITY_DN196190_c22_g5 | Lake Michigan | -4.2587 | |
| TRINITY_DN241319_c7_g2 | Lake Michigan | -1.9420 | |
| TRINITY_DN100240_c0_g1 | Lake Michigan | 5.4648 | |
| TRINITY_DN210784_c14_g1 | Lake Michigan | -2.5527 | |
| TRINITY_DN245280_c10_g1 | Lake Michigan | 2.2728 | RS2 |
| TRINITY_DN169366_c10_g1 | Lake Michigan | -8.3678 | |
| TRINITY_DN213049_c1_g2 | Lake Michigan | -6.4300 | |
| TRINITY_DN216060_c7_g1 | Lake Michigan | -2.3955 | |
| TRINITY_DN189104_c17_g1 | Lake Michigan | 2.3426 | FIBA2 |
| TRINITY_DN171851_c7_g3 | Lake Michigan | -2.5833 | |
| TRINITY_DN221715_c7_g1 | Lake Michigan | 5.7968 | CNTN4,CNTN3,CNTN5 |
| TRINITY_DN225045_c15_g3 | Lake Michigan | -7.7290 | DNAS1 |
| TRINITY_DN203821_c6_g1 | Lake Michigan | -6.6935 | |
| TRINITY_DN198672_c29_g1 | Lake Michigan | -5.5237 | |
| TRINITY_DN144780_c0_g1 | Lake Michigan | 6.2329 | |
| TRINITY_DN184663_c9_g1 | Lake Michigan | -8.4301 | |
| TRINITY_DN90688_c0_g1 | Lake Michigan | 7.2919 | |
| TRINITY_DN159186_c5_g1 | Lake Michigan | -7.1924 | |
| TRINITY_DN167629_c6_g1 | Lake Michigan | -2.9758 | |
| TRINITY_DN160372_c9_g1 | Lake Michigan | -6.0035 | |
| TRINITY_DN242351_c12_g1 | Lake Michigan | 1.7785 | RS2 |
| TRINITY_DN236596_c2_g2 | Lake Michigan | 9.5630 | |

| | | | |
|---|---|---|---|
| TRINITY_DN214417_c0_g1 | Lake Michigan | 5.9551 | |
| TRINITY_DN180979_c12_g1 | Lake Michigan | 2.3393 | CUTA,RL18 |
| TRINITY_DN243239_c15_g1 | Lake Michigan | 2.4872 | VDAC2 |
| TRINITY_DN231658_c4_g1 | Lake Michigan | -2.9530 | |
| TRINITY_DN179019_c4_g3 | Lake Michigan | -2.6487 | |
| TRINITY_DN181648_c21_g1 | Lake Michigan | -3.0485 | |
| TRINITY_DN249546_c9_g1 | Lake Michigan | -2.0601 | TBA |
| TRINITY_DN203301_c3_g2 | Lake Michigan | -6.0221 | |
| TRINITY_DN119668_c0_g1 | Lake Michigan | -2.7739 | |
| TRINITY_DN217249_c3_g1 | Lake Michigan | 4.5799 | VLPB,FAR1 |
| TRINITY_DN204015_c16_g1 | Lake Michigan | -8.3861 | FIBA2 |
| TRINITY_DN162191_c7_g1 | Lake Michigan | -2.9931 | |
| TRINITY_DN243148_c1_g1 | Lake Michigan | -4.3097 | |
| TRINITY_DN166590_c2_g1 | Lake Michigan | -2.9671 | |
| TRINITY_DN236553_c3_g1 | Lake Michigan | -1.9856 | |
| TRINITY_DN234836_c1_g1 | Lake Michigan | -6.2470 | |
| TRINITY_DN221859_c6_g3 | Lake Michigan | -3.0344 | |
| TRINITY_DN166259_c8_g2 | Lake Michigan | -2.6438 | |
| TRINITY_DN82306_c0_g1 | Lake Michigan | 5.0414 | |
| TRINITY_DN183699_c8_g2 | Lake Michigan | -4.1530 | |
| TRINITY_DN242107_c3_g4 | Lake Michigan | -2.6089 | |
| TRINITY_DN236027_c1_g1 | Lake Michigan | -5.4067 | |
| TRINITY_DN249854_c12_g3 | Lake Michigan | -2.2838 | |
| TRINITY_DN202715_c9_g2 | Lake Michigan | -1.9576 | |
| TRINITY_DN222880_c3_g1 | Lake Champlain | -9.9363 | |
| TRINITY_DN247155_c0_g1 | Lake Champlain | -12.5111 | |
| TRINITY_DN188246_c6_g1 | Lake Champlain | -7.8660 | |
| TRINITY_DN198635_c4_g1 | Lake Champlain | 3.1962 | |
| TRINITY_DN248120_c1_g1 | Lake Champlain | -5.9794 | |
| TRINITY_DN200281_c2_g1 | Lake Champlain | -3.0653 | |
| TRINITY_DN245524_c2_g1 | Lake Champlain | -4.5627 | |
| TRINITY_DN246732_c2_g1 | Lake Champlain | 3.3747 | INPP |
| TRINITY_DN220830_c9_g2 | Lake Champlain | -2.8818 | |
| TRINITY_DN171843_c9_g1 | Lake Champlain | 9.6133 | |
| TRINITY_DN176006_c9_g1 | Lake Champlain | -10.2468 | |
| TRINITY_DN156208_c2_g1 | Lake Champlain | -2.7979 | |
| TRINITY_DN221051_c3_g1 | Lake Champlain | -6.7063 | |
| TRINITY_DN216887_c2_g1 | Lake Champlain | 4.9455 | |
| TRINITY_DN204022_c7_g1 | Lake Champlain | 3.3749 | |
| TRINITY_DN237180_c5_g2 | Lake Champlain | 2.4800 | FRIHB |
| TRINITY_DN137947_c0_g1 | Lake Champlain | -3.4874 | |
| TRINITY_DN183682_c10_g3 | Lake Champlain | -4.3898 | |
| TRINITY_DN243237_c0_g1 | Lake Champlain | 5.2604 | |

| | | | |
|---|---|---|---|
| TRINITY_DN217487_c10_g1 | Lake Champlain | 3.5887 | |
| TRINITY_DN237485_c8_g1 | Lake Champlain | 9.2743 | |
| TRINITY_DN190615_c10_g4 | Lake Champlain | -2.7171 | RSSA |
| TRINITY_DN248304_c20_g1 | Lake Champlain | -3.3708 | |
| TRINITY_DN125551_c0_g1 | Lake Champlain | -11.1659 | |
| TRINITY_DN205631_c5_g3 | Lake Champlain | -7.6946 | |
| TRINITY_DN230733_c3_g3 | Lake Champlain | 5.9222 | |
| TRINITY_DN208585_c4_g1 | Lake Champlain | -3.7068 | TCB1,TC1A |
| TRINITY_DN236020_c8_g2 | Lake Champlain | 2.9796 | |
| TRINITY_DN249096_c3_g1 | Lake Champlain | -3.1601 | |
| TRINITY_DN244915_c27_g2 | Lake Champlain | -2.3141 | |
| TRINITY_DN218796_c5_g2 | Lake Champlain | -7.2084 | |
| TRINITY_DN232192_c2_g1 | Lake Champlain | -4.1788 | |
| TRINITY_DN249569_c14_g1 | Lake Champlain | 4.3418 | PEAMT |
| TRINITY_DN223153_c1_g1 | Lake Champlain | -9.5596 | FABPH |
| TRINITY_DN232321_c19_g1 | Lake Champlain | 9.1082 | |
| TRINITY_DN188246_c6_g2 | Lake Champlain | -2.4513 | |
| TRINITY_DN240404_c2_g9 | Lake Champlain | 3.2274 | RL19 |
| TRINITY_DN156381_c2_g1 | Lake Champlain | 4.5021 | UBIQP |
| TRINITY_DN229062_c6_g2 | Lake Champlain | -8.1068 | |
| TRINITY_DN188247_c8_g2 | Lake Champlain | -2.2462 | |
| TRINITY_DN243035_c46_g1 | Lake Champlain | 4.2403 | |
| TRINITY_DN248931_c24_g6 | Connecticut River | 9.9120 | |
| TRINITY_DN184663_c5_g1 | Connecticut River | 12.2385 | |
| TRINITY_DN217408_c4_g3 | Connecticut River | -10.0808 | |
| TRINITY_DN234143_c2_g1 | Connecticut River | 6.7351 | EF2 |
| TRINITY_DN20313_c0_g1 | Connecticut River | -4.8457 | |
| TRINITY_DN185982_c4_g2 | Connecticut River | -13.1648 | SAM9L |
| TRINITY_DN242545_c10_g9 | Connecticut River | -5.3988 | |
| TRINITY_DN163898_c1_g1 | Connecticut River | 4.1598 | |
| TRINITY_DN229918_c3_g1 | Connecticut River | -12.0380 | NED4L,NEDD4,HCE1,NAS14 |
| TRINITY_DN107684_c0_g1 | Connecticut River | -4.4516 | |
| TRINITY_DN243862_c14_g1 | Connecticut River | 4.9513 | |
| TRINITY_DN203584_c0_g2 | Connecticut River | -6.4162 | |
| TRINITY_DN210361_c13_g1 | Connecticut River | 5.6139 | |
| TRINITY_DN184899_c6_g2 | Connecticut River | -10.4726 | |
| TRINITY_DN227577_c12_g5 | Connecticut River | -6.2660 | |
| TRINITY_DN198672_c13_g13 | Connecticut River | -7.8868 | |
| TRINITY_DN245838_c1_g1 | Connecticut River | 6.7886 | |
| TRINITY_DN215548_c14_g1 | Connecticut River | -6.5771 | |
| TRINITY_DN247200_c6_g1 | Connecticut River | -5.5937 | |
| TRINITY_DN249700_c5_g1 | Connecticut River | -9.6804 | SAM9L |
| TRINITY_DN156279_c6_g1 | Connecticut River | -8.5208 | |

| | | | |
|---|---|---|---|
| TRINITY_DN218859_c11_g1 | Connecticut River | -4.2974 | |
| TRINITY_DN243279_c3_g1 | Connecticut River | 3.5257 | |
| TRINITY_DN208430_c7_g1 | Connecticut River | -5.7771 | |
| TRINITY_DN221075_c2_g4 | Connecticut River | 3.6102 | EPYC |
| TRINITY_DN229902_c5_g1 | Connecticut River | -4.9651 | |
| TRINITY_DN158793_c93_g12 | Connecticut River | 4.4559 | APL2 |
| TRINITY_DN187972_c5_g1 | Connecticut River | 10.2795 | |
| TRINITY_DN233673_c6_g1 | Connecticut River | -4.6165 | |
| TRINITY_DN249782_c16_g1 | Connecticut River | 3.3873 | |
| TRINITY_DN164378_c19_g1 | Connecticut River | -4.6778 | |
| TRINITY_DN168334_c12_g1 | Connecticut River | -2.4032 | |
| TRINITY_DN175743_c7_g1 | Connecticut River | 2.7447 | |
| TRINITY_DN175676_c1_g1 | Connecticut River | -8.9055 | |
| TRINITY_DN250642_c0_g1 | Connecticut River | 2.4648 | |
| TRINITY_DN246606_c3_g1 | Connecticut River | 3.2491 | |
| TRINITY_DN242310_c7_g1 | Connecticut River | -2.3377 | |
| TRINITY_DN239484_c6_g1 | Connecticut River | -3.0369 | |
| TRINITY_DN242572_c6_g1 | Connecticut River | 2.1783 | |
| TRINITY_DN245962_c6_g2 | Connecticut River | -2.3010 | DPEP1 |
| TRINITY_DN221051_c3_g1 | Connecticut River | -5.7588 | |
| TRINITY_DN196190_c22_g7 | Connecticut River | -2.6468 | |
| TRINITY_DN224821_c3_g1 | Connecticut River | 2.3220 | KLH26 |
| TRINITY_DN236555_c6_g4 | Connecticut River | -2.9950 | ST2B1,ST1A1 |
| TRINITY_DN249712_c4_g2 | Connecticut River | -2.4399 | |
| TRINITY_DN234667_c4_g1 | Connecticut River | -1.9258 | |
| TRINITY_DN243035_c46_g1 | Connecticut River | 4.1229 | |
| TRINITY_DN249140_c7_g3 | Connecticut River | 2.6035 | |
| TRINITY_DN230333_c2_g1 | Connecticut River | -2.9252 | |
| TRINITY_DN209623_c8_g2 | Connecticut River | 4.3666 | A3LT2,GGTA1 |
| TRINITY_DN193585_c4_g1 | Connecticut River | 2.2375 | CLHC1 |
| TRINITY_DN122795_c0_g1 | Connecticut River | -9.3033 | |
| TRINITY_DN237533_c2_g1 | Connecticut River | 4.6746 | |
| TRINITY_DN250469_c0_g1 | Connecticut River | -4.9372 | |
| TRINITY_DN215385_c12_g1 | Connecticut River | 4.3747 | |
| TRINITY_DN225739_c21_g1 | Connecticut River | 2.1383 | |
| TRINITY_DN158793_c59_g1 | Connecticut River | -3.3272 | |
| TRINITY_DN191292_c2_g5 | Connecticut River | -3.7885 | |
| TRINITY_DN241396_c0_g1 | Connecticut River | -2.6043 | |
| TRINITY_DN250834_c0_g1 | Connecticut River | -2.0615 | |
| TRINITY_DN248931_c24_g7 | Connecticut River | 2.7783 | SEPP1 |
| TRINITY_DN169750_c6_g1 | Connecticut River | 5.1214 | |
| TRINITY_DN207481_c26_g1 | Connecticut River | -3.7116 | |
| TRINITY_DN159375_c1_g1 | Connecticut River | 1.8808 | |

| | | | |
|---|---|---|---|
| TRINITY_DN203543_c1_g1 | Connecticut River | -4.8164 | |
| TRINITY_DN175950_c9_g1 | Connecticut River | -2.3420 | |
| TRINITY_DN145785_c0_g1 | Connecticut River | 3.7401 | |
| TRINITY_DN182006_c13_g1 | Connecticut River | -2.9704 | |
| TRINITY_DN187910_c10_g2 | Connecticut River | 3.5354 | |
| TRINITY_DN248347_c4_g1 | Connecticut River | 10.1039 | |
| TRINITY_DN237912_c4_g1 | Connecticut River | -2.1674 | JPH3 |
| TRINITY_DN205201_c2_g1 | Connecticut River | -8.7677 | HUNIN |
| TRINITY_DN247649_c0_g1 | Connecticut River | -4.4673 | |
| TRINITY_DN194085_c10_g3 | Connecticut River | -3.3503 | |
| TRINITY_DN158793_c93_g16 | Connecticut River | -2.6084 | |
| TRINITY_DN38291_c2_g1 | Connecticut River | -2.1098 | |
| TRINITY_DN213973_c5_g1 | Connecticut River | 3.7251 | |
| TRINITY_DN126485_c0_g1 | Connecticut River | -4.1169 | |
| TRINITY_DN178077_c2_g1 | Connecticut River | 4.4647 | TMPS9,OVCH2,TM11D,PRS40,PCOC1,NRP2 |
| TRINITY_DN242545_c11_g3 | Connecticut River | 5.0125 | |
| TRINITY_DN250253_c20_g1 | Connecticut River | -4.1678 | |
| TRINITY_DN171335_c2_g1 | Connecticut River | 4.8807 | |
| TRINITY_DN243763_c1_g1 | Connecticut River | 9.3323 | |
| TRINITY_DN164938_c2_g1 | Connecticut River | 8.5205 | |
| TRINITY_DN188477_c3_g2 | Connecticut River | 7.3703 | |
| TRINITY_DN18277_c1_g1 | Connecticut River | 2.4035 | |
| TRINITY_DN245573_c2_g1 | Connecticut River | 9.8995 | |
| TRINITY_DN176470_c1_g2 | Connecticut River | -7.4435 | |
| TRINITY_DN206515_c2_g1 | Connecticut River | -3.7981 | |
| TRINITY_DN161615_c10_g1 | Connecticut River | 3.8387 | |
| TRINITY_DN240656_c3_g1 | Connecticut River | -3.0591 | CAD23 |
| TRINITY_DN226795_c23_g8 | Connecticut River | -8.0642 | |
| TRINITY_DN226506_c9_g7 | Connecticut River | -2.4415 | NDUV1 |
| TRINITY_DN212806_c2_g3 | Connecticut River | 10.3671 | |
| TRINITY_DN159571_c0_g1 | Connecticut River | -1.8843 | |
| TRINITY_DN191142_c3_g1 | Connecticut River | 5.2815 | |
| TRINITY_DN196256_c6_g1 | Connecticut River | 6.8999 | |
| TRINITY_DN199869_c3_g3 | Connecticut River | 1.9299 | ANO4 |
| TRINITY_DN228239_c1_g1 | Connecticut River | 2.4116 | |

**Table S6.** Differentially Expressed Genes Detected in Lake Michigan sea lamprey population in response to 0.2 mg/L of TFM with brain tissue samples (GE 1; Table S2). logFC stands for log2-fold changes. Annotation names come from Swiss-Prot and Uniref90 via the Trinotate annotation protocol (http://trinotate.github.io).

| Trinity Gene | Population | logFC | Annotation |
|---|---|---|---|
| TRINITY_DN138776_c1_g1 | Lake Michigan | 8.5935 | |
| TRINITY_DN248346_c1_g1 | Lake Michigan | 5.8879 | POL2,POL |
| TRINITY_DN135565_c0_g1 | Lake Michigan | 5.4862 | |
| TRINITY_DN249591_c0_g1 | Lake Michigan | 3.6490 | |
| TRINITY_DN247669_c1_g1 | Lake Michigan | 2.8062 | |
| TRINITY_DN180979_c12_g1 | Lake Michigan | 2.1859 | CUTA,RL18 |
| TRINITY_DN218372_c10_g1 | Lake Michigan | 1.8765 | |
| TRINITY_DN215307_c8_g1 | Lake Michigan | 1.4713 | SOCS3,CISH |
| TRINITY_DN119046_c0_g1 | Lake Michigan | -1.4266 | |
| TRINITY_DN249546_c9_g1 | Lake Michigan | -1.7890 | TBA |
| TRINITY_DN229777_c3_g1 | Lake Michigan | -1.8026 | |
| TRINITY_DN209469_c0_g4 | Lake Michigan | -2.4799 | |
| TRINITY_DN227073_c3_g2 | Lake Michigan | -2.7393 | |
| TRINITY_DN243148_c1_g1 | Lake Michigan | -3.9381 | |
| TRINITY_DN169366_c10_g1 | Lake Michigan | -4.6916 | |
| TRINITY_DN160372_c9_g1 | Lake Michigan | -6.1170 | |
| TRINITY_DN244915_c27_g1 | Lake Michigan | -6.6869 | |
| TRINITY_DN151112_c0_g1 | Lake Michigan | -7.6199 | |
| TRINITY_DN200770_c3_g3 | Lake Michigan | -7.7286 | |
| TRINITY_DN213049_c1_g2 | Lake Michigan | -8.3103 | |

**Table S7.** Differentially Expressed Genes Detected in Lake Michigan, Lake Champlain and Connecticut River sea lamprey populations in response to 0.2 mg/L of TFM with muscle tissue samples (GE 2; Table S2). logFC stands for log2-fold changes. Annotation names come from Swiss-Prot and Uniref90 via the Trinotate annotation protocol (http://trinotate.github.io).

| Trinity Gene | Population | logFC | Annotation |
|---|---|---|---|
| TRINITY_DN138776_c1_g1 | Lake Michigan | 8.3264 | |
| TRINITY_DN217249_c3_g1 | Lake Michigan | 5.0503 | VLPB,FAR1 |
| TRINITY_DN248346_c1_g1 | Lake Michigan | 4.3331 | POL,POL2 |
| TRINITY_DN250022_c6_g1 | Lake Michigan | -3.5009 | |
| TRINITY_DN198801_c15_g1 | Lake Michigan | 4.2165 | |
| TRINITY_DN177907_c2_g1 | Lake Michigan | 4.5902 | |
| TRINITY_DN135565_c0_g1 | Lake Michigan | 4.2306 | |
| TRINITY_DN225382_c4_g1 | Lake Michigan | -8.1051 | |
| TRINITY_DN234731_c0_g1 | Lake Michigan | -8.8208 | |
| TRINITY_DN214417_c0_g1 | Lake Michigan | 3.8651 | |
| TRINITY_DN246686_c4_g1 | Lake Michigan | 5.9175 | |
| TRINITY_DN202947_c13_g1 | Lake Michigan | 4.0295 | |
| TRINITY_DN169947_c8_g2 | Lake Michigan | -5.3153 | |
| TRINITY_DN229288_c0_g2 | Lake Michigan | 3.6680 | |
| TRINITY_DN144780_c0_g1 | Lake Michigan | 4.4122 | |
| TRINITY_DN178880_c9_g1 | Lake Michigan | 3.5263 | |
| TRINITY_DN241024_c25_g2 | Lake Michigan | 1.8378 | |
| TRINITY_DN244239_c3_g1 | Lake Michigan | 2.1674 | |
| TRINITY_DN217803_c2_g1 | Lake Michigan | 9.6755 | Y7014 |
| TRINITY_DN74931_c0_g1 | Lake Michigan | 2.5567 | |
| TRINITY_DN146704_c0_g1 | Lake Michigan | 4.4866 | |
| TRINITY_DN161794_c1_g1 | Lake Michigan | -6.8341 | |
| TRINITY_DN144575_c1_g1 | Lake Michigan | 6.8172 | |
| TRINITY_DN176111_c7_g1 | Lake Michigan | 2.7580 | |
| TRINITY_DN231826_c4_g1 | Lake Michigan | 3.7454 | |
| TRINITY_DN222391_c5_g4 | Lake Michigan | 1.2884 | |
| TRINITY_DN203447_c9_g1 | Lake Michigan | 2.9490 | |
| TRINITY_DN169546_c7_g1 | Lake Michigan | -2.4318 | |
| TRINITY_DN243148_c1_g1 | Lake Michigan | -3.2739 | |
| TRINITY_DN230376_c14_g1 | Lake Michigan | 1.7004 | |
| TRINITY_DN223020_c5_g1 | Lake Michigan | 3.6328 | |
| TRINITY_DN180979_c12_g1 | Lake Michigan | 2.1050 | CUTA,RL18 |
| TRINITY_DN236653_c9_g1 | Lake Michigan | -2.3610 | |
| TRINITY_DN135213_c0_g1 | Lake Michigan | -7.8051 | |
| TRINITY_DN250843_c4_g1 | Lake Michigan | 3.1622 | |
| TRINITY_DN229479_c4_g3 | Lake Michigan | 4.2292 | |

| | | | |
|---|---|---|---|
| TRINITY_DN196130_c3_g1 | Lake Michigan | -1.7126 | |
| TRINITY_DN100240_c0_g1 | Lake Michigan | 3.4439 | |
| TRINITY_DN168815_c0_g1 | Lake Champlain | -5.3037 | |
| TRINITY_DN205164_c8_g1 | Lake Champlain | -2.8989 | KERA |
| TRINITY_DN248287_c0_g1 | Lake Champlain | -2.4726 | |
| TRINITY_DN245033_c63_g3 | Lake Champlain | -3.0493 | |
| TRINITY_DN186216_c8_g5 | Lake Champlain | -3.0428 | |
| TRINITY_DN178975_c8_g1 | Lake Champlain | -3.3249 | |
| TRINITY_DN248297_c5_g1 | Lake Champlain | -4.0524 | PPM1K |
| TRINITY_DN165427_c8_g1 | Lake Champlain | -3.4241 | MYL3 |
| TRINITY_DN168815_c0_g2 | Lake Champlain | -4.9144 | |
| TRINITY_DN199369_c1_g1 | Lake Champlain | -2.8016 | G3P |
| TRINITY_DN229282_c5_g3 | Lake Champlain | -3.5308 | |
| TRINITY_DN186216_c8_g1 | Lake Champlain | -3.1002 | GLB2,GLB3 |
| TRINITY_DN2553_c0_g1 | Lake Champlain | -2.5918 | |
| TRINITY_DN224894_c3_g2 | Lake Champlain | -2.8114 | MYOC |
| TRINITY_DN240886_c11_g1 | Lake Champlain | -4.1014 | EPYC |
| TRINITY_DN249384_c3_g1 | Lake Champlain | -2.7415 | |
| TRINITY_DN208038_c2_g1 | Lake Champlain | -2.8456 | G3P |
| TRINITY_DN217737_c1_g1 | Lake Champlain | -3.3234 | |
| TRINITY_DN165427_c13_g1 | Lake Champlain | -3.5650 | MLE3 |
| TRINITY_DN185455_c5_g1 | Lake Champlain | -4.0520 | G6PI |
| TRINITY_DN30433_c0_g1 | Lake Champlain | -4.9870 | |
| TRINITY_DN189441_c7_g5 | Lake Champlain | -4.9774 | |
| TRINITY_DN219470_c10_g1 | Lake Champlain | -2.7738 | |
| TRINITY_DN162779_c4_g1 | Lake Champlain | -4.5433 | GLB2,GLB3 |
| TRINITY_DN170662_c5_g1 | Lake Champlain | -2.9863 | FREM2 |
| TRINITY_DN242333_c9_g1 | Lake Champlain | -2.5544 | |
| TRINITY_DN190397_c0_g3 | Lake Champlain | -2.4634 | |
| TRINITY_DN163743_c37_g1 | Lake Champlain | -2.8542 | |
| TRINITY_DN240463_c5_g1 | Lake Champlain | -3.5336 | CO2A1 |
| TRINITY_DN159790_c1_g1 | Lake Champlain | -2.4626 | |
| TRINITY_DN204297_c1_g1 | Lake Champlain | -3.4736 | CO2A1 |
| TRINITY_DN157771_c10_g1 | Lake Champlain | -2.6407 | |
| TRINITY_DN23845_c0_g1 | Lake Champlain | -3.6093 | |
| TRINITY_DN249716_c12_g2 | Lake Champlain | -3.4951 | MYL1,MYL6B |
| TRINITY_DN214738_c4_g2 | Lake Champlain | -3.8219 | |
| TRINITY_DN181359_c14_g1 | Lake Champlain | -3.1773 | |
| TRINITY_DN202230_c6_g1 | Lake Champlain | -3.3788 | MLE1 |
| TRINITY_DN231480_c4_g1 | Lake Champlain | -3.2592 | CO2A1 |
| TRINITY_DN239456_c1_g2 | Lake Champlain | -2.5667 | |
| TRINITY_DN229890_c14_g3 | Lake Champlain | 8.3861 | |
| TRINITY_DN157771_c9_g1 | Lake Champlain | -2.9817 | |

| | | | |
|---|---|---|---|
| TRINITY_DN159823_c0_g2 | Lake Champlain | -2.7849 | |
| TRINITY_DN196668_c6_g2 | Lake Champlain | -4.8718 | |
| TRINITY_DN199279_c3_g1 | Lake Champlain | -2.7237 | GLB,GLB1,GLB2,GLB5 |
| TRINITY_DN247106_c3_g1 | Lake Champlain | -3.0897 | CO2A1 |
| TRINITY_DN245937_c6_g1 | Lake Champlain | -3.8217 | |
| TRINITY_DN200281_c1_g1 | Lake Champlain | -2.8871 | |
| TRINITY_DN14422_c40_g1 | Lake Champlain | -3.0625 | |
| TRINITY_DN199043_c2_g3 | Lake Champlain | -2.4206 | |
| TRINITY_DN218013_c6_g1 | Lake Champlain | 2.3888 | MPRIP,TARA |
| TRINITY_DN160162_c6_g1 | Lake Champlain | -2.8310 | |
| TRINITY_DN185435_c0_g1 | Lake Champlain | -3.6885 | |
| TRINITY_DN249845_c13_g6 | Lake Champlain | -2.3588 | |
| TRINITY_DN190029_c8_g1 | Lake Champlain | -3.4134 | ACT,ACTS |
| TRINITY_DN107086_c0_g1 | Lake Champlain | -2.7438 | |
| TRINITY_DN190397_c0_g2 | Lake Champlain | -2.8727 | |
| TRINITY_DN247106_c3_g5 | Lake Champlain | -2.0573 | CO2A1 |
| TRINITY_DN96454_c0_g1 | Lake Champlain | -2.9564 | |
| TRINITY_DN156151_c1_g1 | Lake Champlain | -2.3033 | |
| TRINITY_DN209960_c7_g1 | Lake Champlain | -3.4411 | |
| TRINITY_DN199279_c3_g2 | Lake Champlain | -3.2693 | |
| TRINITY_DN208037_c5_g17 | Lake Champlain | -3.7256 | |
| TRINITY_DN110996_c0_g1 | Lake Champlain | -2.5457 | |
| TRINITY_DN162890_c7_g1 | Lake Champlain | -3.0285 | |
| TRINITY_DN245033_c66_g1 | Lake Champlain | -2.6344 | |
| TRINITY_DN247506_c1_g1 | Lake Champlain | -2.9362 | PHF3,DIDO1 |
| TRINITY_DN158511_c0_g1 | Lake Champlain | -2.5703 | |
| TRINITY_DN177977_c18_g1 | Lake Champlain | -2.4642 | ACTS |
| TRINITY_DN157443_c1_g1 | Lake Champlain | -2.7712 | |
| TRINITY_DN202612_c11_g2 | Lake Champlain | -3.8029 | |
| TRINITY_DN173616_c5_g1 | Lake Champlain | -2.8685 | G3P,G3PT |
| TRINITY_DN133107_c0_g1 | Lake Champlain | -4.4417 | |
| TRINITY_DN178414_c9_g3 | Lake Champlain | -2.3144 | |
| TRINITY_DN240517_c6_g1 | Lake Champlain | -2.6181 | |
| TRINITY_DN248287_c0_g2 | Lake Champlain | -2.2739 | |
| TRINITY_DN236579_c10_g1 | Lake Champlain | -2.6602 | |
| TRINITY_DN245033_c22_g1 | Lake Champlain | -2.6439 | G3P |
| TRINITY_DN194668_c12_g1 | Lake Champlain | -3.8362 | |
| TRINITY_DN202958_c6_g1 | Lake Champlain | -2.7215 | |
| TRINITY_DN217575_c0_g1 | Lake Champlain | -3.8870 | |
| TRINITY_DN240913_c0_g2 | Lake Champlain | -2.3595 | GLB5 |
| TRINITY_DN233988_c6_g4 | Lake Champlain | -2.2550 | |
| TRINITY_DN182265_c19_g1 | Lake Champlain | -2.3175 | |
| TRINITY_DN247773_c1_g1 | Lake Champlain | -2.8495 | GLB1 |

| | | | |
|---|---|---|---|
| TRINITY_DN207341_c1_g1 | Lake Champlain | -4.0924 | |
| TRINITY_DN11690_c0_g1 | Lake Champlain | -2.3176 | |
| TRINITY_DN182265_c12_g6 | Lake Champlain | -2.1716 | |
| TRINITY_DN177977_c14_g5 | Lake Champlain | -2.6000 | ACTS |
| TRINITY_DN267216_c0_g1 | Lake Champlain | -2.5375 | |
| TRINITY_DN217945_c2_g3 | Lake Champlain | -2.0682 | |
| TRINITY_DN229836_c5_g2 | Lake Champlain | -2.0410 | CLC3A,TETN |
| TRINITY_DN242348_c2_g1 | Lake Champlain | -2.1422 | COBA1,COBA2 |
| TRINITY_DN74931_c0_g1 | Lake Champlain | -2.7217 | |
| TRINITY_DN233988_c6_g2 | Lake Champlain | -1.7659 | F16P2 |
| TRINITY_DN182048_c4_g4 | Lake Champlain | -2.5385 | |
| TRINITY_DN191473_c3_g3 | Lake Champlain | -2.5574 | |
| TRINITY_DN250749_c0_g1 | Lake Champlain | -2.8854 | G3P |
| TRINITY_DN165356_c2_g1 | Lake Champlain | -1.7433 | |
| TRINITY_DN249416_c1_g3 | Lake Champlain | 4.7845 | |
| TRINITY_DN188246_c6_g1 | Lake Champlain | -4.3882 | |
| TRINITY_DN184382_c0_g1 | Lake Champlain | -3.4839 | |
| TRINITY_DN205658_c0_g1 | Lake Champlain | -2.6860 | CAD13 |
| TRINITY_DN217499_c4_g1 | Lake Champlain | -2.7982 | ADA10 |
| TRINITY_DN247106_c3_g7 | Lake Champlain | -1.8809 | CO2A1 |
| TRINITY_DN236314_c3_g1 | Lake Champlain | -2.3953 | MYH7 |
| TRINITY_DN197078_c11_g11 | Lake Champlain | -3.3714 | MLRV |
| TRINITY_DN182265_c7_g1 | Lake Champlain | -2.1730 | |
| TRINITY_DN224108_c10_g1 | Lake Champlain | -2.2189 | |
| TRINITY_DN182265_c12_g8 | Lake Champlain | -2.0917 | |
| TRINITY_DN224144_c8_g1 | Lake Champlain | -2.3641 | |
| TRINITY_DN186216_c8_g3 | Lake Champlain | -2.4631 | GLB1,GLB3 |
| TRINITY_DN244658_c31_g18 | Lake Champlain | -2.1206 | MYH1B |
| TRINITY_DN182265_c12_g16 | Lake Champlain | -1.9728 | |
| TRINITY_DN247352_c10_g1 | Lake Champlain | -2.2373 | TPM1 |
| TRINITY_DN247352_c4_g1 | Lake Champlain | -2.2622 | TPM1,TPM3 |
| TRINITY_DN224901_c0_g3 | Lake Champlain | -2.4371 | TBFA |
| TRINITY_DN212694_c7_g5 | Lake Champlain | -2.5480 | MYPC1 |
| TRINITY_DN209792_c2_g1 | Lake Champlain | 4.8222 | UBIQP |
| TRINITY_DN208038_c0_g2 | Lake Champlain | -2.5621 | G3P |
| TRINITY_DN233489_c1_g1 | Lake Champlain | -1.7205 | F16P1,F16P2 |
| TRINITY_DN244517_c11_g5 | Lake Champlain | 4.5327 | TDH |
| TRINITY_DN197078_c10_g4 | Lake Champlain | -3.2027 | MLRB,MLRS,MLRV,MYL10 |
| TRINITY_DN233837_c4_g2 | Lake Champlain | -2.4286 | |
| TRINITY_DN244996_c3_g1 | Lake Champlain | -2.7433 | G3P |
| TRINITY_DN203123_c4_g2 | Lake Champlain | -2.5713 | ACT |
| TRINITY_DN236699_c9_g1 | Lake Champlain | -4.4046 | |
| TRINITY_DN203783_c2_g1 | Lake Champlain | -2.0542 | GLB1,GLB2 |

| | | | |
|---|---|---|---|
| TRINITY_DN214738_c4_g1 | Lake Champlain | -3.7446 | |
| TRINITY_DN217945_c2_g1 | Lake Champlain | -1.9069 | |
| TRINITY_DN238868_c1_g2 | Lake Champlain | -2.3411 | MYSS,MYH7,MYH8,MYH13 |
| TRINITY_DN225692_c1_g3 | Lake Champlain | -4.2534 | |
| TRINITY_DN241984_c24_g1 | Lake Champlain | -2.2732 | MYSS |
| TRINITY_DN224874_c7_g1 | Lake Champlain | 1.8945 | |
| TRINITY_DN243449_c8_g3 | Lake Champlain | -2.5120 | CO9A1,CO9A3 |
| TRINITY_DN167456_c7_g1 | Lake Champlain | -2.1295 | EMC1 |
| TRINITY_DN182265_c4_g1 | Lake Champlain | -2.0470 | |
| TRINITY_DN202762_c0_g1 | Lake Champlain | -1.8669 | KPYM |
| TRINITY_DN163738_c0_g1 | Lake Champlain | 7.6053 | RL29 |
| TRINITY_DN245355_c9_g1 | Lake Champlain | -8.4114 | |
| TRINITY_DN228356_c13_g5 | Lake Champlain | -1.6989 | TPIS |
| TRINITY_DN213866_c1_g1 | Lake Champlain | -2.0589 | PGS2 |
| TRINITY_DN165427_c8_g2 | Lake Champlain | -3.0097 | MYL1,MYL6,MYL6B,MLEX |
| TRINITY_DN237665_c5_g1 | Lake Champlain | -2.5515 | ACTS,ACTM |
| TRINITY_DN224515_c6_g1 | Lake Champlain | -2.0755 | TPM1 |
| TRINITY_DN244475_c11_g1 | Lake Champlain | -2.1747 | TPM1 |
| TRINITY_DN249841_c4_g1 | Lake Champlain | -2.9674 | |
| TRINITY_DN244658_c36_g2 | Lake Champlain | -2.2485 | MYH4,MYH7,MYSS |
| TRINITY_DN159045_c5_g1 | Lake Champlain | -8.7175 | |
| TRINITY_DN242410_c13_g3 | Lake Champlain | -2.3077 | |
| TRINITY_DN184222_c3_g2 | Lake Champlain | -3.2623 | |
| TRINITY_DN242399_c12_g1 | Lake Champlain | -1.7129 | LDH |
| TRINITY_DN185234_c2_g1 | Lake Champlain | 2.1332 | KCAB1,KCAB2 |
| TRINITY_DN243771_c2_g1 | Lake Champlain | -3.1122 | MLRV |
| TRINITY_DN244996_c2_g1 | Lake Champlain | -2.8961 | G3P |
| TRINITY_DN206156_c4_g2 | Lake Champlain | 2.1648 | |
| TRINITY_DN187287_c7_g1 | Lake Champlain | -2.0012 | TNNC1 |
| TRINITY_DN175891_c9_g1 | Lake Champlain | 1.7803 | TFE3 |
| TRINITY_DN244517_c10_g1 | Lake Champlain | 5.3285 | |
| TRINITY_DN177977_c15_g4 | Lake Champlain | -2.4157 | ACTS |
| TRINITY_DN240228_c2_g1 | Lake Champlain | -3.0748 | |
| TRINITY_DN250186_c4_g1 | Lake Champlain | -2.6104 | |
| TRINITY_DN177977_c28_g1 | Lake Champlain | -2.4720 | ACTS |
| TRINITY_DN249845_c13_g3 | Lake Champlain | -1.5312 | |
| TRINITY_DN209664_c8_g2 | Lake Champlain | -2.8933 | ACYP2 |
| TRINITY_DN169755_c13_g1 | Lake Champlain | -2.8980 | |
| TRINITY_DN171668_c1_g1 | Lake Champlain | -2.4736 | |
| TRINITY_DN163160_c9_g7 | Lake Champlain | -1.4084 | |
| TRINITY_DN163706_c5_g1 | Lake Champlain | -2.1338 | TPM3 |
| TRINITY_DN165427_c12_g1 | Lake Champlain | -2.9297 | MLE3,MYL1 |
| TRINITY_DN196430_c7_g3 | Lake Champlain | 1.9756 | MPRIP |

| | | | |
|---|---|---|---|
| TRINITY_DN231480_c7_g1 | Lake Champlain | -2.1007 | CO2A1 |
| TRINITY_DN224901_c0_g1 | Lake Champlain | -2.3240 | TBFA |
| TRINITY_DN234334_c3_g3 | Lake Champlain | -1.7698 | COBA2,CO5A1,CO9A2 |
| TRINITY_DN159119_c0_g1 | Lake Champlain | 1.7549 | |
| TRINITY_DN177977_c14_g3 | Lake Champlain | -2.4559 | |
| TRINITY_DN214612_c5_g2 | Lake Champlain | -1.4443 | |
| TRINITY_DN246813_c5_g2 | Lake Champlain | -2.3335 | TBFA |
| TRINITY_DN201613_c7_g1 | Lake Champlain | -2.0879 | |
| TRINITY_DN165427_c10_g1 | Lake Champlain | -2.9399 | MYL6 |
| TRINITY_DN247106_c3_g3 | Lake Champlain | -1.9523 | CO2A1 |
| TRINITY_DN247940_c11_g1 | Lake Champlain | 1.5725 | PDLI3 |
| TRINITY_DN158558_c1_g1 | Lake Champlain | -2.4125 | |
| TRINITY_DN198727_c4_g1 | Lake Champlain | -2.6901 | CO9A1,CO1A2,CO9A3,COLL2,COLL7 |
| TRINITY_DN198430_c0_g2 | Lake Champlain | -2.8039 | ACT2,ACTH,ACTC |
| TRINITY_DN210879_c0_g1 | Lake Champlain | -2.3338 | MYPC,MYPC1,MYPC2,MYPC3,TDRD5 |
| TRINITY_DN176525_c5_g1 | Lake Champlain | -2.7265 | G3P |
| TRINITY_DN208871_c3_g1 | Lake Champlain | -2.8298 | KAD1 |
| TRINITY_DN183699_c8_g1 | Lake Champlain | 3.1470 | |
| TRINITY_DN167889_c0_g1 | Lake Champlain | -1.4899 | |
| TRINITY_DN249254_c9_g1 | Lake Champlain | -1.6249 | |
| TRINITY_DN54029_c3_g1 | Lake Champlain | -2.3596 | |
| TRINITY_DN215549_c37_g1 | Lake Champlain | -3.0908 | ACT1 |
| TRINITY_DN224901_c2_g1 | Lake Champlain | -2.4010 | TBFA |
| TRINITY_DN128471_c0_g1 | Lake Champlain | -2.6341 | |
| TRINITY_DN246307_c2_g1 | Lake Champlain | -1.9686 | CO2A1,CO1A1,CO1A2 |
| TRINITY_DN244658_c2_g1 | Lake Champlain | -2.0027 | MYSS |
| TRINITY_DN243076_c3_g2 | Lake Champlain | -1.5125 | |
| TRINITY_DN217653_c0_g1 | Lake Champlain | -2.1142 | |
| TRINITY_DN180883_c2_g1 | Lake Champlain | -2.0556 | TPM3 |
| TRINITY_DN178015_c1_g1 | Lake Champlain | -1.5055 | LDH |
| TRINITY_DN247773_c3_g1 | Lake Champlain | -2.6763 | GLB1,GLB2,GLB3 |
| TRINITY_DN215105_c5_g1 | Lake Champlain | -1.6796 | HACD1,HACD2 |
| TRINITY_DN169873_c5_g4 | Lake Champlain | -1.9709 | APT |
| TRINITY_DN198884_c2_g1 | Lake Champlain | 4.1846 | |
| TRINITY_DN206846_c2_g1 | Lake Champlain | -2.3316 | DUPD1 |
| TRINITY_DN176738_c6_g1 | Lake Champlain | -1.6418 | |
| TRINITY_DN249841_c2_g2 | Lake Champlain | -2.5588 | |
| TRINITY_DN236767_c6_g1 | Lake Champlain | -3.4864 | |
| TRINITY_DN183227_c3_g1 | Lake Champlain | 1.4914 | CYSP2,CATL,CATL1 |
| TRINITY_DN236344_c5_g1 | Lake Champlain | 1.5282 | |
| TRINITY_DN244658_c36_g6 | Lake Champlain | -2.0448 | MYH4 |
| TRINITY_DN215549_c21_g3 | Lake Champlain | -2.3794 | ACTA |
| TRINITY_DN243410_c10_g1 | Lake Champlain | -2.6175 | ACT2,ACT3,ACTA,ACTC,ACTH |

| | | | |
|---|---|---|---|
| TRINITY_DN206539_c4_g1 | Lake Champlain | -1.4115 | |
| TRINITY_DN244517_c11_g3 | Lake Champlain | 5.6460 | |
| TRINITY_DN245496_c2_g1 | Lake Champlain | -1.7744 | MYOZ2,MYOZ3 |
| TRINITY_DN215549_c21_g2 | Lake Champlain | -2.8319 | ACTA,ACTC |
| TRINITY_DN222395_c3_g1 | Lake Champlain | -1.6513 | JPH1,JPH2,JPH3 |
| TRINITY_DN208614_c1_g1 | Lake Champlain | -1.7279 | |
| TRINITY_DN224515_c5_g1 | Lake Champlain | -2.1051 | TPM1 |
| TRINITY_DN215680_c36_g1 | Lake Champlain | -2.2778 | CNMD |
| TRINITY_DN217517_c3_g1 | Lake Champlain | -2.9994 | DCR1A |
| TRINITY_DN243771_c0_g1 | Lake Champlain | -3.1790 | LAMBV,MLRA,MLRB,MLRV |
| TRINITY_DN244517_c4_g1 | Lake Champlain | 4.1436 | |
| TRINITY_DN241984_c17_g4 | Lake Champlain | -2.3142 | MYH6 |
| TRINITY_DN214388_c5_g1 | Lake Champlain | 2.1923 | |
| TRINITY_DN249845_c10_g2 | Lake Champlain | -2.2941 | |
| TRINITY_DN229826_c2_g1 | Lake Champlain | -2.2291 | |
| TRINITY_DN202230_c4_g1 | Lake Champlain | -2.9293 | MYL6 |
| TRINITY_DN248030_c23_g1 | Lake Champlain | -6.0866 | |
| TRINITY_DN177066_c11_g1 | Lake Champlain | -2.0902 | MYSS |
| TRINITY_DN245033_c63_g1 | Lake Champlain | -2.6562 | G3P |
| TRINITY_DN182265_c13_g2 | Lake Champlain | -2.1055 | TNNC1 |
| TRINITY_DN232060_c5_g6 | Lake Champlain | -3.3703 | |
| TRINITY_DN181670_c5_g3 | Lake Champlain | -1.7086 | POPD1 |
| TRINITY_DN187042_c4_g1 | Lake Champlain | 5.2339 | |
| TRINITY_DN213893_c10_g4 | Lake Champlain | -1.9191 | TNNI1 |
| TRINITY_DN242399_c11_g5 | Lake Champlain | -1.5138 | LDH |
| TRINITY_DN197078_c12_g1 | Lake Champlain | -2.9957 | MLRV |
| TRINITY_DN168593_c13_g2 | Lake Champlain | -3.4186 | |
| TRINITY_DN231778_c0_g1 | Lake Champlain | -2.6936 | CO1A2,CO2A1,CO5A2 |
| TRINITY_DN250706_c2_g1 | Lake Champlain | -2.8521 | |
| TRINITY_DN247789_c4_g1 | Lake Champlain | -1.7717 | |
| TRINITY_DN52811_c0_g1 | Lake Champlain | -2.1002 | |
| TRINITY_DN197078_c10_g5 | Lake Champlain | -3.1225 | MYL10,MLRV |
| TRINITY_DN162316_c5_g2 | Lake Champlain | -2.2551 | |
| TRINITY_DN187952_c11_g1 | Lake Champlain | -2.3034 | |
| TRINITY_DN207068_c16_g1 | Lake Champlain | -1.6702 | TNNT2 |
| TRINITY_DN249254_c9_g3 | Lake Champlain | -1.8000 | |
| TRINITY_DN246556_c5_g1 | Lake Champlain | -1.3564 | CO6A3,CO6A4 |
| TRINITY_DN218064_c5_g1 | Lake Champlain | -1.2227 | |
| TRINITY_DN244517_c11_g7 | Lake Champlain | 4.8621 | |
| TRINITY_DN229826_c14_g2 | Lake Champlain | -1.9751 | |
| TRINITY_DN237119_c2_g1 | Lake Champlain | -2.0291 | CO1A2,CO3A1,CO9A1,CO9A3,COLA1,COLL2 |
| TRINITY_DN165792_c7_g1 | Lake Champlain | -1.2408 | IGF1,IGF2 |
| TRINITY_DN192672_c6_g1 | Lake Champlain | -2.0533 | MYSS |

| | | | |
|---|---|---|---|
| TRINITY_DN244658_c35_g6 | Lake Champlain | -2.3282 | MYH4,MYH6 |
| TRINITY_DN240997_c0_g1 | Lake Champlain | -4.3936 | |
| TRINITY_DN227020_c4_g1 | Lake Champlain | 1.6683 | |
| TRINITY_DN216236_c4_g1 | Lake Champlain | -1.5654 | |
| TRINITY_DN182265_c12_g15 | Lake Champlain | -1.9910 | TNNC1 |
| TRINITY_DN241691_c1_g1 | Lake Champlain | -1.3657 | FNDC1 |
| TRINITY_DN243771_c1_g1 | Lake Champlain | -2.9555 | MLRV |
| TRINITY_DN238310_c3_g1 | Lake Champlain | -1.5221 | TNNC2 |
| TRINITY_DN177977_c14_g1 | Lake Champlain | -2.5227 | |
| TRINITY_DN225863_c6_g1 | Lake Champlain | -1.9629 | GLB,GLB2,GLB3 |
| TRINITY_DN185741_c2_g1 | Lake Champlain | -2.1083 | KPYM |
| TRINITY_DN249254_c9_g2 | Lake Champlain | -1.5857 | PYGM |
| TRINITY_DN247854_c4_g4 | Lake Champlain | -1.2691 | LAMA5,NET1 |
| TRINITY_DN247885_c1_g1 | Lake Champlain | 2.4094 | |
| TRINITY_DN369478_c16_g1 | Lake Champlain | -2.3632 | |
| TRINITY_DN244658_c62_g1 | Lake Champlain | -2.0254 | MYH7 |
| TRINITY_DN243281_c1_g1 | Lake Champlain | 1.7196 | GCSP |
| TRINITY_DN204297_c3_g3 | Lake Champlain | -2.6683 | CO1A1,CO1A2 |
| TRINITY_DN215549_c21_g1 | Lake Champlain | -2.5134 | ACT,ACT2,ACT3,ACTA,ACTC,ACTM,ACTS |
| TRINITY_DN247940_c11_g3 | Lake Champlain | 1.3658 | |
| TRINITY_DN225341_c1_g2 | Lake Champlain | -2.4481 | |
| TRINITY_DN246813_c5_g1 | Lake Champlain | -2.2788 | |
| TRINITY_DN244658_c48_g1 | Lake Champlain | -2.0570 | MYH1B |
| TRINITY_DN182265_c12_g12 | Lake Champlain | -2.0104 | TNNC1 |
| TRINITY_DN245388_c7_g2 | Lake Champlain | -3.4911 | |
| TRINITY_DN224515_c10_g1 | Lake Champlain | -2.0832 | TPM1 |
| TRINITY_DN230055_c1_g1 | Lake Champlain | -2.8046 | |
| TRINITY_DN229051_c13_g2 | Lake Champlain | 1.3604 | |
| TRINITY_DN240304_c2_g2 | Lake Champlain | 1.7563 | KCAB1,KCAB2 |
| TRINITY_DN199043_c2_g1 | Lake Champlain | -2.0989 | |
| TRINITY_DN185741_c1_g1 | Lake Champlain | -1.8977 | KPYM |
| TRINITY_DN237789_c3_g3 | Lake Champlain | -2.4005 | G6PI |
| TRINITY_DN181616_c6_g1 | Lake Champlain | -3.8160 | |
| TRINITY_DN196771_c9_g2 | Lake Champlain | -2.3685 | |
| TRINITY_DN232012_c4_g1 | Lake Champlain | -2.5591 | |
| TRINITY_DN232012_c1_g1 | Lake Champlain | -3.6935 | |
| TRINITY_DN197078_c11_g10 | Lake Champlain | -2.9290 | MLRB |
| TRINITY_DN247681_c5_g2 | Lake Champlain | 1.8100 | |
| TRINITY_DN177977_c14_g2 | Lake Champlain | -2.1385 | ACTS |
| TRINITY_DN205499_c6_g2 | Lake Champlain | 1.5813 | GCSP |
| TRINITY_DN233908_c7_g1 | Lake Champlain | -1.0309 | |
| TRINITY_DN218793_c3_g1 | Lake Champlain | 1.7260 | ADAL |
| TRINITY_DN200582_c3_g1 | Lake Champlain | 2.3212 | ASB16,ASB18 |

| | | | |
|---|---|---|---|
| TRINITY_DN198100_c17_g3 | Lake Champlain | -1.7290 | KPYM |
| TRINITY_DN177977_c15_g13 | Lake Champlain | -2.6199 | ACTS |
| TRINITY_DN248676_c0_g1 | Lake Champlain | -2.5624 | GLB1,GLB3 |
| TRINITY_DN211274_c2_g2 | Lake Champlain | -2.6503 | CO2A1,CO3A1 |
| TRINITY_DN244996_c1_g1 | Lake Champlain | -2.6065 | G3P |
| TRINITY_DN172572_c0_g1 | Lake Champlain | -2.4384 | |
| TRINITY_DN235405_c0_g1 | Lake Champlain | -2.4810 | |
| TRINITY_DN186216_c9_g3 | Lake Champlain | -1.8746 | GLB,GLB1 |
| TRINITY_DN243609_c6_g1 | Lake Champlain | -2.2677 | |
| TRINITY_DN211272_c6_g1 | Lake Champlain | -2.4072 | ACTC,ACTS,ACT2 |
| TRINITY_DN249845_c13_g1 | Lake Champlain | -1.4795 | |
| TRINITY_DN172817_c14_g1 | Lake Champlain | 1.9105 | TRI54 |
| TRINITY_DN249845_c13_g5 | Lake Champlain | -1.5774 | |
| TRINITY_DN223766_c2_g1 | Lake Champlain | -1.9878 | CQ067 |
| TRINITY_DN235351_c1_g2 | Lake Champlain | -2.5120 | ACTA |
| TRINITY_DN167893_c0_g1 | Lake Champlain | -2.3589 | |
| TRINITY_DN170085_c10_g1 | Lake Champlain | -2.0706 | OTOR |
| TRINITY_DN225023_c6_g1 | Lake Champlain | 1.5444 | DHTK1 |
| TRINITY_DN199619_c1_g1 | Lake Champlain | -1.5949 | |
| TRINITY_DN213368_c8_g1 | Lake Champlain | 1.2489 | |
| TRINITY_DN239659_c1_g1 | Lake Champlain | -1.9481 | MYSS |
| TRINITY_DN241984_c17_g1 | Lake Champlain | -2.2391 | MYSS |
| TRINITY_DN249845_c13_g2 | Lake Champlain | -1.5211 | |
| TRINITY_DN242336_c14_g2 | Lake Champlain | -1.5274 | MOT2 |
| TRINITY_DN249254_c14_g1 | Lake Champlain | -1.7157 | |
| TRINITY_DN241984_c6_g1 | Lake Champlain | -2.2189 | MYSS |
| TRINITY_DN239161_c8_g1 | Lake Champlain | -1.4550 | |
| TRINITY_DN182265_c21_g1 | Lake Champlain | -2.0709 | TNNC1 |
| TRINITY_DN207068_c6_g3 | Lake Champlain | -1.3069 | |
| TRINITY_DN203167_c3_g2 | Lake Champlain | -1.7688 | GLB |
| TRINITY_DN243453_c2_g1 | Lake Champlain | -1.5470 | |
| TRINITY_DN203205_c2_g1 | Lake Champlain | -3.3925 | |
| TRINITY_DN206403_c9_g1 | Lake Champlain | -2.1853 | DCR1A |
| TRINITY_DN236434_c6_g1 | Lake Champlain | 8.2977 | |
| TRINITY_DN159823_c0_g1 | Lake Champlain | -2.3710 | |
| TRINITY_DN184249_c8_g1 | Lake Champlain | -1.9349 | |
| TRINITY_DN231195_c2_g1 | Lake Champlain | -1.9422 | |
| TRINITY_DN248817_c0_g4 | Lake Champlain | -1.2185 | |
| TRINITY_DN246020_c3_g1 | Lake Champlain | -2.9299 | |
| TRINITY_DN222634_c1_g1 | Lake Champlain | -2.2143 | MYH6 |
| TRINITY_DN228356_c13_g1 | Lake Champlain | -1.6022 | TPIS |
| TRINITY_DN242211_c17_g2 | Lake Champlain | 1.2091 | |
| TRINITY_DN244658_c36_g1 | Lake Champlain | -2.1472 | MYSS |

| | | | |
|---|---|---|---|
| TRINITY_DN182265_c12_g10 | Lake Champlain | -1.9973 | TNNC1 |
| TRINITY_DN190397_c4_g1 | Lake Champlain | -2.4730 | |
| TRINITY_DN197078_c15_g1 | Lake Champlain | -3.1083 | MLRV |
| TRINITY_DN162378_c8_g1 | Lake Champlain | -2.1532 | |
| TRINITY_DN232012_c0_g1 | Lake Champlain | -2.3126 | |
| TRINITY_DN244475_c20_g1 | Lake Champlain | -1.9125 | TPM3 |
| TRINITY_DN163919_c11_g1 | Lake Champlain | -2.2316 | ACTA |
| TRINITY_DN246813_c5_g4 | Lake Champlain | -2.0467 | TBFA |
| TRINITY_DN247789_c8_g1 | Lake Champlain | -1.8656 | |
| TRINITY_DN192672_c2_g1 | Lake Champlain | -2.0049 | MYSS,MYH7 |
| TRINITY_DN238310_c2_g1 | Lake Champlain | -1.4380 | |
| TRINITY_DN196452_c3_g2 | Lake Champlain | -2.3877 | MYH7 |
| TRINITY_DN182265_c12_g14 | Lake Champlain | -1.9926 | TNNC1 |
| TRINITY_DN185920_c7_g1 | Lake Champlain | 2.0004 | |
| TRINITY_DN201262_c0_g4 | Lake Champlain | -3.2587 | |
| TRINITY_DN208255_c3_g1 | Lake Champlain | -1.8205 | |
| TRINITY_DN249845_c8_g2 | Lake Champlain | -1.4305 | |
| TRINITY_DN212438_c5_g1 | Lake Champlain | -1.7142 | |
| TRINITY_DN249845_c11_g4 | Lake Champlain | -1.5254 | |
| TRINITY_DN210975_c0_g1 | Lake Champlain | -1.2690 | CO6A2 |
| TRINITY_DN229651_c4_g2 | Lake Champlain | -1.6091 | |
| TRINITY_DN244658_c31_g16 | Lake Champlain | -1.9186 | MYH1B |
| TRINITY_DN244475_c12_g4 | Lake Champlain | -1.8632 | TPM1 |
| TRINITY_DN233837_c4_g1 | Lake Champlain | -2.1914 | |
| TRINITY_DN242474_c6_g1 | Lake Champlain | -2.9707 | MLRV |
| TRINITY_DN199622_c3_g2 | Lake Champlain | 1.7170 | |
| TRINITY_DN170672_c11_g1 | Lake Champlain | -1.3718 | |
| TRINITY_DN194080_c11_g1 | Lake Champlain | -1.4759 | |
| TRINITY_DN247681_c5_g3 | Lake Champlain | 1.6520 | |
| TRINITY_DN225454_c1_g2 | Lake Champlain | -1.4295 | FBLN4 |
| TRINITY_DN124792_c0_g1 | Lake Champlain | -1.6681 | |
| TRINITY_DN240150_c2_g1 | Lake Champlain | 1.6490 | TRI54 |
| TRINITY_DN196654_c2_g1 | Lake Champlain | -1.5734 | DR9C7,CSGA |
| TRINITY_DN194091_c1_g1 | Lake Champlain | -1.8328 | MYH7,MYH8 |
| TRINITY_DN235400_c2_g3 | Lake Champlain | -1.4161 | |
| TRINITY_DN173705_c12_g4 | Lake Champlain | 1.9819 | CGL |
| TRINITY_DN188609_c13_g1 | Lake Champlain | 2.0613 | |
| TRINITY_DN188579_c5_g1 | Lake Champlain | -2.5824 | |
| TRINITY_DN198100_c16_g5 | Lake Champlain | -2.0432 | KPYM |
| TRINITY_DN189548_c0_g1 | Lake Champlain | -2.7454 | CO9A3 |
| TRINITY_DN208945_c0_g1 | Lake Champlain | -1.7640 | |
| TRINITY_DN244310_c4_g3 | Lake Champlain | -1.7497 | MYOM1,MYOM2 |
| TRINITY_DN240567_c8_g4 | Lake Champlain | -1.6185 | |

| | | | |
|---|---|---|---|
| TRINITY_DN178091_c9_g3 | Lake Champlain | -2.9462 | |
| TRINITY_DN249860_c8_g1 | Lake Champlain | -1.2423 | MDHC |
| TRINITY_DN223153_c3_g1 | Lake Champlain | -2.7152 | |
| TRINITY_DN244049_c4_g2 | Lake Champlain | -2.0694 | |
| TRINITY_DN187952_c4_g1 | Lake Champlain | -2.3324 | |
| TRINITY_DN185741_c1_g2 | Lake Champlain | -1.6243 | KPYM |
| TRINITY_DN12436_c0_g1 | Lake Champlain | 2.1273 | EIF3I |
| TRINITY_DN229263_c5_g1 | Lake Champlain | -1.3993 | BGH3,POSTN |
| TRINITY_DN182265_c12_g7 | Lake Champlain | -1.9572 | |
| TRINITY_DN177977_c14_g4 | Lake Champlain | -2.3624 | |
| TRINITY_DN218888_c5_g1 | Lake Champlain | 1.2239 | GBRL2 |
| TRINITY_DN174447_c0_g1 | Lake Champlain | -1.3232 | LUM |
| TRINITY_DN195548_c6_g1 | Lake Champlain | -1.4388 | DERM |
| TRINITY_DN198110_c3_g2 | Lake Champlain | -2.6293 | GLB2 |
| TRINITY_DN233416_c3_g2 | Lake Champlain | 1.2709 | |
| TRINITY_DN241381_c3_g1 | Lake Champlain | -1.3200 | COCA1,COEA1 |
| TRINITY_DN243609_c5_g1 | Lake Champlain | -2.4566 | |
| TRINITY_DN229783_c19_g1 | Lake Champlain | 2.1181 | |
| TRINITY_DN199811_c0_g1 | Lake Champlain | -2.0013 | ACTS,ACTM,ACT3,ACT |
| TRINITY_DN244658_c31_g4 | Lake Champlain | -2.5911 | MYSS |
| TRINITY_DN205711_c10_g1 | Lake Champlain | -1.4960 | TNR14 |
| TRINITY_DN177977_c15_g2 | Lake Champlain | -2.7866 | |
| TRINITY_DN180020_c7_g1 | Lake Champlain | -2.3645 | |
| TRINITY_DN177977_c15_g5 | Lake Champlain | -2.3925 | |
| TRINITY_DN242526_c5_g1 | Lake Champlain | -1.5677 | |
| TRINITY_DN212694_c7_g1 | Lake Champlain | -2.2840 | MYPC1,MYPC2 |
| TRINITY_DN239659_c0_g1 | Lake Champlain | -2.0821 | MYH4,MYH6,MYSS,MYH1B |
| TRINITY_DN185094_c8_g2 | Lake Champlain | -1.4543 | |
| TRINITY_DN158963_c0_g1 | Lake Champlain | 1.5111 | UBIQP |
| TRINITY_DN192666_c10_g1 | Lake Champlain | 2.0038 | |
| TRINITY_DN238231_c2_g7 | Lake Champlain | -1.0512 | |
| TRINITY_DN171254_c16_g1 | Lake Champlain | 1.8837 | |
| TRINITY_DN204470_c5_g1 | Lake Champlain | -3.8385 | |
| TRINITY_DN243158_c8_g2 | Lake Champlain | -1.6915 | C1QT5 |
| TRINITY_DN243131_c1_g5 | Lake Champlain | 1.3091 | ACACB,ACAC |
| TRINITY_DN242403_c0_g1 | Lake Champlain | -1.3313 | |
| TRINITY_DN176998_c7_g2 | Lake Champlain | -1.1978 | |
| TRINITY_DN233986_c12_g1 | Lake Champlain | -1.3726 | PGS1 |
| TRINITY_DN160136_c9_g1 | Lake Champlain | 6.7645 | |
| TRINITY_DN244658_c41_g1 | Lake Champlain | -2.0323 | MYH7 |
| TRINITY_DN196348_c6_g5 | Lake Champlain | -1.3315 | |
| TRINITY_DN213893_c16_g1 | Lake Champlain | -1.6557 | TNNI1 |
| TRINITY_DN197078_c11_g3 | Lake Champlain | -2.9533 | MLRS,MLRV |

| | | | |
|---|---|---|---|
| TRINITY_DN247940_c7_g1 | Lake Champlain | 1.3004 | |
| TRINITY_DN244658_c35_g8 | Lake Champlain | -1.9291 | MYH6 |
| TRINITY_DN244658_c32_g14 | Lake Champlain | -1.7183 | MYH8,MYSS |
| TRINITY_DN229890_c12_g2 | Lake Champlain | 4.5188 | |
| TRINITY_DN234114_c8_g6 | Lake Champlain | -1.3209 | KAD1 |
| TRINITY_DN245033_c6_g1 | Lake Champlain | -2.3999 | G3P |
| TRINITY_DN225925_c2_g1 | Lake Champlain | -3.3102 | CO9A1,CAS4 |
| TRINITY_DN212063_c8_g1 | Lake Champlain | -2.3827 | |
| TRINITY_DN228356_c29_g1 | Lake Champlain | -2.5795 | |
| TRINITY_DN244049_c11_g1 | Lake Champlain | -1.8143 | TPM1 |
| TRINITY_DN212606_c3_g1 | Lake Champlain | -1.0802 | |
| TRINITY_DN242399_c14_g3 | Lake Champlain | -1.3129 | LDH |
| TRINITY_DN197078_c13_g1 | Lake Champlain | -3.0093 | |
| TRINITY_DN245021_c12_g1 | Lake Champlain | 1.8572 | |
| TRINITY_DN222245_c6_g5 | Lake Champlain | -1.3513 | |
| TRINITY_DN242399_c11_g4 | Lake Champlain | -1.2810 | LDH |
| TRINITY_DN242399_c11_g8 | Lake Champlain | -1.4071 | LDH |
| TRINITY_DN244658_c37_g6 | Lake Champlain | -1.7798 | MYH1 |
| TRINITY_DN209061_c15_g1 | Lake Champlain | 1.4388 | CYTB |
| TRINITY_DN157917_c2_g1 | Lake Champlain | -1.8927 | |
| TRINITY_DN207252_c0_g2 | Lake Champlain | -1.4782 | LRC17 |
| TRINITY_DN224515_c4_g2 | Lake Champlain | -2.0366 | TPM1,TPM3 |
| TRINITY_DN179969_c5_g1 | Lake Champlain | 1.5089 | TRI54,TRI55 |
| TRINITY_DN157982_c0_g1 | Connecticut River | -6.8292 | |
| TRINITY_DN240259_c5_g8 | Connecticut River | -5.4346 | |
| TRINITY_DN217408_c4_g3 | Connecticut River | -10.7690 | |
| TRINITY_DN187825_c15_g1 | Connecticut River | 11.2594 | BGBP |
| TRINITY_DN234143_c2_g1 | Connecticut River | 6.6125 | EF2 |
| TRINITY_DN185982_c4_g2 | Connecticut River | -11.4034 | SAM9L |
| TRINITY_DN159668_c19_g2 | Connecticut River | 4.1247 | |
| TRINITY_DN250469_c0_g1 | Connecticut River | -4.3989 | |
| TRINITY_DN195091_c9_g2 | Connecticut River | 7.2467 | DYRK4 |
| TRINITY_DN188477_c3_g2 | Connecticut River | 8.8424 | |
| TRINITY_DN225176_c4_g1 | Connecticut River | -11.3863 | BI1 |
| TRINITY_DN183100_c6_g5 | Connecticut River | 2.7948 | |
| TRINITY_DN202651_c1_g5 | Connecticut River | -2.4386 | |
| TRINITY_DN172182_c6_g2 | Connecticut River | 3.1113 | |
| TRINITY_DN178227_c7_g1 | Connecticut River | 6.7034 | |
| TRINITY_DN238272_c11_g4 | Connecticut River | 6.7448 | |
| TRINITY_DN213610_c1_g1 | Connecticut River | 2.9498 | ENTK |
| TRINITY_DN206493_c5_g2 | Connecticut River | -3.6856 | |
| TRINITY_DN161631_c1_g1 | Connecticut River | -5.1256 | ADT3 |
| TRINITY_DN138457_c0_g1 | Connecticut River | -8.1517 | |

| | | | |
|---|---|---|---|
| TRINITY_DN221029_c7_g4 | Connecticut River | 7.4917 | |
| TRINITY_DN186605_c16_g1 | Connecticut River | 3.3425 | |
| TRINITY_DN187904_c0_g1 | Connecticut River | 1.7613 | ICEF1,CNIPF |
| TRINITY_DN222899_c3_g4 | Connecticut River | -3.8339 | MYSS,MYH6,MYH7 |
